# Neural correlates of device-based sleep characteristics in adolescents

**DOI:** 10.1101/2024.05.31.596798

**Authors:** Qing Ma, Barbara J Sahakian, Bei Zhang, Zeyu Li, Jin-Tai Yu, Fei Li, Jianfeng Feng, Wei Cheng

## Abstract

Understanding the brain mechanisms underlying objective sleep patterns in adolescents and their implications for psychophysiological development is a complex challenge. Here, we applied sparse canonical correlation (sCCA) analysis on 3300 adolescents from Adolescent Brain Cognitive Development (ABCD) study, integrating extensive device-based sleep characteristics and multimodal imaging data. We revealed two sleep-brain dimensions: one characterized by later being asleep and shorter duration, linked to decreased subcortical-cortical network functional connectivities; the other showed higher heart rate and shorter light sleep duration, associated with lower brain volumes and decreased functional connectivities. Hierarchical clustering based on brain dimension associated with sleep characteristics revealed three biotypes of adolescents, marked by unique sleep profiles: biotype 1 exhibited delayed and shorter sleep, coupled with higher heart rate during sleep; biotype 3 with earlier and longer sleep, accompanied by lower heart rate; and biotype 2 with intermediate pattern. This biotypic differences also extended to cognition, academic attainment, brain structure and function in a gradient order. Longitudinal analysis demonstrated consistent biotypic differences from ages 9 to14, highlighting enduring cognitive and academic advantages in biotype3. The linked sleep-brain dimensions and the associated biotypes were well replicated in a longitudinal sample of 1271 individuals. Collectively, our novel findings delineate a linkage between objective sleep characteristics and developing brain in adolescents, underscoring their significance in cognitive development and academic attainment, which could serve as references for individuals with sleep difficulties and offer insights for optimizing sleep routines to enhance better cognitive development and school achievement.

## Introduction

Sleep is of paramount importance for human survival, brain development and the maintenance of cognitive functions^1^. Adolescence is marked by significant transformations in sleep patterns, including shortened sleep duration, delayed sleep time and shifted circadian rhythm^2^. These predictable changes often coincide with ongoing development of key cognitive and regulatory neural systems that is essential for cognitive development preparing for adult life^1, 3^. Therefore, investigating the relevance between sleep characteristics and the underlying brain development that facilitates cognitive development becomes particularly significant.

Although growing neuroimaging studies suggested a link between specific sleep characteristics and brain structure and function in adolescents^2–4^, most of the sleep measures are self-reported that can be prone to overestimates and inaccuracies^1^. These measures are also often isolated that fail to capture full sleep habits, making it difficult to provide a consistent guideline for research and practical settings^5^. While objective sleep recording like actigraphy receiving increasing attention^6, 7^, existing work involving children and adolescents has been limited by relatively small sample size of a few dozen or, at most, two or three hundred^6, 7^. In addition, the traditional univariate analytical approach in previous studies might impede a comprehensive grasp of the sleep-brain linkage. Recent advancements in machine learning, such as sparse canonical correlation analysis (sCCA), provides a promising approach for systematically exploring associations between two high-dimensional variables, particularly in large datasets. As a multivariate method, sCCA has been successfully applied to healthy populations and individuals with brain disorders^8–11^.

Furthermore, it remains unknown whether the brain, which develops in synchrony with sleep, is able to capture individual developmental differences in the adolescent population. Although individual variations in sleep patterns have been known for a long time from last century^12^, the exploration of such differences has predominantly relied on behavioural sleep indicators. Notably, studies have illuminated a landscape of sleep phenotypes that varied among the elderly^13^ and distinct subtypes among patients with sleep disorders^14, 15^. However, these investigations often lack the integration of objective neurobiological markers, potentially hindering a comprehensive and deeper understanding of overall sleep patterns. This limitation is particularly relevant for adolescents, as this population undergoes significant shifts in sleep patterns^2^. Moreover, electrophysiological studies have revealed individual differences in specific sleep stage, such as non-rapid eye movement, which have been linked to intellectual abilities. These associations may trace back to early childhood development^16^, suggesting potential influences of diverse sleep patterns on adolescent cognitive development. Nonetheless, there is a lack of empirical evidence to substantiate these notions.

In the current study, we sought to investigate the brain structural and functional dimensions associated with various sleep characteristics, as recorded by Fitbit devices. To achieve this, we employed sCCA analysis, using a substantial cohort of adolescents from Adolescent Brain Cognitive Development (ABCD) study at the 2-year follow-up (N = 3300) as discovery sample and the 4-year follow-up dataset (N = 1271) as replication sample. The primary aim was to uncover the dimensions of brain structure and function that are closely associated with specific, interpretable dimensions of sleep. Subsequently, we explored whether the neural dimensions linked with sleep characteristics could reliably capture individual developmental differences in adolescents, including from sleep patterns, to brain structural and functional development, and to cognitive development, as well as academic attainment. (see **Figure 1** for overview of the study design and analyses).

**Figure 1.**
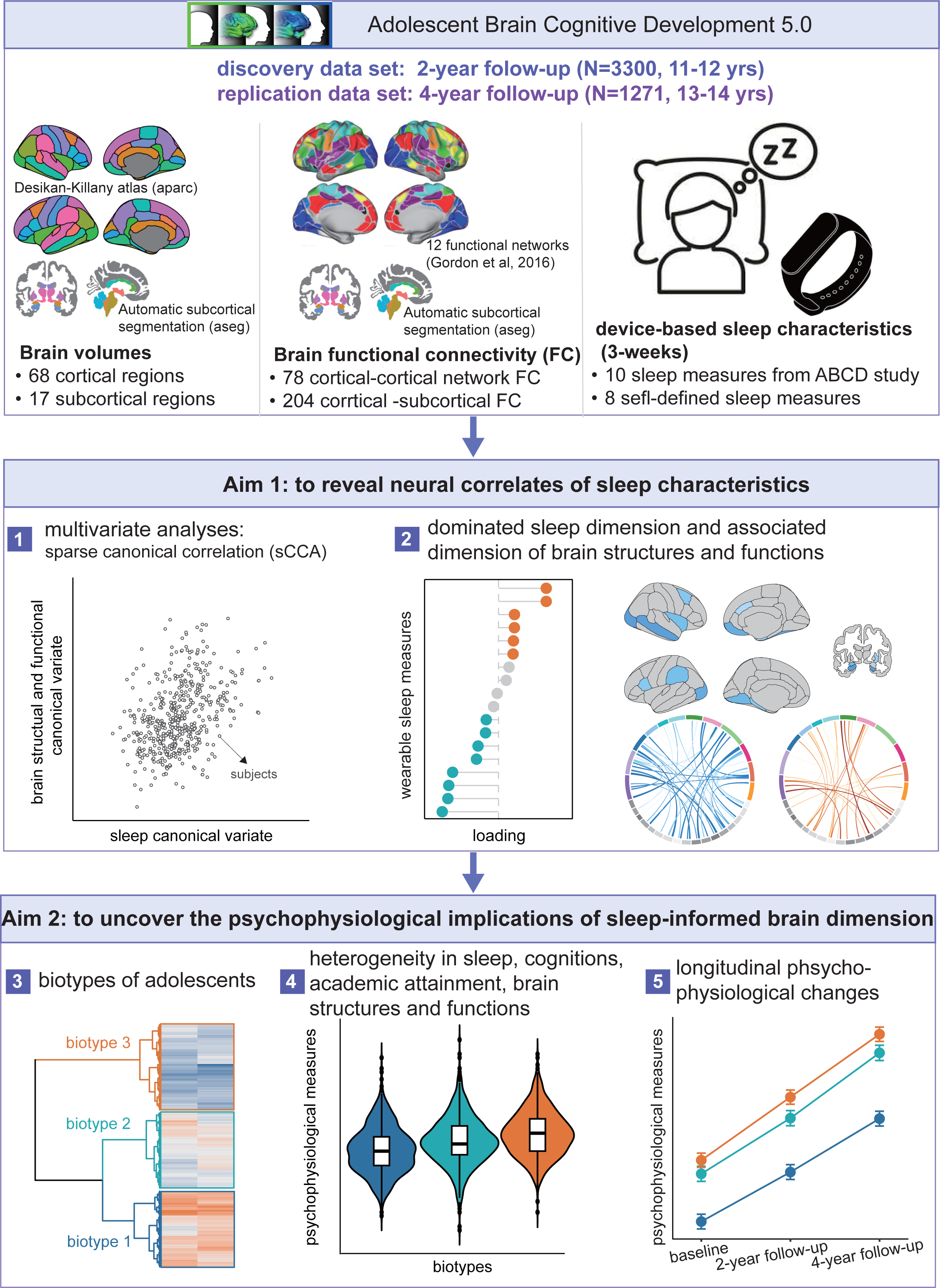
| Overview of the study design and analysis. We used device-base objective sleep characteristics and brain imaging measures, encompassing both brain structure and function, to explore the neural correlates of sleep and their psychological and physiological implications in adolescents. All the data used in this study is sourced from Adolescent Brain Cognitive Development (ABCD) study. Discovery findings were based on the 2-year follow-up data at ages 11-12 (N = 3300), and replication was performed using the 4-year follow-up data (N = 1271).

## Materials and methods

### Participants

A total of 3,300 adolescents with both sleep characteristics and brain imaging scans at 11-12 years old (the 2-year follow-up data) from Adolescent Brain Cognitive Development (ABCD 5.0) study were included as discovery sample in the current study. The replication sample consisted of 1,271 adolescents at the 4-year follow-up. The detailed exclusion criteria for sleep and brain imaging data were provided in Supplementary Methods, **Figure S1** and **S2**. Written and oral informed consent were obtained by the ABCD investigators from parents and adolescents, respectively. More details of the subjects and the collection are provided at the ABCD website (https://abcdstudy.org/scientists/protocols/) and elsewhere^17^.

### Sleep characteristics

Sleep characteristics were objectively collected via Fitbit wristband records over a three-week period, proving weekly summaries based on valid days (i.e. > 180 minutes of sleep). Only participants with at least four records were included. Across the three weeks, an average of 18 sleep indicators were collected daily, encompassing various sleep dimensions such as total sleep duration, duration of wakefulness, duration of light sleep, duration of deep sleep, duration of rapid eye movement (REM) phase, number of awakenings, and averaged heart rate within each of four sleep stages. To provide a more comprehensive insight into sleep patterns, eight additional indicators were calculated based on ABCD measurements and previous studies^13^. These self-defined indicators included four time-point related measures (start time in bed, sleep onset time, wakefulness time, out bed time) and four sleep efficiency related measures (duration between in bed and sleep onset, duration between wakefulness and out of bed, proportion of deep sleep and REM duration to total seep duration). Detailed definitions of these sleep characteristics were provided in Supplementary **Table S1**.

### Brain measures

Magnetic resonance images of brain structure (T1-weighted) and resting-state functional data underwent standardized preprocessing by the ABCD team^18^, and only the data recommended for use after official quality control were included. Brain structure was parcellated into 68 cortical and 40 subcortical regions using Desikan-Killany (DK) Atlas^19^ and Automatic Subcortical Segmentaion (ASEG) Atlas^20^, respectively. Seventeen subcortical regions were selected for analysis after removing non-subcortical areas, including bilateral thalamus, caudate, putamen, globus pallidus, hippocampus, amygdala, accumbens area, ventral diencephalon and the brain stem from the ASEG Atlas. For resting-state functional data, we computed connectivities within and between predefined parcellated networks, excluding misaligned regions^21^. These encompassed 12 networks are auditory, cingulo-opercular, cingulo-parietal, default-mode, dorsal-attention, fronto-parietal, retrosplenial-temporal, salience, sensorimotor-hand, sensorimotor-mouth, ventral-attention, and visual networks. Additionally, connectivities between these networks and 17 subcortical regions from ASEG Atlas were considered. To mitigate confounding variables in canonical correlations, we regressed age and sex from brain imaging and sleep data and the FD head motion was additionally regressed for functional connectivity. After these preprocessing steps, brain imaging and sleep data were normalized to generate z statistics prior to sCCA analysis.

### Cognitive assessment and academic attainment

Cognitive performance was measured using NIH Toolbox, comprising seven tests categorizing crystallized and fluid intelligence. The crystallized intelligence score summarizes scores from (1) the oral reading recognition test and (2) the picture vocabulary test. The fluid intelligence score summarizes scores from (3) the flanker inhibitory control and attention test; (4) the list-sorting working memory test; (5) the dimensional change card sort test; (6) the pattern comparison processing speed test and (7) the picture sequence memory test. Due to missing records, some tests were excluded, resulting in assessments of oral reading recognition, picture vocabulary, flanker inhibitory control and attention, pattern comparison processing speed, picture sequence memory and crystallized intelligent score. Academic attainment was determined through parental evaluation of their children’s school grades, based on the question, “What grades did your child receive in school last year?” (sag_grade_type), during follow-ups 2,3, and4, corresponding to the achievement level of follow-up 1,2, and 3, respectively.

### Sparse canonical correlation analysis

We applied sparse canonical correlation analysis (sCCA) to identify novel linear combinations (canonical variate) of brain imaging measures (brain volume and network functional connectivity) significantly covariant with the combinations of wearable sleep metrics. sCCA identifies optimal canonical coefficient (**u** for brain imaging data, **v** for sleep metrics) that maximize the correlation between two canonical variates. To avoid potential overfitting, we utilized L1 and L2-norm elastic net regularization. The L2 norm parameter were fixed at 1 while those of L1 norm were set by tuning procedure. We tuned the L1-norm parameters for the brain images (including brain volumes and network functional connectivity) and sleep characteristics respectively (Supplementary **Figure S3**), using a grid search (0.1 intervals, range 0 to 1) on 10 randomly chosen two-thirds of the discovery sample. Sparse parameters for optimal canonical coefficients were identified in the first pair of canonical variates, with mean maximal canonical correlations computed across the 10 samples.

The significance of each canonical dimension estimated by correlation between paired canonical variables was assessed via a permutation test, where sCCA analysis was iterated 10,000 times with rows (adolescent individuals) of sleep measures shuffled randomly. Canonical correlation was calculated for each iteration to generate a null distribution. *P*_perm_ value was determined by the count of null correlations surpassing the real average sCCA correlations from the original dataset.

To establish stable canonical loadings (contribution of each measure of sleep and brain imaging on significant canonical dimension), a resampling procedure was conducted with 1,000 iterations, replenishing the remaining one-third from the initial two-thirds selection (akin to bootstrapping with replacement). Canonical loadings with 95% and 99% confidence intervals not spanning zero for sleep and brain imaging measures, respectively, were considered significant and consistent across sampling cohorts. For more detailed sCCA analysis, refer to the work by Xia, Ma ^10^.

### Hierarchical clustering

Subsequently, significant canonical variates derived from brain imaging data were used to cluster adolescents into distinct biotypes. First, we calculated a dissimilarity matrix describing the Euclidean distance between every pair of subjects in the two-dimensional brain feature space. This two-dimensional approach was employed due to the marked correlations observed in the top two pairs of canonical variates. Then, to minimize the total within cluster variance, various linkage methods (average-linkage, single-linkage, complete-linkage, and Ward’s linkage method) were assessed for their agglomerative coefficient. In the current study, Ward’s linkage method, which iteratively link pairs of subjects in closest proximity to form progressively larger clusters in a hierarchical tree, was chosen. The optimal number of clusters was determined by inspecting the dendrogram and evaluating the average silhouette index to consider overall factors.

### Biotypic differences in sleep, cognition, academic attainment, brain and their longitudinal changes

We used one-way analysis of covariance (ANCOVA) to identify sleep characteristics, cognitive performance, academic attainment, and brain imaging measures that differed between relatively homogeneous biotypes. Nuisance covariates, including age, sex, body-mass-index (bmi), puberty, family income, mother and father education level, race and FD motion (only for functional connectivities) were regressed out. The significance level was established at *q* < 0.05 with false discovery rate (FDR) correction across metrics in sleep, cognition, school achievement level, brain volume and brain network functional connectivity respectively. For measures showing significant differences between biotypes, post-hoc analysis of pairwise comparisons were applied by Student’s t-test. Furthermore, we explored the dynamics of these significant biotype differences over time, spanning from baseline to the 2-year and 4-year follow-ups, except for only 1-year, 2-year and 3-year follow-up for academic attainment. Using a two-way analysis of covariance, we detected the potential interaction between biotypes and time, with the same covariates as in the one-way ANCOVA. Given the considerable number of brain imaging measures, we aggregated these measures into four metrics for our longitudinal analysis: averaged cortical volume, subcortical volume, cortico-cortical network functional connectivity, and subcortico-cortical network functional connectivity.

## Results

### Sleep characteristics and brain measures

In current study, we obtained a total of 85 indices of brain volumes (Supplementary **Table S2**), including 68 cortical and 17 subcortical volumes. For brain function, a total of 282 functional connectivity indices were involved, including connectivities within each large-scale network (12), between large-scale networks (66), and between large-scale networks and subcortical regions (204) (**Table S2**). For sleep characteristics, we acquired 18 digital measurements including those provided by ABCD and those in-house calculation (**Table S1**). Supplementary **Table S3** presented demographic details for both discovery and replication samples, and participants selection flowchart was shown in Supplementary **Figure S1** and **Figure S2**.

### Brain-guided dimension of sleep characteristics

sCCA revealed specific patterns, referred to as “canonical variates”, of brain volumes and brain network functional connectivities (FCs) that were associated with specific combinations of sleep measures. Based on the covariance explained (Supplementary **Figure S4**) by each canonical variate, we selected the first two canonical variates for further analysis. Here, we identified two latent sleep-brain dimensions characterized by significant canonical correlates (r = 0.35, *P*_perm_ < 0.001; r = 0.24, *P*_perm_ < 0.001 respectively; 10,000 permutations; **Figure 2A**) that established a clear link between a weighted set of sleep characteristics and a weighted set of brain volumes and functional network connectivities. Then, to better understand the properties of each linked dimension, we implemented a resampling procedure (1,000 bootstrap analysis) to identify sleep and brain measures that were consistently significant across subsets of the data. This procedure demonstrated that the first brain-guided dimension of sleep was “sleep onset-total duration” dominated mode (**Figure 2B**), with more heavily positive weight in being asleep time and negative weight in total sleep duration and number of awakenings. The second dimension was “heart rate-light sleep duration” dominated mode (**Figure 2B**), with the more heavily positive weight in heart rate at all sleep stages and negative weight in light sleep duration.

**Figure 2.**
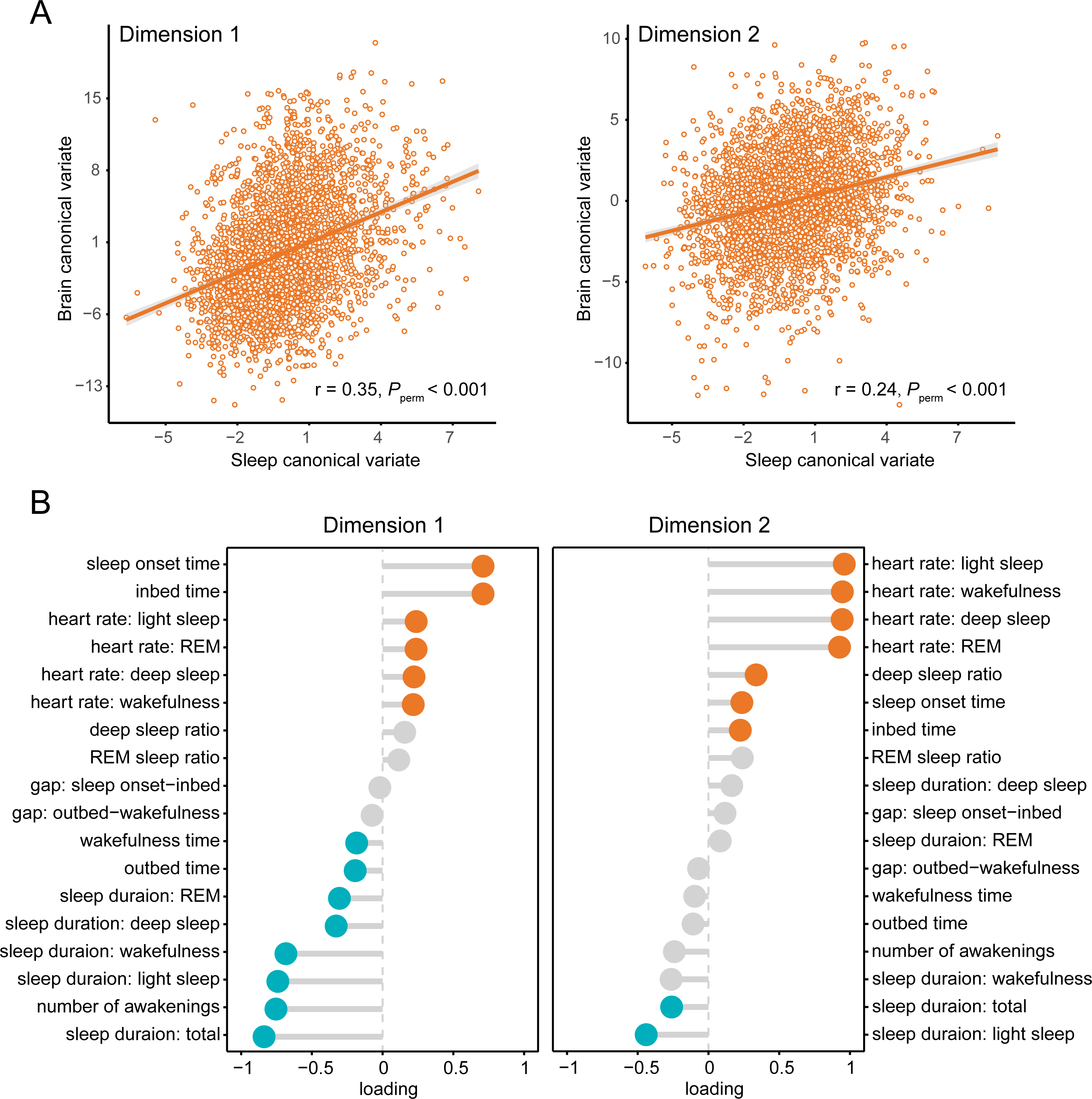
| A. Sparse canonical correlation analysis (sCCA) reveals multivariate patterns of two linked dimensions of sleep and brain. Scatterplots depict the association between brain canonical variate and sleep canonical variate (linear combinations of brain imaging measures, including brain volume and functional network connectivity, and objective sleep characteristics, respectively). The canonical correlations for all two dimensions were statistically significant in discovery data compared to an empirical null distribution (shuffled discovery sample; 10,000 permutations) with *P* < 0.05. **B. Two dimensions of sleep are defined.** Left side indicates “ sleep onset-total duration” dominated mode and right side indicates “heart rate-light sleep duration” dominated mode. Each sleep measure on the Y axis was arranged in descending order according to sleep canonical loadings shown in X axis. The loadings determined to be statistically significant through a resampling procedure (*P* < 0.05, 1,000 bootstrap) are indicated by orange (positive weight) and green (negative weight) colours. Grey circle represents non-significance.

### Sleep-informed dimension of brain measures

Subsequently, we examined the patterns of brain volumes and functional network connectivities underpinning each of the canonical variate. This analysis yielded distinct and unique patterns of brain volumes and functional network connectivities that were related to the two dimensions of sleep (**Figure 2** and Supplementary **Table S4**). As shown in **Figure 3**, it presented the top 20 loadings contributing to the first brain dimension which were linked to first dimension of sleep (“sleep onset-total duration” dominated mode). Specifically, later being asleep and shorter sleep duration were associated with a wide range of decreased functional connectivities between primary and associative cortical networks and subcortical regions, while a relatively small number of increased FCs within and between primary networks. Briefly, the dominate brain functional networks involved were sensorimotor hand network and cingulo-opercular network, while the primarily implicated regions were basal ganglia, amygdala, hippocampus and thalamus. The brain measures showing the strongest negative and positive loads were FC between sensorimotor hand network and pallidum, between sensorimotor hand and sensorimotor mouth network, respectively.

**Figure 3.**
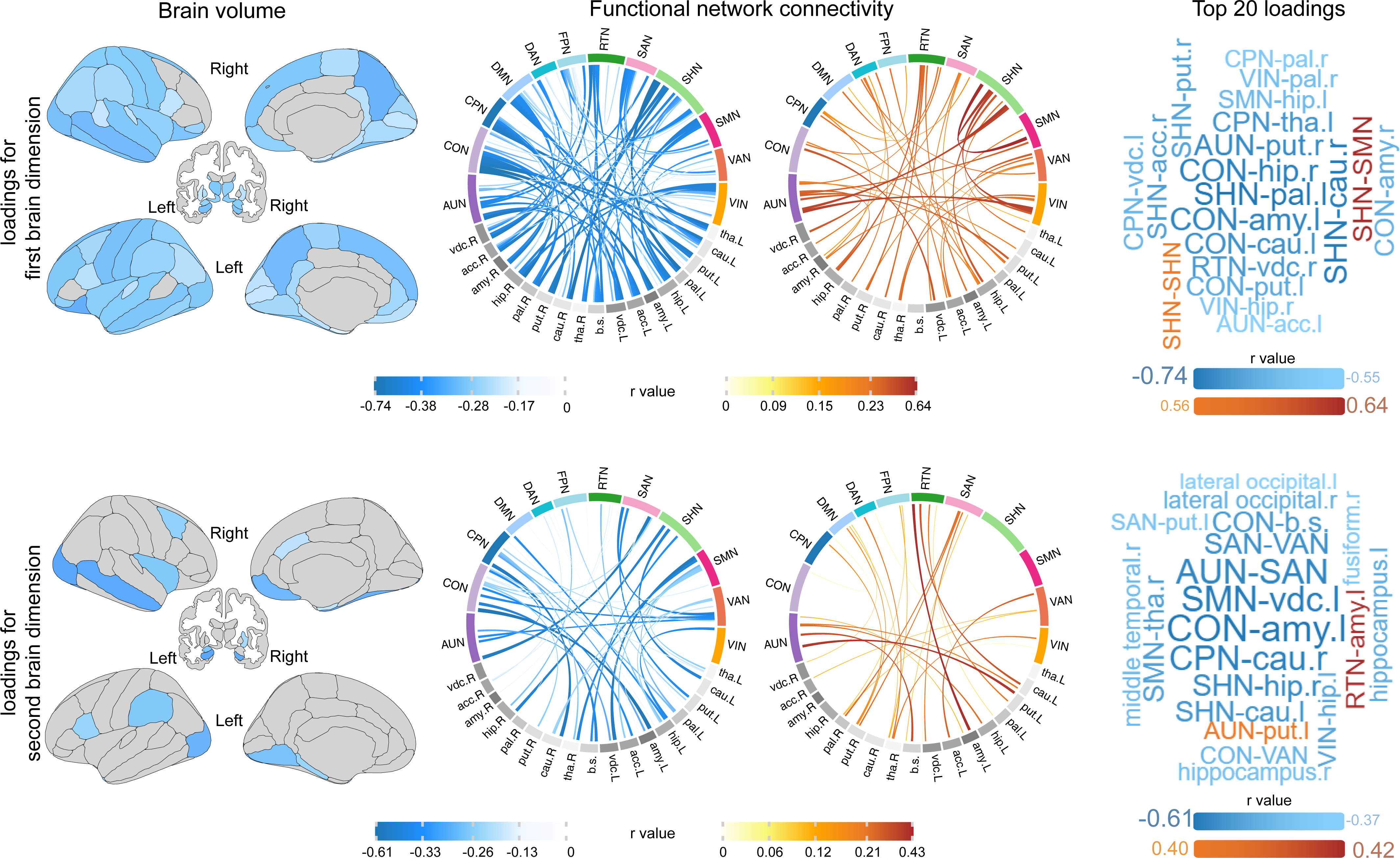
| Patterns of brain volumes and functional network connectivities contribute to linked sleep dimension. Lower brain volumes, decreased and increased connectivities of functional networks are associated with each sleep dimension. For ease of viewing, we present the top 20 most weighted brain imaging canonical loadings in the last column for each dimension. The thickness of the chords represents the loadings of functional connectivity. Only loadings determined to be statistically significant consistent (*P* < 0.05) during 1,000 bootstrap resampling procedure were shown here. AUN, auditory network; CON, cingulo-opercular network; CPN, cingulo-parietal network; DMN, default mode network; DAN, dorsal attention network; FPN, fronto-parietal network; RTN, retrosplenial-temporal network; SHN, somato-hand network; SMN, somato-motor netwok; SAN, salience network; VAN, visual attention network; VIN, visual network; tha., thalamus, cau., caudate; put., putamen; pal., (globus) pallidus; b.s., brain stem; hip., hippocampus; amy., amygdala; acc., accumbens; vdc., ventral diencephalon; L, left (hemisphere); R, right (hemisphere). The visualization tool utilized include the *ggseg* and *circlize* packages in R.

Differently, lower brain volumes, decreased cortical -subcortical FCs, along with FCs between cortical networks were evident in the second (“heart rate-light sleep duration”) dominated mode, which are specifically associated with higher heart rate and shorter sleep duration. The implicated regions showing lower brain volumes were primarily the lateral occipital, hippocampal, middle temporal, and fusiform areas. The decreased FCs were primarily involved functional networks such as sensorimotor hand- and mouth-networks, cingulo-opercular network, cingulo-parietal network and salience network, while the involved regions in the FCs were primarily the amygdala, ventral diencephalon, basal ganglia, hippocampus, thalamus and brain stem. The FC with the highest negative and positive loadings were respectively located between the cingulo-opercular network and the amygdala and between the retrosplenial temporal network and the amygdala.

### Three adolescent biotypes based on dimension of brain measures

Following the identification of two sleep-brain dimensions, we further investigated whether adolescents could be clustered into relatively homogeneous biotypes based on the sleep-informed brain dimensional feature space. Hierarchical clustering along the brain dimension (two canonical variates for each subject) identified an optimal three-cluster solution (**Figure 4A**). Among these clusters, biotype 1 encompassed 1058 individuals, biotype 2 comprised 1049 individuals, and biotype 3 was represented by 1193 individuals. These three biotypic groups demonstrated discernible divergences in terms of their sleep profiles, as elucidated in **Figure 4B**. Biotype 1 deviated far from the average in almost all of the sleep characteristics, showing the worst sleep performance and characterized by the shortest duration in all sleep stages, the latest bedtime and fall sleep onset time, the earliest wake-up time, and the highest heart rate across sleep stages. Conversely, biotype 2 aligned closely with the average level across all sleep characteristics. Biotype 3 demonstrated a sleep profile opposite to that of biotype 1, presenting the best sleep performance. It featured the longest duration across all sleep stages, the earliest bedtime and fall asleep onset time, and the lowest heart rate during all sleep stages. Detailed statistical differences of each sleep characteristic across these biotypes were shown in Supplementary **Table S5**.

**Figure 4.**
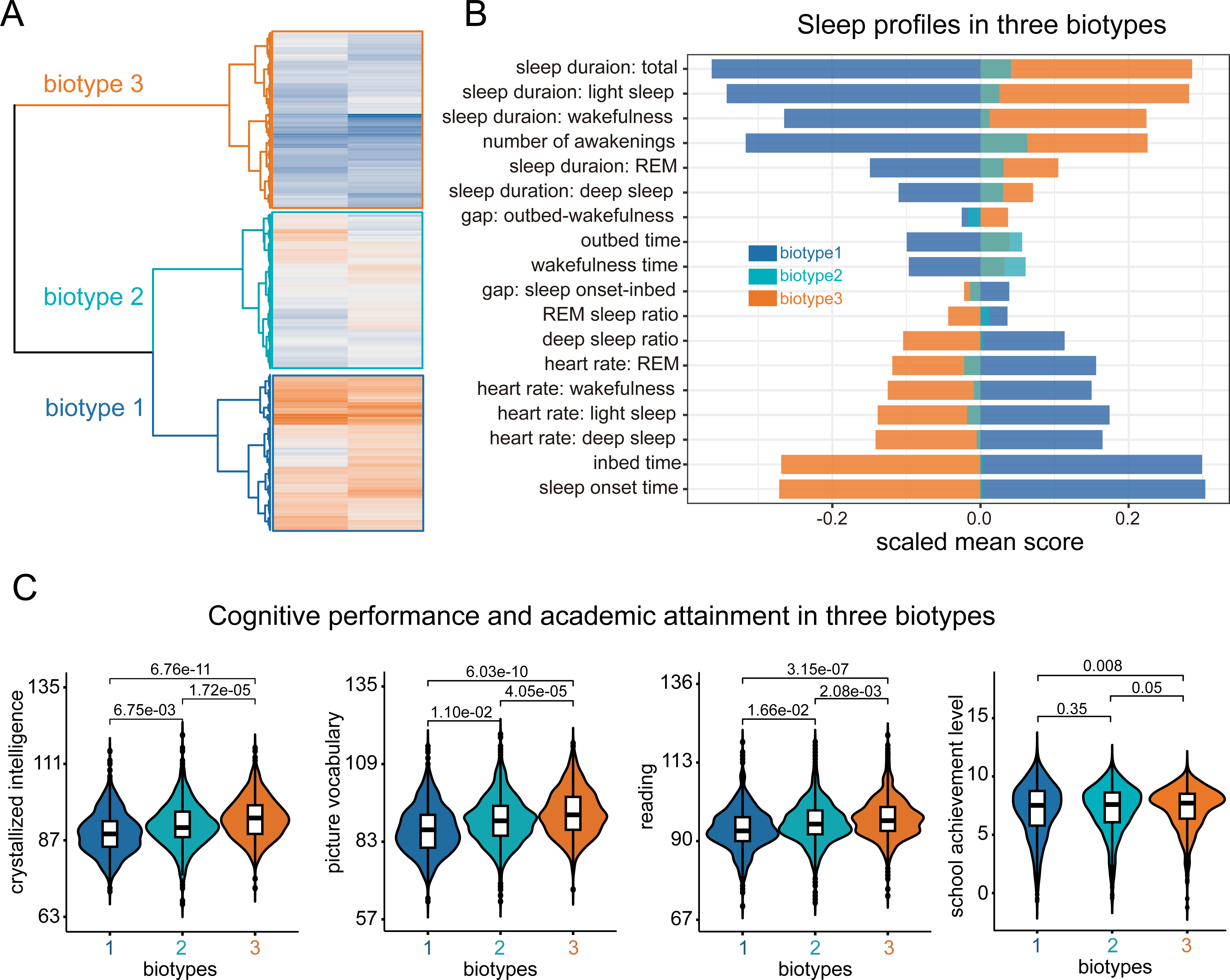
| A. Hierarchical clustering on brain canonical variate reveals three biotypes of adolescents. Heat map and dendrogram depict hierarchical clustering on 3300 individuals in discovery sample (rows) along two dimensions (columns) using Euclidean distance (dendrogram) between two dimensions of brain canonical variates. The height of each linkage in the dendrogram represents the distance between the clusters joined by that link. **B. Sleep patterns in three biotypes.** Averaged z statistics were calculated for each of the three biotypes across all 18 sleep characteristics following the adjustment for age and sex across subjects. Lines and bars coloured blue represent biotype 1 (N = 1058), green represent biotype 2 (N = 1049) and orange represent biotype 3 (N = 1193). **C. Group differences of cognitive performances and academic attainment across three biotypes.** For cognitive domains, only those exhibiting statistically significant differences (as shown in the first three violin plots; *P* < 0.05, FDR corrected) in the main effect of biotypes, as identified by the ANCOVA test, are presented here. The Y-axis represents the uncorrected cognitive performance scores at ages of 11-12, which coincided with the year device-based sleep indicators were recorded. Numbers across any two groups represent raw *P* values. All the significances were adjusted for age, sex, body-mass-index (bmi), puberty, family income, mother and father education level, as well as race.

### Biotype-specific cognitive performance, academic attainment, and brain measures

To further explore the psychophysiological implications of the three biotypes, we tested for differences in cognitive performances, school achievement, brain volumes and functional network connectivities among the three identified groups. Results showed that the three biotypes differed significantly in the cognitive performances (FDR correction at *P* < 0.05), including crystalized intelligence (F = 22.21, *P* <0.001), picture vocabulary (F = 20.04, *P* < 0.001), and oral reading recognition (F = 13.35, *P* <0.001). A closer pairwise examination showed a gradient difference between biotypes in cognition. Specifically, biotype 3 outperformed both biotype 1 and biotype 2, and biotype2 also exhibited better performance than biotype1 in the domains of crystallized intelligence (biotype 2 -biotype 1: t =2.71, *P* = 6.75 × 10^−3^; biotype 3 -biotype 2: t = 4.30, *P* = 1.72 × 10^−5^; biotype 3 -biotype 1: t = 6.55, *P* = 6.76 × 10^−11^), picture vocabulary (t = 2.55, *P* = 1.10 × 10^−2^; t = 4.11, *P* = 4.05 × 10^−5^; t = 6.21, *P* = 6.03 × 10^−10^), and reading recognition ( t = 2.40, *P* = 1.66 × 10^−2^; t = 3.08, *P* = 2.08 × 10^−3^; t = 5.13, *P* = 3.15 × 10^−7^) (**Figure 4C** and Supplementary **Table S6**). Similarly, biotype 3 showed higher school achievement than biotype 1 (t = 2.65, *P* = 0.008) and biotype 2 (t = 1.91, *P* =0.05) (**Figure 4C**).

Three biotypes also exhibited significant differences in brain volumes and brain network functional connectivities (Supplementary **Table S7**). Similar gradient differences were observed in brain volumes, displaying an increase from biotype 1 to biotype 3 (biotype 2 -biotype 1: range of t_range_ = 2.66 −8.98, range of *P*_range_ = 0.008 −4.60 × 10^−19^; biotype 3 -biotype 2: t_range_ = 7.21 −18.94, *P*_range_ = 6.75 × 10^−13^ −5.09 × 10^−76^; biotype 3 -biotype 1: t_range_ = 4.34 −12.51, *P*_range_ = 1.44 × 10^−5^ −4.28 × 10^−35^) (FDR correction at *P* < 0.05; Supplementary **Figure S5A**). In terms of functional connectivities of brain networks, there was also a gradual increase or decrease observed from biotype 1 to biotype3 (all *P* < 0.05, FDR corrected; Supplementary **Figure S5B**).

### Longitudinal changes of cognitive performance, academic attainment, and brain measures of three adolescent biotypes

To gain a deeper insight into implications of these biotypes informed by sleep-brain dimensions, we also examined longitudinal changes of cognitive and brain profiles within the three biotypes over a four-year period spanning from 9-10 years to 13-14 years. Results revealed that all three biotypes exhibited progressive improvements in cognitive ability, specifically in crystallized intelligence, picture vocabulary, and oral reading recognition, over the four-year span (**Figure 5A**). No significant interaction was detected between biotypes and time (all *P* > 0.05, FDR corrected; Supplementary **Table S8**). In other words, this order gradient differences observed at 11-12 years (2-year follow-up) remained consistent both at 9-10 years (baseline) and at 13-14 years (4-year follow-up) (**Figure 5A**). For academic attainment, three biotypes exhibited no significant interaction with time (F = 1.97, *P* = 0.09), with biotype 3 showed consistent highest school achievement across time. Regarding brain volume changes, all three biotypes demonstrated a decrease in averaged cortical brain volume and an increase in averaged subcortical brain volume. Similar to cognitive development, biotype 1 exhibited the smallest volumes, biotype 3 displayed the largest, and biotype 2 fell in between, across all the three time points (**Figure 5B**). Such gradient differences between three biotypes were also evident in the averaged subcortical-cortical network functional connectivity. In contrast, the averaged cortico-cortical network functional connectivity exhibited a significant interaction with time (F = 9.18, *P* < 0.001), where the biotypic differences that appeared at age 9-10 and 11-12 vanished by age 13-14. With respect to individual brain region, nearly all brain regions exhibited no significant interaction in volume. In terms of individual functional connectivity, 57.8% showed significant interactions (Supplementary **Table S9**).

**Figure 5.**
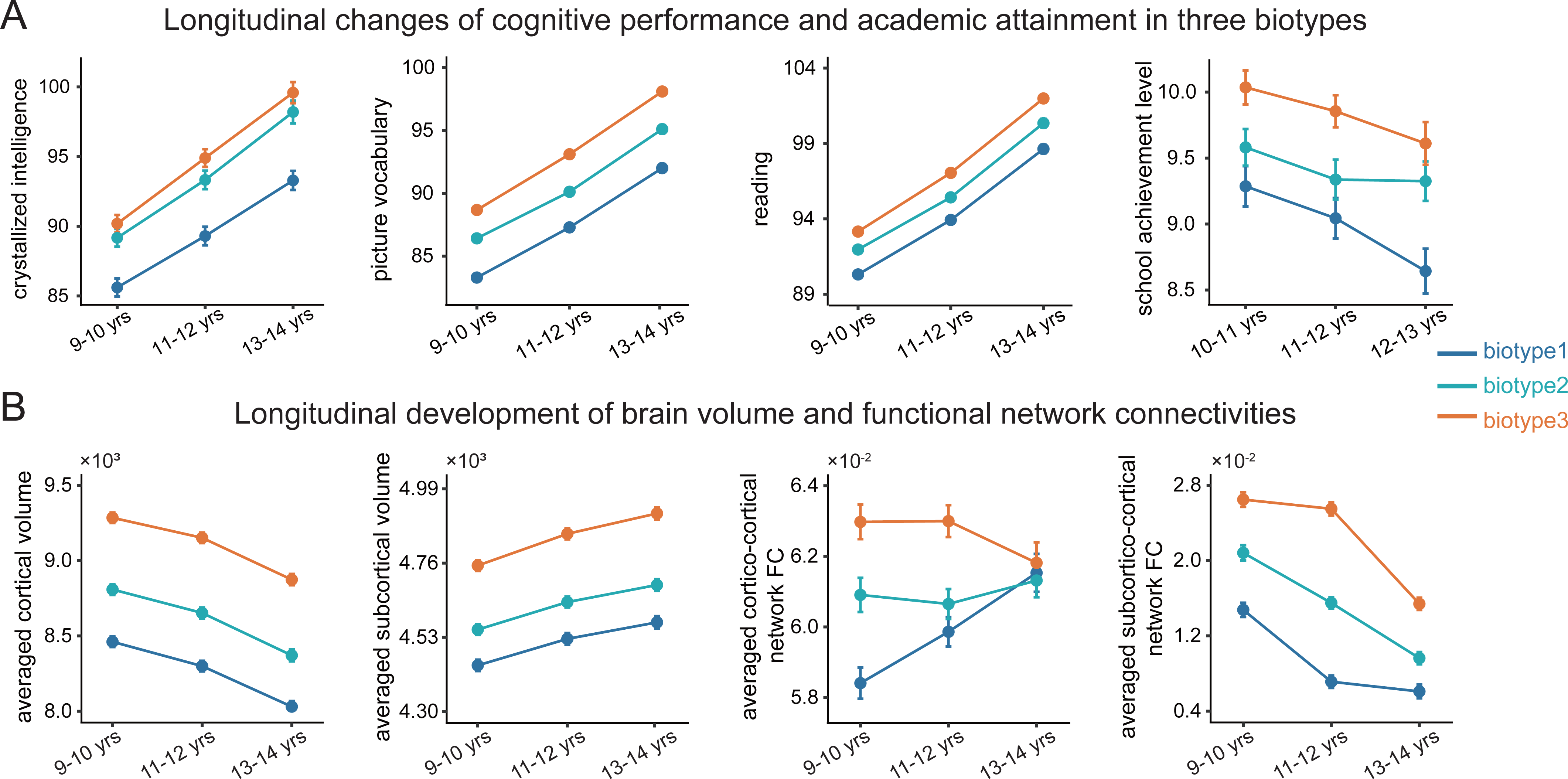
| Longitudinal changes of cognitive performances and academic attainment (A) as well as development of brain volumes and functional network connectivities (B) in three biotypes of adolescents. Only cognitive domains exhibiting statistically significant (*P* < 0.05, FDR corrected) are shown. For academic attainment, due to limited data availability, only the data collected one year before and one year after the two-year follow-up are utilized. Regarding brain measures, only the average values are presented due to the impracticality of displaying an extensive array of measures. The statistics of longitudinal developmental changes for each brain region and each brain functional connectivity are provided in the Supplementary **Table S9**. All statistical analyses were adjusted for age, sex, body-mass-index (bmi), puberty, family income, mother education level, father education level, race and frame-wise head movement (this is only for measures involving functional connectivities).

### Replication of the linked sleep-brain dimensions and associated adolescent biotypes

In this step, we aimed to assess whether the relationship between sleep and brain remained two years later. In the replication sample, we successfully identified two significant latent sleep-brain dimensions (r = 0.29, *P*_perm_ < 0.001; r = 0.21, *P*_perm_ = 0.04 respectively; 10000 permutations; **Figure 6A**). Specifically, sCCA identified two canonical variates in the replication sample that closely resemble the original two linked dimensions of sleep identified in the discovery sample. This is evident from the strong correlations observed between the sleep loadings in the discovery sample and those in the replication sample: r = 0.98 for the first dimension (*P* = 1.82 × 10^−12^) and r = 0.91 for the second dimension (*P* = 2.45 × 10^−7^). The two brain dimensions in the replication sample also exhibited significant correlations of brain loadings with those revealed in the discovery sample (r = 0.98, *P* = 8.86×10^−277^; r = 0.17, *P* = 002). Bootstrap analysis found that the overlapping proportion of brain measures with stable loadings identified in discovery and replication sample were 64% and 27.4%, respectively.

**Figure 6.**
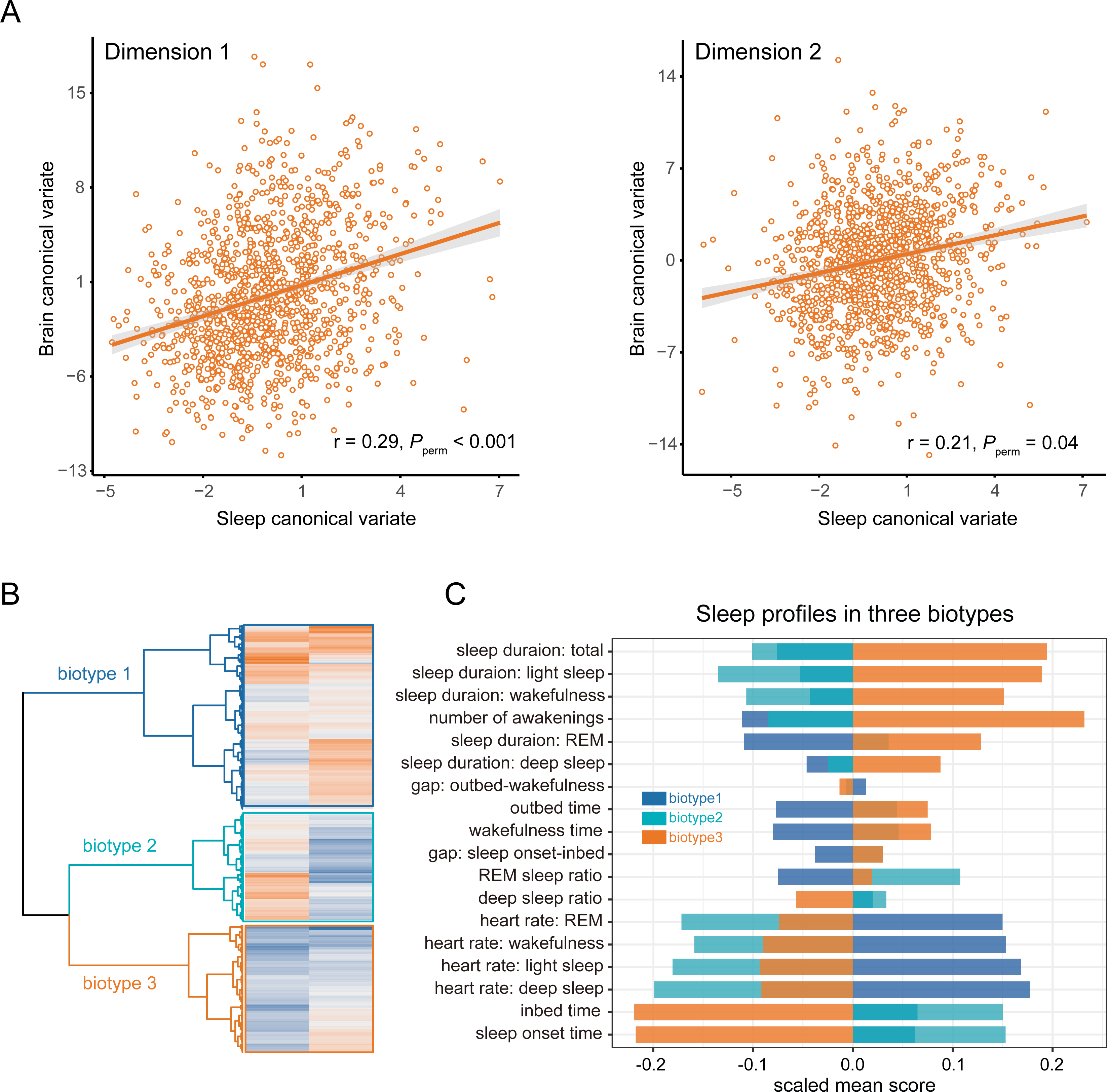
| Sleep-brain dimensions and three sleep profiles are replicated. All the procedures in A and B were repeated in the 4-year follow-up sample of 1271 participants. **A.** The first two dimensions of brain in the replication sample were significantly associated with the first two dimensions of sleep, respectively (*P* < 0.05, 10,000 permutations). **B.** Hierarchical clustering on first two dimensions of brain reveals three biotypes of adolescents with each exhibiting unique sleep profiles.

Hierarchical clustering based on brain dimensions also revealed three distinct biotypes (comprising 561, 323, and 387 individuals, respectively) that exhibited sleep profiles similar to those identified in the discovery sample (**Figure 6B**). Specifically, in the replication sample, biotype 1 was characterized by the shortest sleep duration and the latest sleep onset, whereas biotype 3, in contrast, had the longest sleep duration and the earliest sleep onset. However, the replication of biotype 2 was slightly more complex, with most sleep characteristics aligning closely with the average, except for a higher heart rate. To further validate the findings, we conducted clustering analysis restricted to the first brain dimension due to the weak statistical correlation of the second one. The analysis also revealed three biotypes with highly similar sleep profiles (see Supplement **Figure S6**).

## Discussion

Leveraging a large multimodal imaging data and objective sleep characteristics, coupled with advanced machine learning methodologies, we successfully identified two distinct dimensions of sleep-brain covariance in adolescents. Hierarchical clustering along brain dimension that covary with sleep revealed three adolescent biotypes. These putative biotypes presented gradient differences in sleep profiles, brain volumes and functional connectivities, as well as cognitive performances. Interestingly, all of these psychophysiological differences between three biotypes have already apparent at age of 9-10 and persisted in 13-14 years. The linked sleep-brain dimension and the associated biotypes were well replicable, demonstrating the robustness of the latent dimension of naturalistic sleep and generalizability of the biotypes.

Our evidence suggests that there are two distinct dimensions of sleep characteristics associated with developing brain in the general adolescent population. Our characterization of first dimensional sleep aligns with circadian and physiological changes in adolescent, characterised by sleep loss, restriction and deprivation compared to childhood being plausible^2, 22, 23^. Here, the highly associated characteristics of developing brain were increased connectivities involving sensorimotor network and decreased connetivities between subcortical and cortical networks dominated by sensorimotor network, visual network, cingulo-opercular network and cingulo-parietal network. The sensory systems have been shown to play a vital role in regulating sleep-wake cycles and facilitate neural plasticity essential for learning and memory^24^, with primary sensory cortex, notably the sensorimotor cortex, crucial in sleep physiology^25^. Poor sleep quality, as indicated by questionnaire, also correlates with increased connectivity in the sensory/somatomotor network^26^. During adolescence, increased connectivity in brain regions linked to basic sensory and motor functions is observed, often seen as a “conservative” factor in brain development^27^. Given its complexity, further investigation is needed to determine whether this sleep dimension is “conservative” or “disruptive” to the brain. Moreover, lower integration between subcortical regions and cortical networks underscores the significance of connectivities involving basal ganglia. Basal ganglia has been served as a pivotal regulatory region in control sleep-wake cycle to maintain functions during wakefulness, like motor function, habit formation, and reward/additive behaviors^28^. The maturation of sleep and its regulatory systems spans from infancy to late adolescence^29^, making it unsurprisingly the developmental regulatory system’s sensitivity to sleep patterns. Alteration of basal ganglia structure or function have been found in people with sleep disorders (i.e., obstructive sleep apnea^30^), sleep deprivation^31^, or primary insomnia^32^. The reciprocal relationship between shortened sleep duration and heightened impulsivity has been evidenced in the connectivity between the striatum and cortex^33^.

Compared to the first dimension, the second one is obviously different. With the benefit of objective assessment, we found a significant contribution of heart rate in sleep-brain covariance, a novel insight compared to prior researches. Lower heart rates usually signify better health^34, 35^, whereas higher rates often accompany poor sleep performances like restless sleep, frequent awakenings and excessive daytime sleepiness^36^. Our findings support the long-standing and still popular hypothesis of a heart-brain connection, and their imbalances may lead to negative effect on health^37, 38^. Altered heart rate patterns during sleep have been demonstrated as an objective biomarker of depression^39^ and mild cognitive impairment^40^. Taking together with current study, these, to some extent, imply autonomic functions are relevant to neurodevelopment, prompting further detailed exploration of relationships between heart rate, sleep and brain in developing adolescents. Another difference from first dimension is that brain structure is involved as top contribution in the second sleep-brain dimensional covariation, particularly lower brain volume in memory-related regions like the hippocampus, middle temporal gyrus, fusiform gyrus, and lateral occipital gyrus. Abundant works have supported the benefits of sleep on memory, especially on memory consolidation^41, 42^. Studies have shown that people with poor sleep exhibit a decrease in the volume of the hippocampus in both elderly and adolescents^43, 44^. The hippocampus is known to play a crucial role in transferring new information from temporary to long-term storage in neocortex like the frontal and posterior parietal areas^45, 46^, a process that may entail reactivating neural activity, including sharp wave ripples during slow-wave sleep^47–49^.

Current study confirmed the existence of three distinct biological subtypes of adolescents, as revealed through sleep-informed brain structure and functions, enhancing previous researches on individual differences. Compare to a recent study that utilized large-scale sample from UKB dataset and uncovered different elderly sleep patterns^13^, our research addresses a gap by identifying three distinct sleep profiles among adolescents. Differently, we chose brain structure and function characteristics as classification features, as they offer a greater level of objectivity and represent essential elements compared to behavioral measurements. Interestingly, brain and cognitive differences among the three biotypes, characterized by distinct sleep profiles, exhibited a consistent gradient effect. For instance, the group with the unhealthiest sleep profile (biotype 1) is linked to the lowest cognitive ability, while the one with the healthiest sleep profile (biotype 3) is associated with highest cognitive ability, and the other one was in middle. Moreover, the longitudinal results showed that this gradient difference is very stable for at least two years forward and backward. Similar stable gradient differences were also found in brain structure and function. These stability and differences indicated that sleep-informed brain dimension may serve as a potential biomarker for classification. A recent EEG-based characterization of the brain activity divided the adolescent population into five groups, two of which had cognitive and brain electrical activity that closely resembled the gradient differences we found^50^. Our findings support the close relationship between sleep, brain, and cognition, suggesting the potential to improve adolescent cognitive abilities by adjusting sleep patterns, possibly by falling asleep earlier and getting more sleep, which holds important practical and societal implications.

The strength of the present study includes the utilization of a large longitudinal sample, advanced multivariate methods, objective sleep characteristics and replication. The dimension we identified with brain volumes and functional connectivities not only differentiates adolescents with different psychophysiological performances, but also have the potential to track varied longitudinal trajectories. Nonetheless, several limitations should be noted. First, although we utilized various sleep measures from the ABCD study along with self-defined indicators, it’s important to note that not all sleep measures were covered (i.e., sleep regularity) to comprehensively depict an individual sleep pattern, indicating the potential for incorporating more sleep-related variables in future research. Second, although our current analysis considered objective sleep characteristics and incorporated as much multimodal imaging data as possible, future researches could expand by incorporating more dimensional information for example genomics, to explore the genetic determines of biotypes, to achieve a comprehensive and deep understanding of adolescent sleep.

### Summary

In summary, our findings offer reproducible evidence of two distinct dimensions, with each evident by specific regional and circuit-level patterns characterized by brain’s structural and functional network connectivities that is closely linked to objective sleep characteristics. These results shed light on the brain mechanisms that underlie multidimensional sleep characteristics. Importantly, our study revealed the existence of three distinct biotypes within the general adolescent population, delineated by sleep-informed brain dimensions. These putative biotypes exhibit notable and enduring brain and cognitive variations, offering potential utility in clinical risk prediction for cognitive development throughout adolescence.

## Acknowledgments

We thank Adolescent Brain Cognitive Development (ABCD) Study (https://abcdstudy.org), held in the NIMH Data Archive (NDA), from which we obtained data used in the preparation of this article. This is a multisite, longitudinal study designed to recruit more than 10,000 children age 9–10 and follow them over 10 years into early adulthood. The ABCD Study is supported by the National Institutes of Health and additional federal partners under award numbers U01DA041048, U01DA050989, U01DA051016, U01DA041022, U01DA051018, U01DA051037, U01DA050987, U01DA041174, U01DA041106, U01DA041117, U01DA041028, U01DA041134, U01DA050988, U01DA051039, U01DA041156, U01DA041025, U01DA041120, U01DA051038, U01DA041148, U01DA041093, U01DA041089, U24DA041123, U24DA041147. A full list of supporters is available at https://abcdstudy.org/federal-partners.html. A listing of participating sites and a complete listing of the study investigators can be found at https://abcdstudy.org/consortium_members/. ABCD consortium investigators designed and implemented that study and/or provided data but did not participate in the analysis for or writing of this report. This manuscript reflects the views of the authors and may not reflect the opinions or views of the NIH or ABCD consortium investigators. Wei Cheng was funded by the National Natural Science Foundation of China (82071997), National Key Research and Development Program of China(2023YFC3605400), and the Shanghai Rising-Star Program (21QA1408700). Qing Ma was supported by grants from National Postdoctoral Foundation of China (No. 2021M690700) and Shanghai Postdoctoral Excellence Program (No. 2020045).

## References

1. Leong RLF, Chee MWL. Understanding the Need for Sleep to Improve Cognition. Annual Review of Psychology 2023; 74(1): 27–57.

2. Galván A. The Need for Sleep in the Adolescent Brain. Trends Cogn Sci 2020; 24(1): 79–89.

3. Anastasiades PG, de Vivo L, Bellesi M, Jones MW. Adolescent sleep and the foundations of prefrontal cortical development and dysfunction. Prog Neurobiol 2022; 218: 102338.

4. Yang FN, Xie W, Wang Z. Effects of sleep duration on neurocognitive development in early adolescents in the USA: a propensity score matched, longitudinal, observational study. The Lancet Child & adolescent health 2022; 6(10): 705–712.

5. Buysse DJ. Sleep health: can we define it? Does it matter? Sleep 2014; 37(1): 9–17.

6. Jalbrzikowski M, Hayes RA, Scully KE, Franzen PL, Hasler BP, Siegle GJ et al. Associations between brain structure and sleep patterns across adolescent development. Sleep 2021; 44(10).

7. Tashjian SM, Goldenberg D, Monti MM, Galván A. Sleep quality and adolescent default mode network connectivity. Social cognitive and affective neuroscience 2018; 13(3): 290–299.

8. Drysdale AT, Grosenick L, Downar J, Dunlop K, Mansouri F, Meng Y et al. Resting-state connectivity biomarkers define neurophysiological subtypes of depression. Nature medicine 2017; 23(1): 28–38.

9. Buch AM, Vértes PE, Seidlitz J, Kim SH, Grosenick L, Liston C. Molecular and network-level mechanisms explaining individual differences in autism spectrum disorder. Nature Neuroscience 2023.

10. Xia CH, Ma Z, Ciric R, Gu S, Betzel RF, Kaczkurkin AN et al. Linked dimensions of psychopathology and connectivity in functional brain networks. Nature Communications 2018; 9(1): 3003.

11. Smith SM, Nichols TE, Vidaurre D, Winkler AM, Behrens TEJ, Glasser MF et al. A positive-negative mode of population covariation links brain connectivity, demographics and behavior. Nature Neuroscience 2015; 18(11): 1565–1567.

12. Davies DR. Individual differences in sleep patterns. Postgrad Med J 1976; 52(603): 10–14.

13. Katori M, Shi S, Ode KL, Tomita Y, Ueda HR. The 103,200-arm acceleration dataset in the UK Biobank revealed a landscape of human sleep phenotypes. Proc Natl Acad Sci U S A 2022; 119(12): e2116729119.

14. Blanken TF, Benjamins JS, Borsboom D, Vermunt JK, Paquola C, Ramautar J et al. Insomnia disorder subtypes derived from life history and traits of affect and personality. The lancet Psychiatry 2019; 6(2): 151–163.

15. Romero-Peralta S, García-Rio F, Resano Barrio P, Viejo-Ayuso E, Izquierdo JL, Sabroso R et al. Defining the Heterogeneity of Sleep Apnea Syndrome: A Cluster Analysis With Implications for Patient Management. Arch Bronconeumol 2022; 58(2): 125–134.

16. Smith D, Fang Z, Thompson K, Fogel S. Sleep and individual differences in intellectual abilities. Current Opinion in Behavioral Sciences 2020; 33: 126–131.

17. Garavan H, Bartsch H, Conway K, Decastro A, Goldstein RZ, Heeringa S et al. Recruiting the ABCD sample: Design considerations and procedures. Dev Cogn Neurosci 2018; 32: 16–22.

18. Hagler DJ, Jr., Hatton S, Cornejo MD, Makowski C, Fair DA, Dick AS et al. Image processing and analysis methods for the Adolescent Brain Cognitive Development Study. Neuroimage 2019; 202: 116091.

19. Desikan RS, Ségonne F, Fischl B, Quinn BT, Dickerson BC, Blacker D et al. An automated labeling system for subdividing the human cerebral cortex on MRI scans into gyral based regions of interest. Neuroimage 2006; 31(3): 968–980.

20. Fischl B, Salat DH, Busa E, Albert M, Dieterich M, Haselgrove C et al. Whole brain segmentation: automated labeling of neuroanatomical structures in the human brain. Neuron 2002; 33(3): 341–355.

21. Gordon EM, Laumann TO, Adeyemo B, Huckins JF, Kelley WM, Petersen SE. Generation and Evaluation of a Cortical Area Parcellation from Resting-State Correlations. Cereb Cortex 2016; 26(1): 288–303.

22. Colrain IM, Baker FC. Changes in sleep as a function of adolescent development. Neuropsychol Rev 2011; 21(1): 5–21.

23. Vandendriessche A, Verloigne M, Boets L, Joriskes J, DeSmet A, Dhondt K et al. Psychosocial factors related to sleep in adolescents and their willingness to participate in the development of a healthy sleep intervention: a focus group study. BMC Public Health 2022; 22(1): 1876.

24. Blumberg MS, Dooley JC, Tiriac A. Sleep, plasticity, and sensory neurodevelopment. Neuron 2022; 110(20): 3230–3242.

25. Fasiello E, Gorgoni M, Scarpelli S, Alfonsi V, Ferini Strambi L, De Gennaro L. Functional connectivity changes in insomnia disorder: A systematic review. Sleep Med Rev 2022; 61: 101569.

26. Bai Y, Tan J, Liu X, Cui X, Li D, Yin H. Resting-state functional connectivity of the sensory/somatomotor network associated with sleep quality: evidence from 202 young male samples. Brain Imaging Behav 2022; 16(4): 1832–1841.

27. Váša F, Romero-Garcia R, Kitzbichler MG, Seidlitz J, Whitaker KJ, Vaghi MM et al. Conservative and disruptive modes of adolescent change in human brain functional connectivity. Proceedings of the National Academy of Sciences 2020; 117(6): 3248–3253.

28. Lazarus M, Chen JF, Urade Y, Huang ZL. Role of the basal ganglia in the control of sleep and wakefulness. Curr Opin Neurobiol 2013; 23(5): 780–785.

29. Roffwarg HP, Muzio JN, Dement WC. Ontogenetic development of the human sleep-dream cycle. Science 1966; 152(3722): 604–619.

30. Santarnecchi E, Sicilia I, Richiardi J, Vatti G, Polizzotto NR, Marino D et al. Altered cortical and subcortical local coherence in obstructive sleep apnea: a functional magnetic resonance imaging study. Journal of sleep research 2013; 22(3): 337–347.

31. Maski KP, Kothare SV. Sleep deprivation and neurobehavioral functioning in children. International journal of psychophysiology : official journal of the International Organization of Psychophysiology 2013; 89(2): 259–264.

32. Wu Y, Liu M, Zeng S, Ma X, Yan J, Lin C et al. Abnormal Topology of the Structural Connectome in the Limbic Cortico-Basal-Ganglia Circuit and Default-Mode Network Among Primary Insomnia Patients. Front Neurosci 2018; 12: 860.

33. Yang FN, Liu TT, Wang Z. Corticostriatal connectivity mediates the reciprocal relationship between sleep and impulsivity in early adolescents. medRxiv 2022: 2022.2011.2007.22282025.

34. Zhang D, Wang W, Li F. Association between resting heart rate and coronary artery disease, stroke, sudden death and noncardiovascular diseases: a meta-analysis. Cmaj 2016; 188(15): E384–e392.

35. Olshansky B, Ricci F, Fedorowski A. Importance of resting heart rate. Trends in Cardiovascular Medicine 2022.

36. Penzel T, Kantelhardt JW, Lo C-C, Voigt K, Vogelmeier C. Dynamics of Heart Rate and Sleep Stages in Normals and Patients with Sleep Apnea. Neuropsychopharmacology 2003; 28(1): S48–S53.

37. Silvani A, Calandra-Buonaura G, Dampney RA, Cortelli P. Brain-heart interactions: physiology and clinical implications. Philosophical transactions Series A, Mathematical, physical, and engineering sciences 2016; 374(2067).

38. Zhao B, Li T, Fan Z, Yang Y, Shu J, Yang X et al. Heart-brain connections: Phenotypic and genetic insights from magnetic resonance images. Science 2023; 380(6648): abn6598.

39. Saad M, Ray LB, Bujaki B, Parvaresh A, Palamarchuk I, De Koninck J et al. Using heart rate profiles during sleep as a biomarker of depression. BMC psychiatry 2019; 19(1): 168.

40. Kong SDX, Hoyos CM, Phillips CL, McKinnon AC, Lin P, Duffy SL et al. Altered heart rate variability during sleep in mild cognitive impairment. Sleep 2021; 44(4).

41. Klinzing JG, Niethard N, Born J. Mechanisms of systems memory consolidation during sleep. Nat Neurosci 2019; 22(10): 1598–1610.

42. Rasch B, Born J. About sleep’s role in memory. Physiological reviews 2013; 93(2): 681–766.

43. Liu C, Lee SH, Hernandez_-_Cardenache R, Loewenstein D, Kather J, Alperin N. Poor sleep is associated with small hippocampal subfields in cognitively normal elderly individuals. Journal of sleep research 2021; 30(5): e13362.

44. Lapidaire W, Urrila AS, Artiges E, Miranda R, Vulser H, Bezivin-Frere P et al. Irregular sleep habits, regional grey matter volumes, and psychological functioning in adolescents. PLoS One 2021; 16(2): e0243720.

45. Diekelmann S, Born J. The memory function of sleep. Nat Rev Neurosci 2010; 11(2): 114–126.

46. Himmer L, Schönauer M, Heib DPJ, Schabus M, Gais S. Rehearsal initiates systems memory consolidation, sleep makes it last. Sci Adv 2019; 5(4): eaav1695.

47. Euston DR, Tatsuno M, McNaughton BL. Fast-forward playback of recent memory sequences in prefrontal cortex during sleep. Science 2007; 318(5853): 1147–1150.

48. Ji D, Wilson MA. Coordinated memory replay in the visual cortex and hippocampus during sleep. Nat Neurosci 2007; 10(1): 100–107.

49. Nitzan N, Swanson R, Schmitz D, Buzsáki G. Brain-wide interactions during hippocampal sharp wave ripples. Proc Natl Acad Sci U S A 2022; 119(20): e2200931119.

50. Forbes O, Schwenn PE, Wu PP-Y, Santos-Fernandez E, Xie H-B, Lagopoulos J et al. EEG-based clusters differentiate psychological distress, sleep quality and cognitive function in adolescents. Biological psychology 2022; 173: 108403.

