## Supplementary Materials for "Neural correlates of device-based sleep characteristics in adolescents"

**Short title: Device-based sleep and adolescent brain**

**Materials and methods**

**Study design and participants**

Brain imaging and wearable sleep characteristics were acquired from Adolescent Brain Cognitive Development (ABCD) study, a large ongoing nationwide study conducted across 21 research sites in the United States and aimed at tracking the developmental trajectory of adolescents from the ages of 9 to 10 onwards. Detailed protocols and design of the ABCD study has been described previously(1). The data used in the current study were obtained from the ABCD data release 5.0, with a specific focus on the 2-year follow-up data (11-12 years old), during which both sleep characteristics and imaging scans were available (see Supplementary Fig. S1 for the participant selection flowchart). Initially, 5,017 adolescents underwent sleep measures. Subsequently, 487 participants were excluded due to inadequate wearable recording spanning fewer than four days. For brain imaging scans, 8,092 adolescents possessed structural volume data and 7,912 had functional network connectivity data. Following the rigorous quality control (QC) of images recommended by the ABCD authority team, 400 participants were excluded from the brain volumetric datasets and none from the functional network connectivity datasets. Furthermore, 712 adolescents with a mean frame-wise displacement (FD) of head movement greater than 0.5 mm(2), as well as 648 adolescents lacking either QC records or the FD motion values, were excluded from functional network connectivity data. Finally, a total of 3,300 adolescents with both sleep characteristics and brain imaging scans were included in the current study. The replication sample consisted of 1,271 adolescents at the 4-year follow-up. The exclusion criteria mirrored those used in the discovery sample (Supplementary Fig. S2). Written and oral informed consent were obtained by the ABCD investigators from parents and adolescents, respectively. More details of the subjects and the collection are provided at the ABCD website (<https://abcdstudy.org/scientists/protocols/>) and elsewhere(3).

**References:**

1.	Saragosa-Harris NM, Chaku N, MacSweeney N, Guazzelli Williamson V, Scheuplein M, Feola B, et al. (2022): A practical guide for researchers and reviewers using the ABCD Study and other large longitudinal datasets. *Dev Cogn Neurosci*. 55:101115.

2.	Power JD, Mitra A, Laumann TO, Snyder AZ, Schlaggar BL, Petersen SE (2014): Methods to detect, characterize, and remove motion artifact in resting state fMRI. *Neuroimage*. 84:320-341.

3.	Garavan H, Bartsch H, Conway K, Decastro A, Goldstein RZ, Heeringa S, et al. (2018): Recruiting the ABCD sample: Design considerations and procedures. *Dev Cogn Neurosci*. 32:16-22.

**Supplementary Figures & Tables**

### **Figure S1. Participants selection process for discovery sample**

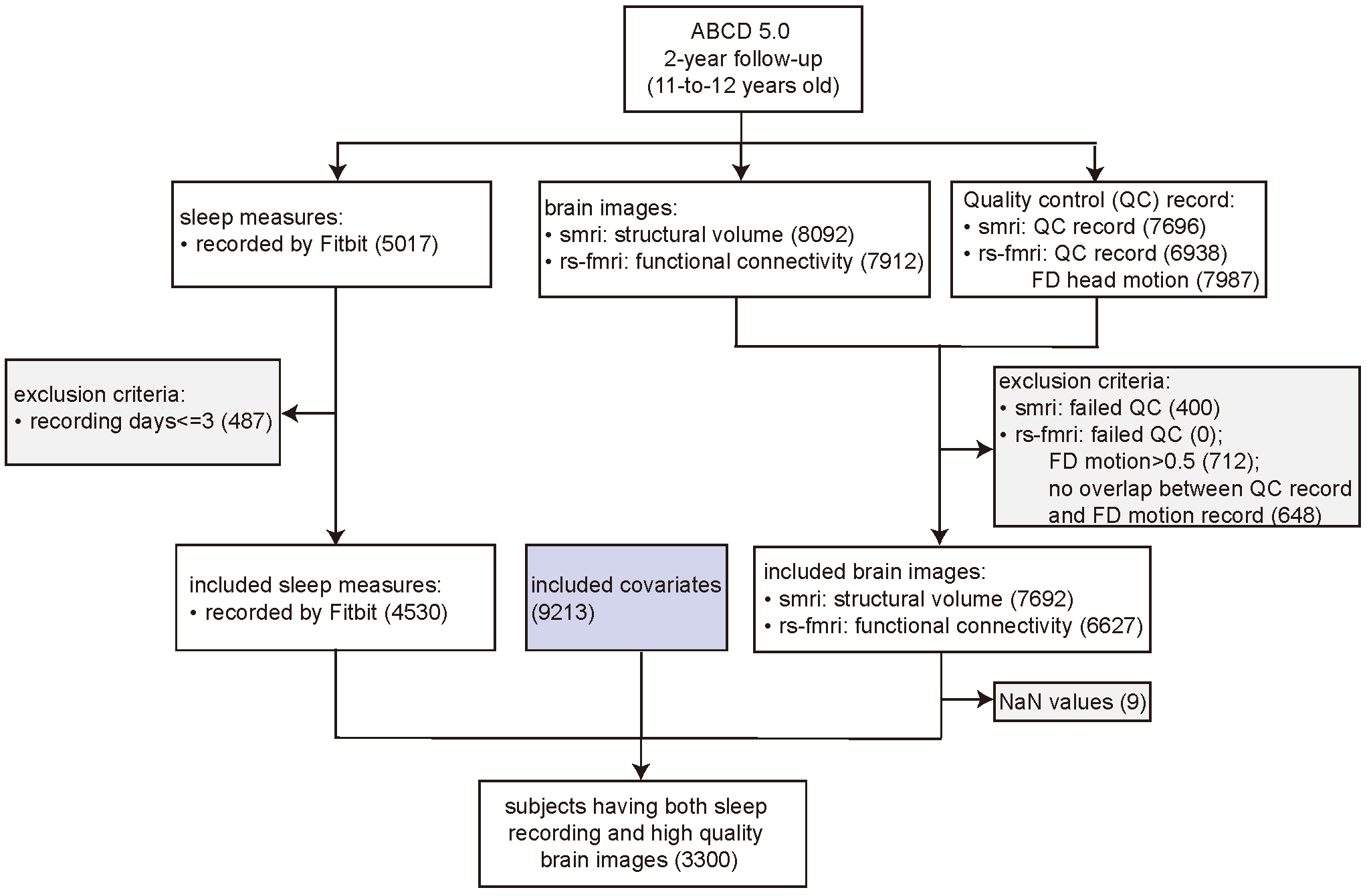

### **Figure S2. Participants selection process for replication sample**

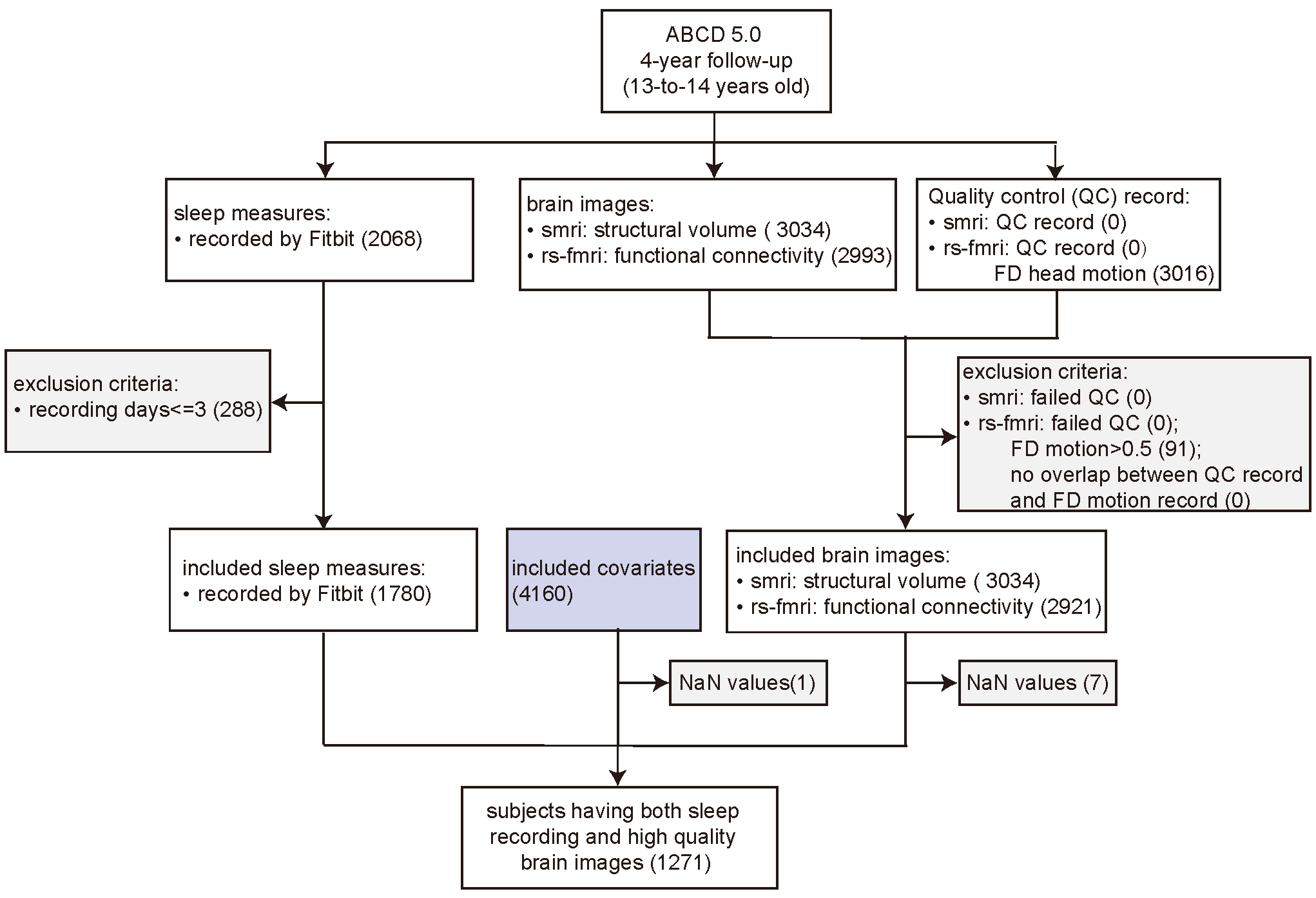

### **Figure S3. Grid search for optimal penalty parameters**

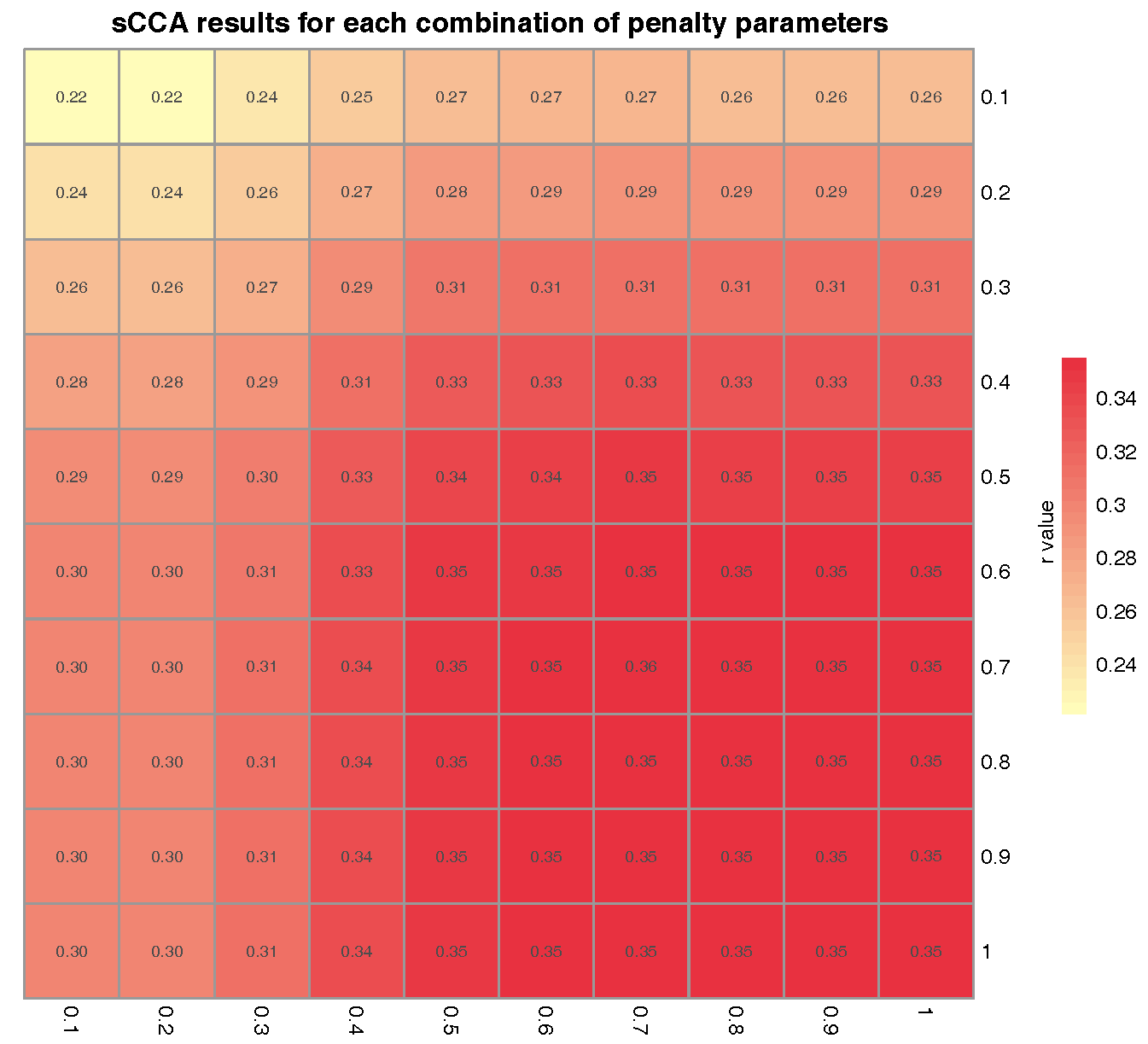

### **Figure S4. Covariance explained at 2-year follow-up**

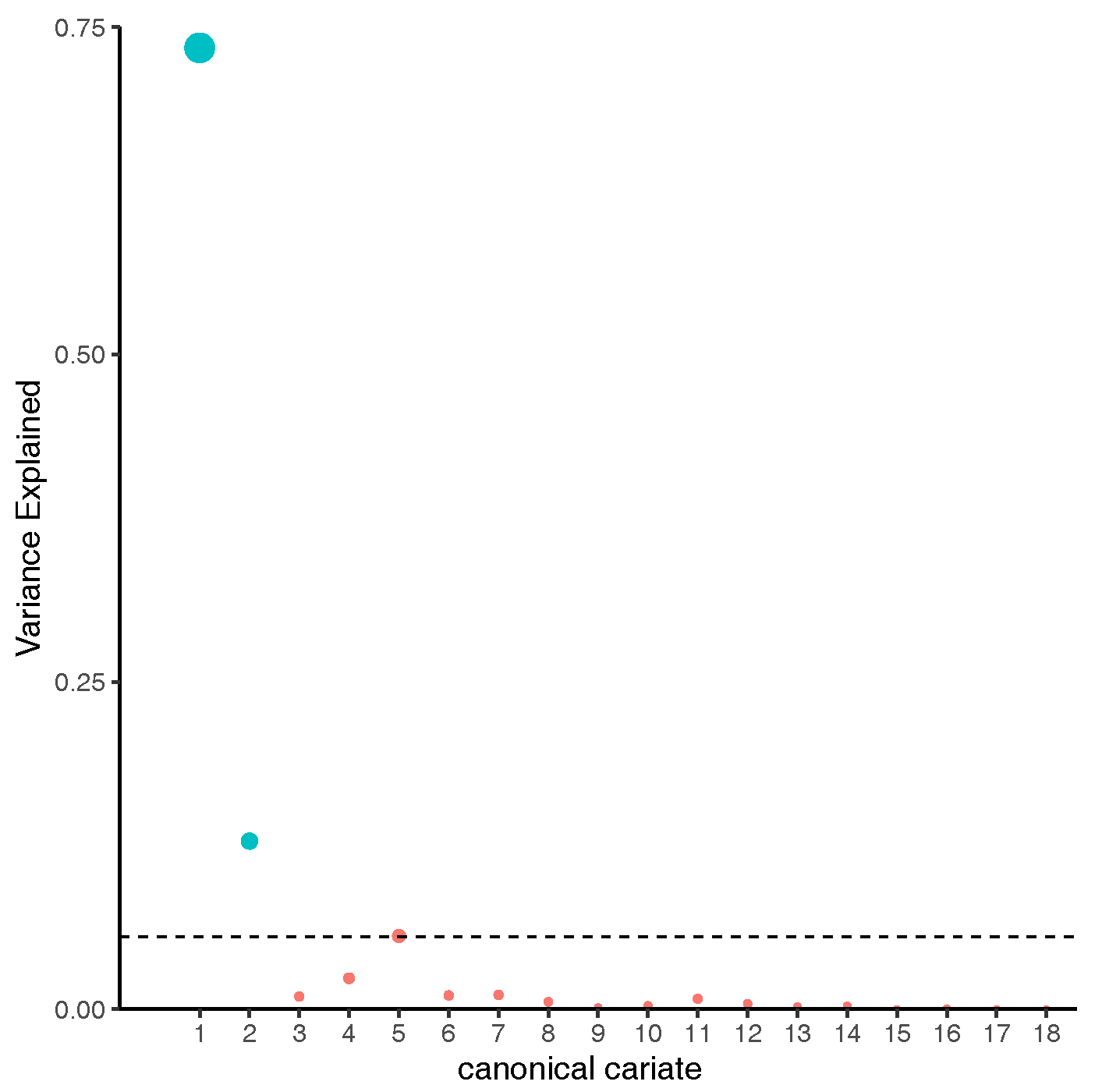

**Figure S5. Pair-wise comparisons of brain volumes and functional network connectivities across three biotypes.**

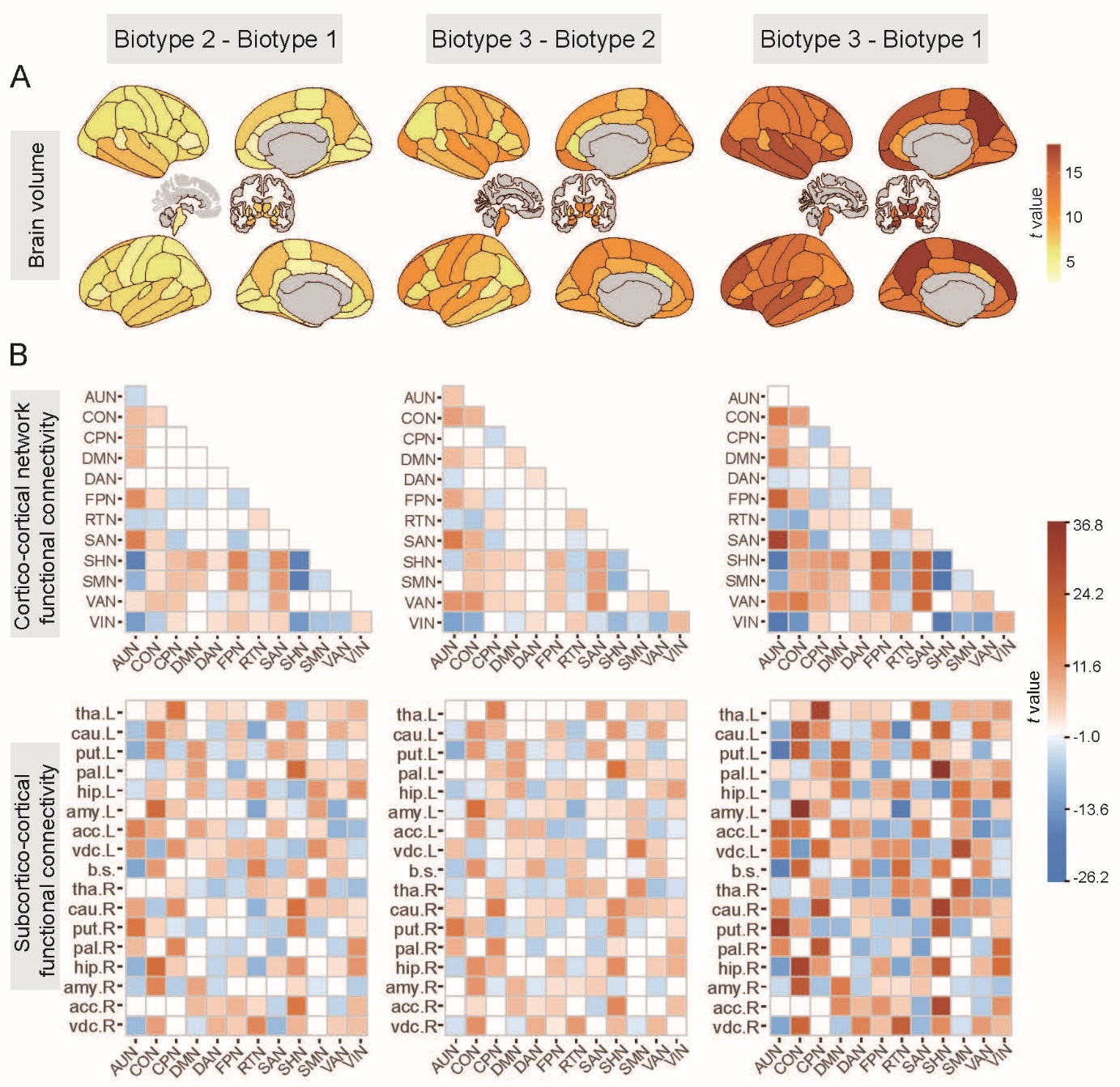

Functional connectivities are categorized into within and between network connectivities, as well as connectivities between networks and subcortical areas. They are respectively displayed at the second and third rows. Measures determined to be statistically significant (*P* < 0.05, FDR corrected) are shown here, adjusting for age, sex, body-mass-index (bmi), puberty, family income, mother and father education level, as well as race. For functional connectivity, additional adjustment for frame-wise head movement was applied.

### **Figure S6. Three sleep profiles are replicated by the first dimension of brain**

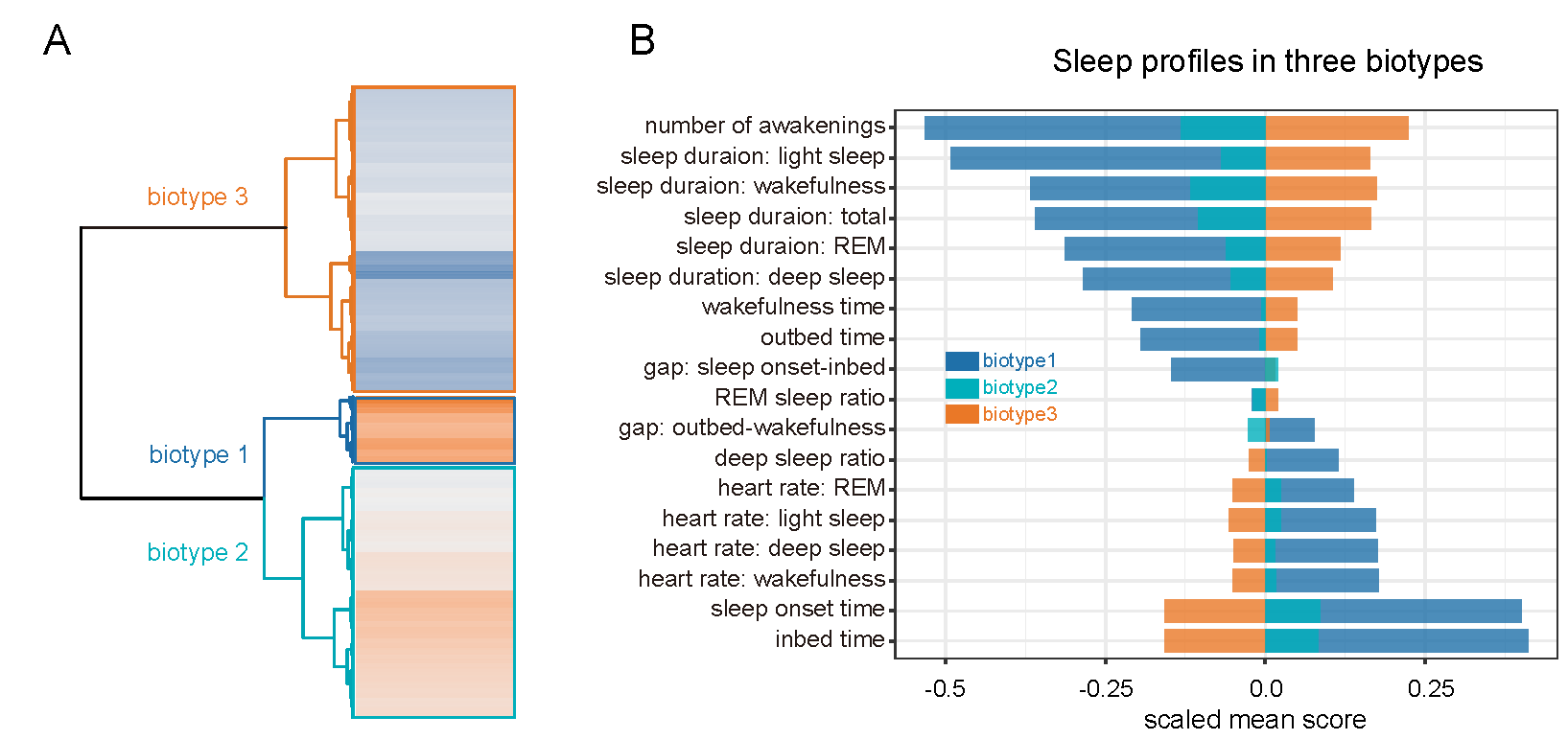

### **Table S1. Sleep characteristics and definitions**

| **ID** | **Sleep features** | **Definition** |
| --- | --- | --- |
| 1 | sleep duration: total | total duration of sleep per night, expressed in minutes |
| 2 | sleep duration: wakefulness | total duration of being wake during sleep, expressed in minutes |
| 3 | sleep duration: light sleep | total duration of light sleep during sleep, expressed in minutes |
| 4 | sleep duration: deep sleep | total duration of deep sleep during sleep. expressed in minutes |
| 5 | sleep duration: REM | total duration of the rapid eye movement (REM) phase of sleep, expressed in minutes |
| 6 | number of awakenings | count of be awake during sleep, expressed in numbers |
| 7 | heart rate: wakefulness | average heart rate during wake phase of sleep, expressed in beats per minute |
| 8 | heart rate: light sleep | average heart rate during light sleep, expressed in beats per minute |
| 9 | heart rate: deep sleep | average heart rate during deep sleep, expressed in beats per minute |
| 10 | heart rate: REM | average heart rate during the rapid eye movement phase of sleep, expressed in beats per minute |
| 11 | inbed time | start time in bed, expressed in hours since midnight of the previous day |
| 12 | sleep onset time | start time of primary sleep period, expressed in hours since midnight of the previous day |
| 13 | wakefulness time | end time of final sleep period, expressed in hours since midnight of the previous day |
| 14 | out bed time | time out of bed, expressed in hours in since midnight of the previous day |
| 15 | gap: sleep onset-inbed | duration spent after in bed and before sleep onset, expressed in minutes |
| 16 | gap: outbed-wakefulness | duration spent after final awake and before getting up, expressed in minutes |
| 17 | deep sleep ratio | ratio of deep sleep duration to total sleep duration |
| 18 | REM sleep ratio | ratio of duration of the rapid eye movement to total sleep duration |

REM, rapid eye movement. The in-house calculation measures include gap: sleep onset -inbed, gap: outbed-wakefulness, deep sleep ratio, REM sleep ratio.

**Table S2. Brain imaging measures**

| **ID** | **Brain volumes** | | | **ID** | **Brain functional connectivity** | | |
| --- | --- | --- | --- | --- | --- | --- | --- |
|  | **brain regions** | **hemisphere** | **atlas** |  | **Cortical network – subcortical region functional connectivity** | | |
|  | **Cortical areas** | | |  | **network 1** | **region 2** | **abbreviation** |
| 1 | bankssts | left | aparc | 184 | Cingulo_opercularNetwork | Lh_pallidus | CON-pal.L |
| 2 | caudal anterior cingulate | left | aparc | 185 | Cingulo_opercularNetwork | Lh_hippocampus | CON-hip.L |
| 3 | caudal middle frontal | left | aparc | 186 | Cingulo_opercularNetwork | Lh_amygda | CON-amy.L |
| 4 | cuneus | left | aparc | 187 | Cingulo_opercularNetwork | Lh_accumbens | CON-acc.L |
| 5 | entorhinal | left | aparc | 188 | Cingulo_opercularNetwork | Lh_ventradc | CON-vdc.L |
| 6 | fusiform | left | aparc | 189 | Cingulo_opercularNetwork | Brain_stem | CON-b.s. |
| 7 | inferior parietal | left | aparc | 190 | Cingulo_opercularNetwork | Rh_thalamus | CON-tha.R |
| 8 | inferior temporal | left | aparc | 191 | Cingulo_opercularNetwork | Rh_caudate | CON-cau.R |
| 9 | isthmus cingulate | left | aparc | 192 | Cingulo_opercularNetwork | Rh_putamen | CON-put.R |
| 10 | lateral occipital | left | aparc | 193 | Cingulo_opercularNetwork | Rh_pallidum | CON-pal.R |
| 11 | lateral orbitofrontal | left | aparc | 194 | Cingulo_opercularNetwork | Rh_hippocampus | CON-hip.R |
| 12 | lingual | left | aparc | 195 | Cingulo_opercularNetwork | Rh_amygdala | CON-amy.R |
| 13 | medial orbitofrontal | left | aparc | 196 | Cingulo_opercularNetwork | Rh_accumbens | CON-acc.R |
| 14 | middle temporal | left | aparc | 197 | Cingulo_opercularNetwork | Rh_ventradc | CON-vdc.R |
| 15 | parahippocampal | left | aparc | 198 | Cingulo_parietalNetwork | Lh_thalamus | CPN-tha.L |
| 16 | paracentral | left | aparc | 199 | Cingulo_parietalNetwork | Lh_caudate | CPN-cau.L |
| 17 | pars opercularis | left | aparc | 200 | Cingulo_parietalNetwork | Lh_putamen | CPN-put.L |
| 18 | pars orbitalis | left | aparc | 201 | Cingulo_parietalNetwork | Lh_pallidus | CPN-pal.L |
| 19 | pars triangularis | left | aparc | 202 | Cingulo_parietalNetwork | Lh_hippocampus | CPN-hip.L |
| 20 | pericalcarine | left | aparc | 203 | Cingulo_parietalNetwork | Lh_amygda | CPN-amy.L |
| 21 | postcentral | left | aparc | 204 | Cingulo_parietalNetwork | Lh_accumbens | CPN-acc.L |
| 22 | posterior cingulate | left | aparc | 205 | Cingulo_parietalNetwork | Lh_ventradc | CPN-vdc.L |
| 23 | precentral | left | aparc | 206 | Cingulo_parietalNetwork | Brain_stem | CPN-b.s. |
| 24 | precuneus | left | aparc | 207 | Cingulo_parietalNetwork | Rh_thalamus | CPN-tha.R |
| 25 | rostral anterior cingulate | left | aparc | 208 | Cingulo_parietalNetwork | Rh_caudate | CPN-cau.R |
| 26 | rostral middle frontal | left | aparc | 209 | Cingulo_parietalNetwork | Rh_putamen | CPN-put.R |
| 27 | superior frontal | left | aparc | 210 | Cingulo_parietalNetwork | Rh_pallidum | CPN-pal.R |
| 28 | superior parietal | left | aparc | 211 | Cingulo_parietalNetwork | Rh_hippocampus | CPN-hip.R |
| 29 | superior temporal | left | aparc | 212 | Cingulo_parietalNetwork | Rh_amygdala | CPN-amy.R |
| 30 | supramarginal | left | aparc | 213 | Cingulo_parietalNetwork | Rh_accumbens | CPN-acc.R |
| 31 | frontal pole | left | aparc | 214 | Cingulo_parietalNetwork | Rh_ventradc | CPN-vdc.R |
| 32 | temporal pole | left | aparc | 215 | DefaultNetwork | Lh_thalamus | DMN-tha.L |
| 33 | transverse temporal | left | aparc | 216 | DefaultNetwork | Lh_caudate | DMN-cau.L |
| 34 | insula | left | aparc | 217 | DefaultNetwork | Lh_putamen | DMN-put.L |
| 35 | bankssts | right | aparc | 218 | DefaultNetwork | Lh_pallidus | DMN-pal.L |
| 36 | caudal anterior cingulate | right | aparc | 219 | DefaultNetwork | Lh_hippocampus | DMN-hip.L |
| 37 | caudal middle frontal | right | aparc | 220 | DefaultNetwork | Lh_amygda | DMN-amy.L |
| 38 | cuneus | right | aparc | 221 | DefaultNetwork | Lh_accumbens | DMN-acc.L |
| 39 | entorhinal | right | aparc | 222 | DefaultNetwork | Lh_ventradc | DMN-vdc.L |
| 40 | fusiform | right | aparc | 223 | DefaultNetwork | Brain_stem | DMN-b.s. |
| 41 | inferior parietal | right | aparc | 224 | DefaultNetwork | Rh_thalamus | DMN-tha.R |
| 42 | inferior temporal | right | aparc | 225 | DefaultNetwork | Rh_caudate | DMN-cau.R |
| 43 | isthmus cingulate | right | aparc | 226 | DefaultNetwork | Rh_putamen | DMN-put.R |
| 44 | lateral occipital | right | aparc | 227 | DefaultNetwork | Rh_pallidum | DMN-pal.R |
| 45 | lateral orbitofrontal | right | aparc | 228 | DefaultNetwork | Rh_hippocampus | DMN-hip.R |
| 46 | lingual | right | aparc | 229 | DefaultNetwork | Rh_amygdala | DMN-amy.R |
| 47 | medial orbitofrontal | right | aparc | 230 | DefaultNetwork | Rh_accumbens | DMN-acc.R |
| 48 | middle temporal | right | aparc | 231 | DefaultNetwork | Rh_ventradc | DMN-vdc.R |
| 49 | parahippocampal | right | aparc | 232 | DorsalAttentionNetwork | Lh_thalamus | DAN-tha.L |
| 50 | paracentral | right | aparc | 233 | DorsalAttentionNetwork | Lh_caudate | DAN-cau.L |
| 51 | pars opercularis | right | aparc | 234 | DorsalAttentionNetwork | Lh_putamen | DAN-put.L |
| 52 | pars orbitalis | right | aparc | 235 | DorsalAttentionNetwork | Lh_pallidus | DAN-pal.L |
| 53 | pars triangularis | right | aparc | 236 | DorsalAttentionNetwork | Lh_hippocampus | DAN-hip.L |
| 54 | pericalcarine | right | aparc | 237 | DorsalAttentionNetwork | Lh_amygda | DAN-amy.L |
| 55 | postcentral | right | aparc | 238 | DorsalAttentionNetwork | Lh_accumbens | DAN-acc.L |
| 56 | posterior cingulate | right | aparc | 239 | DorsalAttentionNetwork | Lh_ventradc | DAN-vdc.L |
| 57 | precentral | right | aparc | 240 | DorsalAttentionNetwork | Brain_stem | DAN-b.s. |
| 58 | precuneus | right | aparc | 241 | DorsalAttentionNetwork | Rh_thalamus | DAN-tha.R |
| 59 | rostral anterior cingulate | right | aparc | 242 | DorsalAttentionNetwork | Rh_caudate | DAN-cau.R |
| 60 | rostral middle frontal | right | aparc | 243 | DorsalAttentionNetwork | Rh_putamen | DAN-put.R |
| 61 | superior frontal | right | aparc | 244 | DorsalAttentionNetwork | Rh_pallidum | DAN-pal.R |
| 62 | superior parietal | right | aparc | 245 | DorsalAttentionNetwork | Rh_hippocampus | DAN-hip.R |
| 63 | superior temporal | right | aparc | 246 | DorsalAttentionNetwork | Rh_amygdala | DAN-amy.R |
| 64 | supramarginal | right | aparc | 247 | DorsalAttentionNetwork | Rh_accumbens | DAN-acc.R |
| 65 | frontal pole | right | aparc | 248 | DorsalAttentionNetwork | Rh_ventradc | DAN-vdc.R |
| 66 | temporal pole | right | aparc | 249 | Fronto_parietalNetwork | Lh_thalamus | FPN-tha.L |
| 67 | transverse temporal | right | aparc | 250 | Fronto_parietalNetwork | Lh_caudate | FPN-cau.L |
| 68 | insula | right | aparc | 251 | Fronto_parietalNetwork | Lh_putamen | FPN-put.L |
|  | **Subcortical areas** | | | 252 | Fronto_parietalNetwork | Lh_pallidus | FPN-pal.L |
| 69 | thalamus proper | left | aseg | 253 | Fronto_parietalNetwork | Lh_hippocampus | FPN-hip.L |
| 70 | caudate | left | aseg | 254 | Fronto_parietalNetwork | Lh_amygda | FPN-amy.L |
| 71 | putamen | left | aseg | 255 | Fronto_parietalNetwork | Lh_accumbens | FPN-acc.L |
| 72 | pallidum | left | aseg | 256 | Fronto_parietalNetwork | Lh_ventradc | FPN-vdc.L |
| 73 | brain stem |  | aseg | 257 | Fronto_parietalNetwork | Brain_stem | FPN-b.s. |
| 74 | hippocampus | left | aseg | 258 | Fronto_parietalNetwork | Rh_thalamus | FPN-tha.R |
| 75 | amygdala | left | aseg | 259 | Fronto_parietalNetwork | Rh_caudate | FPN-cau.R |
| 76 | accumbens area | left | aseg | 260 | Fronto_parietalNetwork | Rh_putamen | FPN-put.R |
| 77 | ventral DC | left | aseg | 261 | Fronto_parietalNetwork | Rh_pallidum | FPN-pal.R |
| 78 | thalamus proper | right | aseg | 262 | Fronto_parietalNetwork | Rh_hippocampus | FPN-hip.R |
| 79 | caudate | right | aseg | 263 | Fronto_parietalNetwork | Rh_amygdala | FPN-amy.R |
| 80 | putamen | right | aseg | 264 | Fronto_parietalNetwork | Rh_accumbens | FPN-acc.R |
| 81 | pallidum | right | aseg | 265 | Fronto_parietalNetwork | Rh_ventradc | FPN-vdc.R |
| 82 | hippocampus | right | aseg | 266 | RetrosplenialTemporalNetwork | Lh_thalamus | RTN-tha.L |
| 83 | amygdala | right | aseg | 267 | RetrosplenialTemporalNetwork | Lh_caudate | RTN-cau.L |
| 84 | accumbens area | right | aseg | 268 | RetrosplenialTemporalNetwork | Lh_putamen | RTN-put.L |
| 85 | ventral DC | right | aseg | 269 | RetrosplenialTemporalNetwork | Lh_pallidus | RTN-pal.L |
| **ID** | **Brain functional network connectivity** | | | 270 | RetrosplenialTemporalNetwork | Lh_hippocampus | RTN-hip.L |
|  | **Cortical network – cortical network functional connectivity** | | | 271 | RetrosplenialTemporalNetwork | Lh_amygda | RTN-amy.L |
|  | **network 1** | **network 2** | **abbreviation** | 272 | RetrosplenialTemporalNetwork | Lh_accumbens | RTN-acc.L |
| 86 | AuditoryNetwork | AuditoryNetwork | AUN-AUN | 273 | RetrosplenialTemporalNetwork | Lh_ventradc | RTN-vdc.L |
| 87 | AuditoryNetwork | Cingulo_opercularNetwork | AUN-CON | 274 | RetrosplenialTemporalNetwork | Brain_stem | RTN-b.s. |
| 88 | AuditoryNetwork | Cingulo_parietalNetwork | AUN-CPN | 275 | RetrosplenialTemporalNetwork | Rh_thalamus | RTN-tha.R |
| 89 | AuditoryNetwork | DefaultNetwork | AUN-DMN | 276 | RetrosplenialTemporalNetwork | Rh_caudate | RTN-cau.R |
| 90 | AuditoryNetwork | DorsalAttentionNetwork | AUN-DAN | 277 | RetrosplenialTemporalNetwork | Rh_putamen | RTN-put.R |
| 91 | AuditoryNetwork | Fronto_parietalNetwork | AUN-FPN | 278 | RetrosplenialTemporalNetwork | Rh_pallidum | RTN-pal.R |
| 92 | AuditoryNetwork | RetrosplenialTemporalNetwork | AUN-RTN | 279 | RetrosplenialTemporalNetwork | Rh_hippocampus | RTN-hip.R |
| 93 | AuditoryNetwork | SalienceNetwork | AUN-SAN | 280 | RetrosplenialTemporalNetwork | Rh_amygdala | RTN-amy.R |
| 94 | AuditoryNetwork | SensorimotorHandNetwork | AUN-SHN | 281 | RetrosplenialTemporalNetwork | Rh_accumbens | RTN-acc.R |
| 95 | AuditoryNetwork | SensorimotorMouthNetwork | AUN-SMN | 282 | RetrosplenialTemporalNetwork | Rh_ventradc | RTN-vdc.R |
| 96 | AuditoryNetwork | VentralAttentionNetwork | AUN-VAN | 283 | SalienceNetwork | Lh_thalamus | SAN-tha.L |
| 97 | AuditoryNetwork | VisualNetwork | AUN-VIN | 284 | SalienceNetwork | Lh_caudate | SAN-cau.L |
| 98 | Cingulo_opercularNetwork | Cingulo_opercularNetwork | CON-CON | 285 | SalienceNetwork | Lh_putamen | SAN-put.L |
| 99 | Cingulo_opercularNetwork | Cingulo_parietalNetwork | CON-CPN | 286 | SalienceNetwork | Lh_pallidus | SAN-pal.L |
| 100 | Cingulo_opercularNetwork | DefaultNetwork | CON-DMN | 287 | SalienceNetwork | Lh_hippocampus | SAN-hip.L |
| 101 | Cingulo_opercularNetwork | DorsalAttentionNetwork | CON-DAN | 288 | SalienceNetwork | Lh_amygda | SAN-amy.L |
| 102 | Cingulo_opercularNetwork | Fronto_parietalNetwork | CON-FPN | 289 | SalienceNetwork | Lh_accumbens | SAN-acc.L |
| 103 | Cingulo_opercularNetwork | RetrosplenialTemporalNetwork | CON-RTN | 290 | SalienceNetwork | Lh_ventradc | SAN-vdc.L |
| 104 | Cingulo_opercularNetwork | SalienceNetwork | CON-SAN | 291 | SalienceNetwork | Brain_stem | SAN-b.s. |
| 105 | Cingulo_opercularNetwork | SensorimotorHandNetwork | CON-SHN | 292 | SalienceNetwork | Rh_thalamus | SAN-tha.R |
| 106 | Cingulo_opercularNetwork | SensorimotorMouthNetwork | CON-SMN | 293 | SalienceNetwork | Rh_caudate | SAN-cau.R |
| 107 | Cingulo_opercularNetwork | VentralAttentionNetwork | CON-VAN | 294 | SalienceNetwork | Rh_putamen | SAN-put.R |
| 108 | Cingulo_opercularNetwork | VisualNetwork | CON-VIN | 295 | SalienceNetwork | Rh_pallidum | SAN-pal.R |
| 109 | Cingulo_parietalNetwork | Cingulo_parietalNetwork | CPN-CPN | 296 | SalienceNetwork | Rh_hippocampus | SAN-hip.R |
| 110 | Cingulo_parietalNetwork | DefaultNetwork | CPN-DMN | 297 | SalienceNetwork | Rh_amygdala | SAN-amy.R |
| 111 | Cingulo_parietalNetwork | DorsalAttentionNetwork | CPN-DAN | 298 | SalienceNetwork | Rh_accumbens | SAN-acc.R |
| 112 | Cingulo_parietalNetwork | Fronto_parietalNetwork | CPN-FPN | 299 | SalienceNetwork | Rh_ventradc | SAN-vdc.R |
| 113 | Cingulo_parietalNetwork | RetrosplenialTemporalNetwork | CPN-RTN | 300 | SensorimotorHandNetwork | Lh_thalamus | SHN-tha.L |
| 114 | Cingulo_parietalNetwork | SalienceNetwork | CPN-SAN | 301 | SensorimotorHandNetwork | Lh_caudate | SHN-cau.L |
| 115 | Cingulo_parietalNetwork | SensorimotorHandNetwork | CPN-SHN | 302 | SensorimotorHandNetwork | Lh_putamen | SHN-put.L |
| 116 | Cingulo_parietalNetwork | SensorimotorMouthNetwork | CPN-SMN | 303 | SensorimotorHandNetwork | Lh_pallidus | SHN-pal.L |
| 117 | Cingulo_parietalNetwork | VentralAttentionNetwork | CPN-VAN | 304 | SensorimotorHandNetwork | Lh_hippocampus | SHN-hip.L |
| 118 | Cingulo_parietalNetwork | VisualNetwork | CPN-VIN | 305 | SensorimotorHandNetwork | Lh_amygda | SHN-amy.L |
| 119 | DefaultNetwork | DefaultNetwork | DMN-DMN | 306 | SensorimotorHandNetwork | Lh_accumbens | SHN-acc.L |
| 120 | DefaultNetwork | DorsalAttentionNetwork | DMN-DAN | 307 | SensorimotorHandNetwork | Lh_ventradc | SHN-vdc.L |
| 121 | DefaultNetwork | Fronto_parietalNetwork | DMN-FPN | 308 | SensorimotorHandNetwork | Brain_stem | SHN-b.s. |
| 122 | DefaultNetwork | RetrosplenialTemporalNetwork | DMN-RTN | 309 | SensorimotorHandNetwork | Rh_thalamus | SHN-tha.R |
| 123 | DefaultNetwork | SalienceNetwork | DMN-SAN | 310 | SensorimotorHandNetwork | Rh_caudate | SHN-cau.R |
| 124 | DefaultNetwork | SensorimotorHandNetwork | DMN-SHN | 311 | SensorimotorHandNetwork | Rh_putamen | SHN-put.R |
| 125 | DefaultNetwork | SensorimotorMouthNetwork | DMN-SMN | 312 | SensorimotorHandNetwork | Rh_pallidum | SHN-pal.R |
| 126 | DefaultNetwork | VentralAttentionNetwork | DMN-VAN | 313 | SensorimotorHandNetwork | Rh_hippocampus | SHN-hip.R |
| 127 | DefaultNetwork | VisualNetwork | DMN-VIN | 314 | SensorimotorHandNetwork | Rh_amygdala | SHN-amy.R |
| 128 | DorsalAttentionNetwork | DorsalAttentionNetwork | DAN-DAN | 315 | SensorimotorHandNetwork | Rh_accumbens | SHN-acc.R |
| 129 | DorsalAttentionNetwork | Fronto_parietalNetwork | DAN-FPN | 316 | SensorimotorHandNetwork | Rh_ventradc | SHN-vdc.R |
| 130 | DorsalAttentionNetwork | RetrosplenialTemporalNetwork | DAN-RTN | 317 | SensorimotorMouthNetwork | Lh_thalamus | SMN-tha.L |
| 131 | DorsalAttentionNetwork | SalienceNetwork | DAN-SAN | 318 | SensorimotorMouthNetwork | Lh_caudate | SMN-cau.L |
| 132 | DorsalAttentionNetwork | SensorimotorHandNetwork | DAN-SHN | 319 | SensorimotorMouthNetwork | Lh_putamen | SMN-put.L |
| 133 | DorsalAttentionNetwork | SensorimotorMouthNetwork | DAN-SMN | 320 | SensorimotorMouthNetwork | Lh_pallidus | SMN-pal.L |
| 134 | DorsalAttentionNetwork | VentralAttentionNetwork | DAN-VAN | 321 | SensorimotorMouthNetwork | Lh_hippocampus | SMN-hip.L |
| 135 | DorsalAttentionNetwork | VisualNetwork | DAN-VIN | 322 | SensorimotorMouthNetwork | Lh_amygda | SMN-amy.L |
| 136 | Fronto_parietalNetwork | Fronto_parietalNetwork | FPN-FPN | 323 | SensorimotorMouthNetwork | Lh_accumbens | SMN-acc.L |
| 137 | Fronto_parietalNetwork | RetrosplenialTemporalNetwork | FPN-RTN | 324 | SensorimotorMouthNetwork | Lh_ventradc | SMN-vdc.L |
| 138 | Fronto_parietalNetwork | SalienceNetwork | FPN-SAN | 325 | SensorimotorMouthNetwork | Brain_stem | SMN-b.s. |
| 139 | Fronto_parietalNetwork | SensorimotorHandNetwork | FPN-SHN | 326 | SensorimotorMouthNetwork | Rh_thalamus | SMN-tha.R |
| 140 | Fronto_parietalNetwork | SensorimotorMouthNetwork | FPN-SMN | 327 | SensorimotorMouthNetwork | Rh_caudate | SMN-cau.R |
| 141 | Fronto_parietalNetwork | VentralAttentionNetwork | FPN-VAN | 328 | SensorimotorMouthNetwork | Rh_putamen | SMN-put.R |
| 142 | Fronto_parietalNetwork | VisualNetwork | FPN-VIN | 329 | SensorimotorMouthNetwork | Rh_pallidum | SMN-pal.R |
| 143 | RetrosplenialTemporalNetwork | RetrosplenialTemporalNetwork | RTN-RTN | 330 | SensorimotorMouthNetwork | Rh_hippocampus | SMN-hip.R |
| 144 | RetrosplenialTemporalNetwork | SalienceNetwork | RTN-SAN | 331 | SensorimotorMouthNetwork | Rh_amygdala | SMN-amy.R |
| 145 | RetrosplenialTemporalNetwork | SensorimotorHandNetwork | RTN-SHN | 332 | SensorimotorMouthNetwork | Rh_accumbens | SMN-acc.R |
| 146 | RetrosplenialTemporalNetwork | SensorimotorMouthNetwork | RTN-SMN | 333 | SensorimotorMouthNetwork | Rh_ventradc | SMN-vdc.R |
| 147 | RetrosplenialTemporalNetwork | VentralAttentionNetwork | RTN-VAN | 334 | VentralAttentionNetwork | Lh_thalamus | VAN-tha.L |
| 148 | RetrosplenialTemporalNetwork | VisualNetwork | RTN-VIN | 335 | VentralAttentionNetwork | Lh_caudate | VAN-cau.L |
| 149 | SalienceNetwork | SalienceNetwork | SAN-SAN | 336 | VentralAttentionNetwork | Lh_putamen | VAN-put.L |
| 150 | SalienceNetwork | SensorimotorHandNetwork | SAN-SHN | 337 | VentralAttentionNetwork | Lh_pallidus | VAN-pal.L |
| 151 | SalienceNetwork | SensorimotorMouthNetwork | SAN-SMN | 338 | VentralAttentionNetwork | Lh_hippocampus | VAN-hip.L |
| 152 | SalienceNetwork | VentralAttentionNetwork | SAN-VAN | 339 | VentralAttentionNetwork | Lh_amygda | VAN-amy.L |
| 153 | SalienceNetwork | VisualNetwork | SAN-VIN | 340 | VentralAttentionNetwork | Lh_accumbens | VAN-acc.L |
| 154 | SensorimotorHandNetwork | SensorimotorHandNetwork | SHN-SHN | 341 | VentralAttentionNetwork | Lh_ventradc | VAN-vdc.L |
| 155 | SensorimotorHandNetwork | SensorimotorMouthNetwork | SHN-SMN | 342 | VentralAttentionNetwork | Brain_stem | VAN-b.s. |
| 156 | SensorimotorHandNetwork | VentralAttentionNetwork | SHN-VAN | 343 | VentralAttentionNetwork | Rh_thalamus | VAN-tha.R |
| 157 | SensorimotorHandNetwork | VisualNetwork | SHN-VIN | 344 | VentralAttentionNetwork | Rh_caudate | VAN-cau.R |
| 158 | SensorimotorMouthNetwork | SensorimotorMouthNetwork | SMN-SMN | 345 | VentralAttentionNetwork | Rh_putamen | VAN-put.R |
| 159 | SensorimotorMouthNetwork | VentralAttentionNetwork | SMN-VAN | 346 | VentralAttentionNetwork | Rh_pallidum | VAN-pal.R |
| 160 | SensorimotorMouthNetwork | VisualNetwork | SMN-VIN | 347 | VentralAttentionNetwork | Rh_hippocampus | VAN-hip.R |
| 161 | VentralAttentionNetwork | VentralAttentionNetwork | VAN-VAN | 348 | VentralAttentionNetwork | Rh_amygdala | VAN-amy.R |
| 162 | VentralAttentionNetwork | VisualNetwork | VAN-VIN | 349 | VentralAttentionNetwork | Rh_accumbens | VAN-acc.R |
| 163 | VisualNetwork | VisualNetwork | VIN-VIN | 350 | VentralAttentionNetwork | Rh_ventradc | VAN-vdc.R |
|  | **Cortical network – subcortical region functional connectivity** | | | 351 | VisualNetwork | Lh_thalamus | VIN-tha.L |
|  | **network 1** | **region 2** | **abbreviation** | 352 | VisualNetwork | Lh_caudate | VIN-cau.L |
| 164 | AuditoryNetwork | Lh_thalamus | AUN-tha.L | 353 | VisualNetwork | Lh_putamen | VIN-put.L |
| 165 | AuditoryNetwork | Lh_caudate | AUN-cau.L | 354 | VisualNetwork | Lh_pallidus | VIN-pal.L |
| 166 | AuditoryNetwork | Lh_putamen | AUN-put.L | 355 | VisualNetwork | Lh_hippocampus | VIN-hip.L |
| 167 | AuditoryNetwork | Lh_pallidus | AUN-pal.L | 356 | VisualNetwork | Lh_amygda | VIN-amy.L |
| 168 | AuditoryNetwork | Lh_hippocampus | AUN-hip.L | 357 | VisualNetwork | Lh_accumbens | VIN-acc.L |
| 169 | AuditoryNetwork | Lh_amygda | AUN-amy.L | 358 | VisualNetwork | Lh_ventradc | VIN-vdc.L |
| 170 | AuditoryNetwork | Lh_accumbens | AUN-acc.L | 359 | VisualNetwork | Brain_stem | VIN-b.s. |
| 171 | AuditoryNetwork | Lh_ventradc | AUN-vdc.L | 360 | VisualNetwork | Rh_thalamus | VIN-tha.R |
| 172 | AuditoryNetwork | Brain_stem | AUN-b.s. | 361 | VisualNetwork | Rh_caudate | VIN-cau.R |
| 173 | AuditoryNetwork | Rh_thalamus | AUN-tha.R | 362 | VisualNetwork | Rh_putamen | VIN-put.R |
| 174 | AuditoryNetwork | Rh_caudate | AUN-cau.R | 363 | VisualNetwork | Rh_pallidum | VIN-pal.R |
| 175 | AuditoryNetwork | Rh_putamen | AUN-put.R | 364 | VisualNetwork | Rh_hippocampus | VIN-hip.R |
| 176 | AuditoryNetwork | Rh_pallidum | AUN-pal.R | 365 | VisualNetwork | Rh_amygdala | VIN-amy.R |
| 177 | AuditoryNetwork | Rh_hippocampus | AUN-hip.R | 366 | VisualNetwork | Rh_accumbens | VIN-acc.R |
| 178 | AuditoryNetwork | Rh_amygdala | AUN-amy.R | 367 | VisualNetwork | Rh_ventradc | VIN-vdc.R |
| 179 | AuditoryNetwork | Rh_accumbens | AUN-acc.R |  |  |  |  |
| 180 | AuditoryNetwork | Rh_ventradc | AUN-vdc.R |  |  |  |  |
| 181 | Cingulo_opercularNetwork | Lh_thalamus | CON-tha.L |  |  |  |  |
| 182 | Cingulo_opercularNetwork | Lh_caudate | CON-cau.L |  |  |  |  |
| 183 | Cingulo_opercularNetwork | Lh_putamen | CON-put.L |  |  |  |  |

**Table S3. Participants’ information**

|  | Discovery sample (N=3300)  2-year follow-up | | Replication sample (N=1271)  4-year follow-up | |
| --- | --- | --- | --- | --- |
| **Demographic information** | **Mean or No.** | **SD or %** | **Mean or No.** | **SD or %** |
| **Age, month** | 143.43 | 7.78 | 168.86 | 8.25 |
| **Sex** |  |  |  |  |
| Male | 1666 | 50.48% | 628 | 49.41% |
| Female | 1634 | 49.52% | 643 | 50.59% |
| **BMI** | 20.16 | 4.47 |  |  |
| **family income, level** | 7.84 | 1.94 | 8.09 | 1.87 |
| **Mother education, years** | 17.04 | 2.43 | 17.04 | 2.44 |
| **Father education, years** | 16.90 | 2.57 | 16.9 | 2.58 |
| **Race** |  |  |  |  |
| White | 1983 | 60.09% | 760 | 59.8% |
| Black | 291 | 8.82% | 103 | 8.10% |
| Hispanic | 637 | 19.30% | 253 | 19.91% |
| Asian | 64 | 1.94% | 29 | 2.28% |
| Others | 325 | 9.85% | 126 | 9.91% |
| **Puberty** | 2.12 | 0.70 | 2.93 | 0.59 |
| **Device based sleep measures** |  |  |  |  |
| sleep duration: total, min | 445.93 | 41.87 | 420.43 | 67.19 |
| sleep duration: wakefulness, min | 57.24 | 10.98 | 54.11 | 11.31 |
| sleep duration: light sleep, min | 259.87 | 31.57 | 246.51 | 34.67 |
| sleep duration: deep sleep, min | 89.39 | 14.70 | 83.90 | 15.10 |
| sleep duration: REM, min | 97.38 | 18.10 | 90.45 | 20.01 |
| number of awakenings | 29.64 | 5.27 | 28.44 | 5.86 |
| heart rate: wakefulness, bpm | 73.55 | 7.60 | 70.71 | 7.64 |
| heart rate: light sleep, bpm | 69.54 | 7.70 | 66.16 | 7.80 |
| heart rate: deep sleep, bpm | 72.37 | 8.46 | 68.57 | 8.71 |
| heart rate: REM, bpm | 72.45 | 7.51 | 68.84 | 7.93 |
| inbed time, hour | 10.96 | 1.26 | 11.95 | 1.47 |
| sleep onset time, hour | 11.08 | 1.25 | 12.06 | 1.47 |
| wakefulness time, hour | 19.02 | 1.51 | 18.85 | 2.31 |
| outbed time, hour | 19.07 | 1.54 | 18.88 | 2.35 |
| gap: sleep onset-inbed, min | 6.88 | 6.63 | 6.78 | 6.84 |
| gap: outbed-wakefulness, min | 3.13 | 18.14 | 2.02 | 18.80 |
| deep sleep ratio, % | 0.20 | 0.03 | 0.20 | 0.03 |
| REM sleep ratio, % | 0.22 | 0.03 | 0.21 | 0.04 |

REM, rapid eye movement; bpm, beats per minute; min, minutes; SD, standard deviaton. All the measures related to time point (inbed time, sleep onset time, wakefulness time, outbed time) were offset by 12 hours so that 00:00=noon, 23:59=11:59 AM

### **Table S4. Loadings for brain volumes and functional network connectivities**

| **ID** | **brain measures** | **Loading for first canonical variate** | **ID** | **brain measures** | **Loadings for second canonical variate** |
| --- | --- | --- | --- | --- | --- |
| 303 | SHN-pal.Lh | **-0.74** | 186 | CON-amy.Lh | **-0.61** |
| 186 | CON-amy.Lh | **-0.73** | 324 | SMN-vdc.Lh | **-0.57** |
| 194 | CON-hip.Rh | **-0.73** | 93 | AUN-SAN | **-0.56** |
| 310 | SHN-cau.Rh | **-0.73** | 208 | CPN-cau.Rh | **-0.56** |
| 175 | AUN-put.Rh | **-0.69** | 313 | SHN-hip.Rh | **-0.48** |
| 182 | CON-cau.Lh | **-0.67** | 152 | SAN-VAN | **-0.47** |
| 198 | CPN-tha.Lh | **-0.66** | 189 | CON-b.s. | **-0.47** |
| 282 | RTN-vdc.Rh | **-0.65** | 301 | SHN-cau.Lh | **-0.47** |
| 155 | SHN-SMN | **0.64** | 326 | SMN-tha.Rh | **-0.46** |
| 311 | SHN-put.Rh | **-0.64** | 271 | RTN-amy.Lh | **0.42** |
| 315 | SHN-acc.Rh | **-0.64** | 355 | VIN-hip.Lh | **-0.41** |
| 183 | CON-put.Lh | **-0.63** | 44 | Rh_lateraloccipital | **-0.4** |
| 195 | CON-amy.Rh | **-0.62** | 74 | Lh_hippocampus | **-0.4** |
| 205 | CPN-vdc.Lh | **-0.61** | 166 | AUN-put.Lh | **0.4** |
| 321 | SMN-hip.Lh | **-0.59** | 48 | Rh_middletemporal | **-0.39** |
| 363 | VIN-pal.Rh | **-0.59** | 107 | CON-VAN | **-0.39** |
| 154 | SHN-SHN | **0.56** | 82 | Rh_hippocampus | **-0.38** |
| 210 | CPN-pal.Rh | **-0.56** | 285 | SAN-put.Lh | **-0.38** |
| 364 | VIN-hip.Rh | **-0.56** | 40 | Rh_fusiform | **-0.37** |
| 170 | AUN-acc.Lh | **-0.55** | 10 | Lh_lateraloccipital | **-0.36** |
| 217 | DMN-put.Lh | **-0.55** | 47 | Rh_medialorbitofrontal | **-0.36** |
| 355 | VIN-hip.Lh | **-0.55** | 87 | AUN-CON | **-0.36** |
| 274 | RTN-b.s. | **-0.54** | 219 | DMN-hip.Lh | **-0.36** |
| 97 | AUN-VIN | **0.53** | 96 | AUN-VAN | **-0.35** |
| 218 | DMN-pal.Lh | **-0.52** | 187 | CON-acc.Lh | **-0.35** |
| 326 | SMN-tha.Rh | **-0.52** | 308 | SHN-b.s. | **-0.34** |
| 94 | AUN-SHN | **0.51** | 12 | Lh_lingual | **-0.33** |
| 157 | SHN-VIN | **0.51** | 172 | AUN-b.s. | **0.33** |
| 283 | SAN-tha.Lh | **-0.51** | 75 | Lh_amygdala | **-0.32** |
| 322 | SMN-amy.Lh | **-0.51** | 30 | Lh_supramarginal | **-0.31** |
| 197 | CON-vdc.Rh | **-0.5** | 68 | Rh_insula | **-0.31** |
| 313 | SHN-hip.Rh | **-0.5** | 267 | RTN-cau.Lh | **0.31** |
| 171 | AUN-vdc.Lh | **-0.49** | 65 | Rh_frontalpole | **-0.3** |
| 93 | AUN-SAN | **-0.48** | 171 | AUN-vdc.Lh | **-0.3** |
| 91 | AUN-FPN | **-0.46** | 203 | CPN-amy.Lh | **-0.3** |
| 150 | SAN-SHN | **-0.46** | 37 | Rh_caudalmiddlefrontal | **-0.29** |
| 221 | DMN-acc.Lh | **-0.46** | 15 | Lh_parahippocampal | **-0.28** |
| 229 | DMN-amy.Rh | **-0.46** | 17 | Lh_parsopercularis | **-0.28** |
| 11 | Lh_lateralorbitofrontal | **-0.45** | 320 | SMN-pal.Lh | **-0.28** |
| 58 | Rh_precuneus | **-0.44** | 201 | CPN-pal.Lh | **-0.27** |
| 139 | FPN-SHN | **-0.44** | 262 | FPN-hip.Rh | **-0.27** |
| 189 | CON-b.s. | **-0.44** | 270 | RTN-hip.Lh | **-0.27** |
| 270 | RTN-hip.Lh | **-0.44** | 340 | VAN-acc.Lh | **0.27** |
| 208 | CPN-cau.Rh | **-0.43** | 80 | Rh_putamen | **-0.26** |
| 230 | DMN-acc.Rh | **-0.43** | 207 | CPN-tha.Rh | **-0.26** |
| 24 | Lh_precuneus | **-0.42** | 323 | SMN-acc.Lh | **-0.26** |
| 48 | Rh_middletemporal | **-0.42** | 32 | Lh_temporalpole | **-0.25** |
| 275 | RTN-tha.Rh | **-0.42** | 192 | CON-put.Rh | **-0.25** |
| 324 | SMN-vdc.Lh | **-0.42** | 317 | SMN-tha.Lh | **-0.25** |
| 23 | Lh_precentral | **-0.41** | 327 | SMN-cau.Rh | **-0.25** |
| 27 | Lh_superiorfrontal | **-0.41** | 98 | CON-CON | **-0.24** |
| 44 | Rh_lateraloccipital | **-0.41** | 168 | AUN-hip.Lh | **0.24** |
| 74 | Lh_hippocampus | **-0.41** | 295 | SAN-pal.Rh | **0.24** |
| 188 | CON-vdc.Lh | **0.41** | 338 | VAN-hip.Lh | **-0.24** |
| 289 | SAN-acc.Lh | **-0.41** | 36 | Rh_caudalanteriorcingulate | **-0.23** |
| 304 | SHN-hip.Lh | **0.41** | 39 | Rh_entorhinal | **-0.23** |
| 45 | Rh_lateralorbitofrontal | **-0.4** | 104 | CON-SAN | **-0.23** |
| 82 | Rh_hippocampus | **-0.4** | 251 | FPN-put.Lh | **-0.23** |
| 180 | AUN-vdc.Rh | **0.4** | 344 | VAN-cau.Rh | **-0.23** |
| 14 | Lh_middletemporal | **-0.39** | 117 | CPN-VAN | **-0.22** |
| 21 | Lh_postcentral | **-0.39** | 294 | SAN-put.Rh | **0.22** |
| 40 | Rh_fusiform | **-0.39** | 165 | AUN-cau.Lh | **0.21** |
| 42 | Rh_inferiortemporal | **-0.39** | 268 | RTN-put.Lh | **0.21** |
| 61 | Rh_superiorfrontal | **-0.39** | 360 | VIN-tha.Rh | **0.21** |
| 63 | Rh_superiortemporal | **-0.39** | 241 | DAN-tha.Rh | **0.2** |
| 69 | Lh_thalamus | **-0.39** | 290 | SAN-vdc.Lh | **0.2** |
| 151 | SAN-SMN | **-0.39** | 334 | VAN-tha.Lh | **-0.18** |
| 176 | AUN-pal.Rh | **-0.39** | 209 | CPN-put.Rh | **0.17** |
| 240 | DAN-b.s. | **-0.39** | 226 | DMN-put.Rh | **0.17** |
| 10 | Lh_lateraloccipital | **-0.38** | 247 | DAN-acc.Rh | **-0.16** |
| 26 | Lh_rostralmiddlefrontal | **-0.38** | 106 | CON-SMN | **-0.15** |
| 57 | Rh_precentral | **-0.38** | 181 | CON-tha.Lh | **-0.15** |
| 166 | AUN-put.Lh | **0.38** | 281 | RTN-acc.Rh | **-0.15** |
| 248 | DAN-vdc.Rh | **-0.38** | 169 | AUN-amy.Lh | **0.12** |
| 279 | RTN-hip.Rh | **0.38** | 202 | CPN-hip.Lh | **-0.12** |
| 308 | SHN-b.s. | **-0.38** | 258 | FPN-tha.Rh | **0.12** |
| 335 | VAN-cau.Lh | **-0.38** | 280 | RTN-amy.Rh | **-0.12** |
| 6 | Lh_fusiform | **-0.37** | 224 | DMN-tha.Rh | **0.11** |
| 29 | Lh_superiortemporal | **-0.37** | 346 | VAN-pal.Rh | **0.11** |
| 34 | Lh_insula | **-0.37** | 349 | VAN-acc.Rh | **0.11** |
| 55 | Rh_postcentral | **-0.37** | 122 | DMN-RTN | **-0.1** |
| 62 | Rh_superiorparietal | **-0.37** | 149 | SAN-SAN | **-0.1** |
| 83 | Rh_amygdala | **-0.37** | 231 | DMN-vdc.Rh | **0.1** |
| 85 | Rh_ventraldc | **-0.37** | 261 | FPN-pal.Rh | **0.1** |
| 152 | SAN-VAN | **-0.37** | 288 | SAN-amy.Lh | **-0.1** |
| 162 | VAN-VIN | **0.37** | 291 | SAN-b.s. | **0.1** |
| 301 | SHN-cau.Lh | **-0.37** | 319 | SMN-put.Lh | **-0.1** |
| 28 | Lh_superiorparietal | **-0.36** | 128 | DAN-DAN | **-0.09** |
| 47 | Rh_medialorbitofrontal | **-0.36** | 235 | DAN-pal.Lh | **-0.09** |
| 95 | AUN-SMN | **0.36** | 129 | DAN-FPN | **-0.08** |
| 8 | Lh_inferiortemporal | **-0.35** | 193 | CON-pal.Rh | **-0.08** |
| 18 | Lh_parsorbitalis | **-0.35** | 269 | RTN-pal.Lh | **0.08** |
| 68 | Rh_insula | **-0.35** | 325 | SMN-b.s. | **-0.06** |
| 75 | Lh_amygdala | **-0.35** | 99 | CON-CPN | **0.05** |
| 77 | Lh_ventraldc | **-0.35** | 190 | CON-tha.Rh | **-0.05** |
| 78 | Rh_thalamus | **-0.35** | 233 | DAN-cau.Lh | **0.05** |
| 16 | Lh_paracentral | **-0.34** | 306 | SHN-acc.Lh | **-0.05** |
| 30 | Lh_supramarginal | **-0.34** | 358 | VIN-vdc.Lh | **0.04** |
| 52 | Rh_parsorbitalis | **-0.34** | 110 | CPN-DMN | **-0.03** |
| 177 | AUN-hip.Rh | **0.34** | 1 | Lh_BanksOfSuperiorTemporalSulcus | -0.21 |
| 187 | CON-acc.Lh | **-0.34** | 2 | Lh_caudalanteriorcingulate | -0.2 |
| 264 | FPN-acc.Rh | **-0.34** | 3 | Lh_caudalmiddlefrontal | -0.29 |
| 316 | SHN-vdc.Rh | **-0.34** | 4 | Lh_cuneus | -0.3 |
| 25 | Lh_rostralanteriorcingulate | **-0.33** | 5 | Lh_entorhinal | -0.22 |
| 98 | CON-CON | **-0.33** | 6 | Lh_fusiform | -0.36 |
| 163 | VIN-VIN | **-0.33** | 7 | Lh_inferiorparietal | -0.25 |
| 199 | CPN-cau.Lh | **-0.33** | 8 | Lh_inferiortemporal | -0.32 |
| 351 | VIN-tha.Lh | **-0.33** | 9 | Lh_isthmuscingulate | -0.28 |
| 3 | Lh_caudalmiddlefrontal | **-0.32** | 11 | Lh_lateralorbitofrontal | -0.4 |
| 9 | Lh_isthmuscingulate | **-0.32** | 13 | Lh_medialorbitofrontal | -0.29 |
| 12 | Lh_lingual | **-0.32** | 14 | Lh_middletemporal | -0.34 |
| 38 | Rh_cuneus | **-0.32** | 16 | Lh_paracentral | -0.32 |
| 41 | Rh_inferiorparietal | **-0.32** | 18 | Lh_parsorbitalis | -0.31 |
| 64 | Rh_supramarginal | **-0.32** | 19 | Lh_parstriangularis | -0.21 |
| 79 | Rh_caudate | **-0.32** | 20 | Lh_pericalcarine | -0.26 |
| 89 | AUN-DMN | **-0.32** | 21 | Lh_postcentral | -0.33 |
| 140 | FPN-SMN | **-0.32** | 22 | Lh_posteriorcingulate | -0.27 |
| 276 | RTN-cau.Rh | **0.32** | 23 | Lh_precentral | -0.35 |
| 277 | RTN-put.Rh | **0.32** | 24 | Lh_precuneus | -0.39 |
| 46 | Rh_lingual | **-0.31** | 25 | Lh_rostralanteriorcingulate | -0.3 |
| 70 | Lh_caudate | **-0.31** | 26 | Lh_rostralmiddlefrontal | -0.35 |
| 256 | FPN-vdc.Lh | **-0.31** | 27 | Lh_superiorfrontal | -0.4 |
| 360 | VIN-tha.Rh | **0.31** | 28 | Lh_superiorparietal | -0.32 |
| 50 | Rh_paracentral | **-0.3** | 29 | Lh_superiortemporal | -0.33 |
| 72 | Lh_pallidum | **-0.3** | 31 | Lh_frontalpole | -0.28 |
| 73 | brain stem | **-0.3** | 33 | Lh_transversetemporal | -0.26 |
| 174 | AUN-cau.Rh | **-0.3** | 34 | Lh_insula | -0.33 |
| 211 | CPN-hip.Rh | **-0.3** | 35 | Rh_BanksOfSuperiorTemporalSulcus | -0.22 |
| 219 | DMN-hip.Lh | **-0.3** | 38 | Rh_cuneus | -0.33 |
| 7 | Lh_inferiorparietal | **-0.29** | 41 | Rh_inferiorparietal | -0.28 |
| 13 | Lh_medialorbitofrontal | **-0.29** | 42 | Rh_inferiortemporal | -0.36 |
| 17 | Lh_parsopercularis | **-0.29** | 43 | Rh_isthmuscingulate | -0.27 |
| 43 | Rh_isthmuscingulate | **-0.29** | 45 | Rh_lateralorbitofrontal | -0.37 |
| 4 | Lh_cuneus | **-0.28** | 46 | Rh_lingual | -0.31 |
| 15 | Lh_parahippocampal | **-0.28** | 49 | Rh_parahippocampal | -0.27 |
| 33 | Lh_transversetemporal | **-0.28** | 50 | Rh_paracentral | -0.28 |
| 49 | Rh_parahippocampal | **-0.28** | 51 | Rh_parsopercularis | -0.27 |
| 65 | Rh_frontalpole | **-0.28** | 52 | Rh_parsorbitalis | -0.31 |
| 80 | Rh_putamen | **-0.28** | 53 | Rh_parstriangularis | -0.2 |
| 84 | Rh_accumbens | **-0.28** | 54 | Rh_pericalcarine | -0.26 |
| 124 | DMN-SHN | **-0.28** | 55 | Rh_postcentral | -0.31 |
| 250 | FPN-cau.Lh | **-0.28** | 56 | Rh_posteriorcingulate | -0.27 |
| 271 | RTN-amy.Lh | **0.28** | 57 | Rh_precentral | -0.33 |
| 342 | VAN-b.s. | **-0.28** | 58 | Rh_precuneus | -0.41 |
| 343 | VAN-tha.Rh | **0.28** | 59 | Rh_rostralanteriorcingulate | -0.23 |
| 20 | Lh_pericalcarine | **-0.27** | 60 | Rh_rostralmiddlefrontal | -0.34 |
| 35 | Rh_BanksOfSuperiorTemporalSulcus | **-0.27** | 61 | Rh_superiorfrontal | -0.37 |
| 51 | Rh_parsopercularis | **-0.27** | 62 | Rh_superiorparietal | -0.33 |
| 54 | Rh_pericalcarine | **-0.27** | 63 | Rh_superiortemporal | -0.35 |
| 67 | Rh_transversetemporal | **-0.27** | 64 | Rh_supramarginal | -0.28 |
| 76 | Lh_accumbens | **-0.27** | 66 | Rh_temporalpole | -0.25 |
| 120 | DMN-DAN | **0.27** | 67 | Rh_transversetemporal | -0.26 |
| 285 | SAN-put.Lh | **-0.27** | 69 | Lh_thalamus | -0.35 |
| 302 | SHN-put.Lh | **-0.27** | 70 | Lh_caudate | -0.26 |
| 366 | VIN-acc.Rh | **-0.27** | 71 | Lh_putamen | -0.23 |
| 31 | Lh_frontalpole | **-0.26** | 72 | Lh_pallidum | -0.25 |
| 87 | AUN-CON | **-0.26** | 73 | brain stem | -0.29 |
| 103 | CON-RTN | **0.26** | 76 | Lh_accumbens | -0.29 |
| 200 | CPN-put.Lh | **0.26** | 77 | Lh_ventraldc | -0.32 |
| 255 | FPN-acc.Lh | **0.26** | 78 | Rh_thalamus | -0.33 |
| 300 | SHN-tha.Lh | **0.26** | 79 | Rh_caudate | -0.28 |
| 357 | VIN-acc.Lh | **0.26** | 81 | Rh_pallidum | -0.22 |
| 71 | Lh_putamen | **-0.25** | 83 | Rh_amygdala | -0.35 |
| 126 | DMN-VAN | **-0.25** | 84 | Rh_accumbens | -0.28 |
| 134 | DAN-VAN | **0.25** | 85 | Rh_ventraldc | -0.33 |
| 143 | RTN-RTN | **-0.25** | 86 | AUN-AUN | -0.08 |
| 92 | AUN-RTN | **0.24** | 88 | AUN-CPN | -0.13 |
| 236 | DAN-hip.Lh | **0.24** | 89 | AUN-DMN | -0.31 |
| 39 | Rh_entorhinal | **-0.23** | 90 | AUN-DAN | 0.08 |
| 244 | DAN-pal.Rh | **0.23** | 91 | AUN-FPN | -0.4 |
| 298 | SAN-acc.Rh | **0.23** | 92 | AUN-RTN | 0.12 |
| 328 | SMN-put.Rh | **0.23** | 94 | AUN-SHN | 0.36 |
| 365 | VIN-amy.Rh | **0.23** | 95 | AUN-SMN | 0.2 |
| 367 | VIN-vdc.Rh | **-0.23** | 97 | AUN-VIN | 0.41 |
| 119 | DMN-DMN | **-0.22** | 100 | CON-DMN | -0.12 |
| 135 | DAN-VIN | **-0.22** | 101 | CON-DAN | 0.04 |
| 165 | AUN-cau.Lh | **0.22** | 102 | CON-FPN | -0.15 |
| 184 | CON-pal.Lh | **0.22** | 103 | CON-RTN | 0.17 |
| 290 | SAN-vdc.Lh | **0.22** | 105 | CON-SHN | -0.16 |
| 336 | VAN-put.Lh | **0.22** | 108 | CON-VIN | 0.25 |
| 5 | Lh_entorhinal | **-0.21** | 109 | CPN-CPN | 0.09 |
| 105 | CON-SHN | **-0.21** | 111 | CPN-DAN | -0.03 |
| 168 | AUN-hip.Lh | **0.21** | 112 | CPN-FPN | 0.13 |
| 212 | CPN-amy.Rh | **0.21** | 113 | CPN-RTN | -0.05 |
| 340 | VAN-acc.Lh | **0.21** | 114 | CPN-SAN | 0.21 |
| 262 | FPN-hip.Rh | **-0.2** | 115 | CPN-SHN | -0.18 |
| 287 | SAN-hip.Lh | **0.2** | 116 | CPN-SMN | -0.17 |
| 338 | VAN-hip.Lh | **-0.2** | 118 | CPN-VIN | -0.05 |
| 114 | CPN-SAN | **0.19** | 119 | DMN-DMN | -0.1 |
| 128 | DAN-DAN | **-0.19** | 120 | DMN-DAN | 0.07 |
| 251 | FPN-put.Lh | **-0.18** | 121 | DMN-FPN | 0.04 |
| 259 | FPN-cau.Rh | **-0.18** | 123 | DMN-SAN | -0.08 |
| 112 | CPN-FPN | **0.17** | 124 | DMN-SHN | -0.21 |
| 141 | FPN-VAN | **-0.17** | 125 | DMN-SMN | -0.13 |
| 246 | DAN-amy.Rh | **0.17** | 126 | DMN-VAN | -0.13 |
| 107 | CON-VAN | **-0.16** | 127 | DMN-VIN | 0.04 |
| 96 | AUN-VAN | **-0.15** | 130 | DAN-RTN | -0.01 |
| 117 | CPN-VAN | **-0.15** | 131 | DAN-SAN | 0.03 |
| 127 | DMN-VIN | **0.14** | 132 | DAN-SHN | -0.07 |
| 293 | SAN-cau.Rh | **-0.13** | 133 | DAN-SMN | 0.01 |
| 362 | VIN-put.Rh | **-0.12** | 134 | DAN-VAN | 0.04 |
| 169 | AUN-amy.Lh | **0.03** | 135 | DAN-VIN | -0.12 |
| 269 | RTN-pal.Lh | **-0.01** | 136 | FPN-FPN | 0.06 |
| 1 | Lh_BanksOfSuperiorTemporalSulcus | -0.22 | 137 | FPN-RTN | 0.03 |
| 2 | Lh_caudalanteriorcingulate | -0.21 | 138 | FPN-SAN | 0.05 |
| 19 | Lh_parstriangularis | -0.22 | 139 | FPN-SHN | -0.38 |
| 22 | Lh_posteriorcingulate | -0.29 | 140 | FPN-SMN | -0.3 |
| 32 | Lh_temporalpole | -0.21 | 141 | FPN-VAN | -0.19 |
| 36 | Rh_caudalanteriorcingulate | -0.21 | 142 | FPN-VIN | -0.05 |
| 37 | Rh_caudalmiddlefrontal | -0.3 | 143 | RTN-RTN | -0.16 |
| 53 | Rh_parstriangularis | -0.21 | 144 | RTN-SAN | 0.02 |
| 56 | Rh_posteriorcingulate | -0.25 | 145 | RTN-SHN | 0.14 |
| 59 | Rh_rostralanteriorcingulate | -0.23 | 146 | RTN-SMN | 0.12 |
| 60 | Rh_rostralmiddlefrontal | -0.36 | 147 | RTN-VAN | 0.03 |
| 66 | Rh_temporalpole | -0.25 | 148 | RTN-VIN | -0.12 |
| 81 | Rh_pallidum | -0.24 | 150 | SAN-SHN | -0.36 |
| 86 | AUN-AUN | 0.11 | 151 | SAN-SMN | -0.38 |
| 88 | AUN-CPN | -0.2 | 153 | SAN-VIN | 0.05 |
| 90 | AUN-DAN | 0.1 | 154 | SHN-SHN | 0.38 |
| 99 | CON-CPN | -0.07 | 155 | SHN-SMN | 0.43 |
| 100 | CON-DMN | 0.09 | 156 | SHN-VAN | 0.02 |
| 101 | CON-DAN | -0.03 | 157 | SHN-VIN | 0.33 |
| 102 | CON-FPN | 0 | 158 | SMN-SMN | 0.08 |
| 104 | CON-SAN | -0.12 | 159 | SMN-VAN | -0.11 |
| 106 | CON-SMN | -0.1 | 160 | SMN-VIN | 0.17 |
| 108 | CON-VIN | 0.18 | 161 | VAN-VAN | -0.15 |
| 109 | CPN-CPN | 0.01 | 162 | VAN-VIN | 0.25 |
| 110 | CPN-DMN | 0.11 | 163 | VIN-VIN | -0.19 |
| 111 | CPN-DAN | -0.13 | 164 | AUN-tha.Lh | 0.07 |
| 113 | CPN-RTN | -0.1 | 167 | AUN-pal.Lh | -0.06 |
| 115 | CPN-SHN | -0.25 | 170 | AUN-acc.Lh | -0.37 |
| 116 | CPN-SMN | -0.22 | 173 | AUN-tha.Rh | 0.15 |
| 118 | CPN-VIN | -0.16 | 174 | AUN-cau.Rh | -0.25 |
| 121 | DMN-FPN | 0.09 | 175 | AUN-put.Rh | -0.53 |
| 122 | DMN-RTN | -0.05 | 176 | AUN-pal.Rh | -0.28 |
| 123 | DMN-SAN | 0.01 | 177 | AUN-hip.Rh | 0.21 |
| 125 | DMN-SMN | -0.18 | 178 | AUN-amy.Rh | 0.01 |
| 129 | DAN-FPN | 0.04 | 179 | AUN-acc.Rh | 0 |
| 130 | DAN-RTN | -0.14 | 180 | AUN-vdc.Rh | 0.18 |
| 131 | DAN-SAN | 0.08 | 182 | CON-cau.Lh | -0.38 |
| 132 | DAN-SHN | -0.07 | 183 | CON-put.Lh | -0.38 |
| 133 | DAN-SMN | 0.09 | 184 | CON-pal.Lh | 0 |
| 136 | FPN-FPN | 0.07 | 185 | CON-hip.Lh | -0.05 |
| 137 | FPN-RTN | 0.04 | 188 | CON-vdc.Lh | 0.22 |
| 138 | FPN-SAN | 0.11 | 191 | CON-cau.Rh | 0.09 |
| 142 | FPN-VIN | -0.04 | 194 | CON-hip.Rh | -0.47 |
| 144 | RTN-SAN | 0.14 | 195 | CON-amy.Rh | -0.43 |
| 145 | RTN-SHN | 0.16 | 196 | CON-acc.Rh | -0.11 |
| 146 | RTN-SMN | 0.14 | 197 | CON-vdc.Rh | -0.45 |
| 147 | RTN-VAN | 0.2 | 198 | CPN-tha.Lh | -0.54 |
| 148 | RTN-VIN | -0.19 | 199 | CPN-cau.Lh | -0.24 |
| 149 | SAN-SAN | 0 | 200 | CPN-put.Lh | 0.14 |
| 153 | SAN-VIN | -0.02 | 204 | CPN-acc.Lh | -0.06 |
| 156 | SHN-VAN | 0.1 | 205 | CPN-vdc.Lh | -0.31 |
| 158 | SMN-SMN | 0.12 | 206 | CPN-b.s. | 0.05 |
| 159 | SMN-VAN | -0.01 | 210 | CPN-pal.Rh | -0.48 |
| 160 | SMN-VIN | 0.22 | 211 | CPN-hip.Rh | -0.26 |
| 161 | VAN-VAN | -0.21 | 212 | CPN-amy.Rh | 0.13 |
| 164 | AUN-tha.Lh | -0.14 | 213 | CPN-acc.Rh | -0.02 |
| 167 | AUN-pal.Lh | -0.09 | 214 | CPN-vdc.Rh | -0.03 |
| 172 | AUN-b.s. | 0.27 | 215 | DMN-tha.Lh | -0.12 |
| 173 | AUN-tha.Rh | -0.01 | 216 | DMN-cau.Lh | 0.01 |
| 178 | AUN-amy.Rh | 0.2 | 217 | DMN-put.Lh | -0.35 |
| 179 | AUN-acc.Rh | -0.07 | 218 | DMN-pal.Lh | -0.35 |
| 181 | CON-tha.Lh | -0.11 | 220 | DMN-amy.Lh | 0.05 |
| 185 | CON-hip.Lh | -0.17 | 221 | DMN-acc.Lh | -0.3 |
| 190 | CON-tha.Rh | 0.01 | 222 | DMN-vdc.Lh | -0.11 |
| 191 | CON-cau.Rh | 0.23 | 223 | DMN-b.s. | -0.08 |
| 192 | CON-put.Rh | -0.16 | 225 | DMN-cau.Rh | -0.04 |
| 193 | CON-pal.Rh | 0.09 | 227 | DMN-pal.Rh | -0.01 |
| 196 | CON-acc.Rh | 0.06 | 228 | DMN-hip.Rh | -0.06 |
| 201 | CPN-pal.Lh | -0.01 | 229 | DMN-amy.Rh | -0.28 |
| 202 | CPN-hip.Lh | 0.12 | 230 | DMN-acc.Rh | -0.23 |
| 203 | CPN-amy.Lh | -0.06 | 232 | DAN-tha.Lh | -0.08 |
| 204 | CPN-acc.Lh | 0.11 | 234 | DAN-put.Lh | 0.08 |
| 206 | CPN-b.s. | 0.02 | 236 | DAN-hip.Lh | 0.2 |
| 207 | CPN-tha.Rh | 0.03 | 237 | DAN-amy.Lh | -0.04 |
| 209 | CPN-put.Rh | 0.07 | 238 | DAN-acc.Lh | -0.21 |
| 213 | CPN-acc.Rh | 0.14 | 239 | DAN-vdc.Lh | -0.15 |
| 214 | CPN-vdc.Rh | 0 | 240 | DAN-b.s. | -0.25 |
| 215 | DMN-tha.Lh | 0.06 | 242 | DAN-cau.Rh | -0.16 |
| 216 | DMN-cau.Lh | 0.16 | 243 | DAN-put.Rh | -0.11 |
| 220 | DMN-amy.Lh | -0.01 | 244 | DAN-pal.Rh | 0.14 |
| 222 | DMN-vdc.Lh | -0.2 | 245 | DAN-hip.Rh | 0.07 |
| 223 | DMN-b.s. | 0.12 | 246 | DAN-amy.Rh | 0.08 |
| 224 | DMN-tha.Rh | 0.12 | 248 | DAN-vdc.Rh | -0.3 |
| 225 | DMN-cau.Rh | -0.1 | 249 | FPN-tha.Lh | -0.08 |
| 226 | DMN-put.Rh | 0.16 | 250 | FPN-cau.Lh | -0.25 |
| 227 | DMN-pal.Rh | 0.06 | 252 | FPN-pal.Lh | 0.22 |
| 228 | DMN-hip.Rh | -0.06 | 253 | FPN-hip.Lh | -0.2 |
| 231 | DMN-vdc.Rh | 0.12 | 254 | FPN-amy.Lh | -0.07 |
| 232 | DAN-tha.Lh | -0.14 | 255 | FPN-acc.Lh | 0.18 |
| 233 | DAN-cau.Lh | 0.04 | 256 | FPN-vdc.Lh | -0.24 |
| 234 | DAN-put.Lh | 0.19 | 257 | FPN-b.s. | 0.21 |
| 235 | DAN-pal.Lh | 0 | 259 | FPN-cau.Rh | -0.18 |
| 237 | DAN-amy.Lh | -0.08 | 260 | FPN-put.Rh | 0.12 |
| 238 | DAN-acc.Lh | -0.25 | 263 | FPN-amy.Rh | -0.12 |
| 239 | DAN-vdc.Lh | -0.11 | 264 | FPN-acc.Rh | -0.19 |
| 241 | DAN-tha.Rh | 0.11 | 265 | FPN-vdc.Rh | -0.12 |
| 242 | DAN-cau.Rh | -0.09 | 266 | RTN-tha.Lh | -0.01 |
| 243 | DAN-put.Rh | -0.15 | 272 | RTN-acc.Lh | 0.21 |
| 245 | DAN-hip.Rh | 0.09 | 273 | RTN-vdc.Lh | -0.22 |
| 247 | DAN-acc.Rh | -0.08 | 274 | RTN-b.s. | -0.36 |
| 249 | FPN-tha.Lh | -0.05 | 275 | RTN-tha.Rh | -0.27 |
| 252 | FPN-pal.Lh | 0.16 | 276 | RTN-cau.Rh | 0.22 |
| 253 | FPN-hip.Lh | -0.1 | 277 | RTN-put.Rh | 0.06 |
| 254 | FPN-amy.Lh | -0.12 | 278 | RTN-pal.Rh | -0.07 |
| 257 | FPN-b.s. | 0.15 | 279 | RTN-hip.Rh | 0.17 |
| 258 | FPN-tha.Rh | 0.17 | 282 | RTN-vdc.Rh | -0.37 |
| 260 | FPN-put.Rh | 0.13 | 283 | SAN-tha.Lh | -0.34 |
| 261 | FPN-pal.Rh | 0.13 | 284 | SAN-cau.Lh | -0.06 |
| 263 | FPN-amy.Rh | -0.1 | 286 | SAN-pal.Lh | 0.02 |
| 265 | FPN-vdc.Rh | -0.22 | 287 | SAN-hip.Lh | 0.1 |
| 266 | RTN-tha.Lh | 0 | 289 | SAN-acc.Lh | -0.29 |
| 267 | RTN-cau.Lh | 0.29 | 292 | SAN-tha.Rh | -0.22 |
| 268 | RTN-put.Lh | 0.1 | 293 | SAN-cau.Rh | -0.16 |
| 272 | RTN-acc.Lh | -0.03 | 296 | SAN-hip.Rh | -0.19 |
| 273 | RTN-vdc.Lh | -0.21 | 297 | SAN-amy.Rh | -0.12 |
| 278 | RTN-pal.Rh | -0.17 | 298 | SAN-acc.Rh | 0.14 |
| 280 | RTN-amy.Rh | 0.08 | 299 | SAN-vdc.Rh | 0.14 |
| 281 | RTN-acc.Rh | -0.11 | 300 | SHN-tha.Lh | 0.02 |
| 284 | SAN-cau.Lh | 0.15 | 302 | SHN-put.Lh | -0.09 |
| 286 | SAN-pal.Lh | -0.06 | 303 | SHN-pal.Lh | -0.6 |
| 288 | SAN-amy.Lh | -0.02 | 304 | SHN-hip.Lh | 0.23 |
| 291 | SAN-b.s. | 0.07 | 305 | SHN-amy.Lh | -0.04 |
| 292 | SAN-tha.Rh | -0.25 | 307 | SHN-vdc.Lh | 0.09 |
| 294 | SAN-put.Rh | 0.1 | 309 | SHN-tha.Rh | -0.05 |
| 295 | SAN-pal.Rh | 0.15 | 310 | SHN-cau.Rh | -0.47 |
| 296 | SAN-hip.Rh | -0.09 | 311 | SHN-put.Rh | -0.44 |
| 297 | SAN-amy.Rh | -0.13 | 312 | SHN-pal.Rh | -0.09 |
| 299 | SAN-vdc.Rh | 0.15 | 314 | SHN-amy.Rh | 0 |
| 305 | SHN-amy.Lh | 0.12 | 315 | SHN-acc.Rh | -0.53 |
| 306 | SHN-acc.Lh | -0.02 | 316 | SHN-vdc.Rh | -0.25 |
| 307 | SHN-vdc.Lh | 0.21 | 318 | SMN-cau.Lh | 0.04 |
| 309 | SHN-tha.Rh | 0.12 | 321 | SMN-hip.Lh | -0.29 |
| 312 | SHN-pal.Rh | 0.07 | 322 | SMN-amy.Lh | -0.29 |
| 314 | SHN-amy.Rh | -0.06 | 328 | SMN-put.Rh | 0.17 |
| 317 | SMN-tha.Lh | 0.03 | 329 | SMN-pal.Rh | 0 |
| 318 | SMN-cau.Lh | 0 | 330 | SMN-hip.Rh | -0.04 |
| 319 | SMN-put.Lh | 0.14 | 331 | SMN-amy.Rh | 0.05 |
| 320 | SMN-pal.Lh | -0.05 | 332 | SMN-acc.Rh | -0.06 |
| 323 | SMN-acc.Lh | 0.04 | 333 | SMN-vdc.Rh | 0.03 |
| 325 | SMN-b.s. | 0.12 | 335 | VAN-cau.Lh | -0.32 |
| 327 | SMN-cau.Rh | -0.29 | 336 | VAN-put.Lh | 0.18 |
| 329 | SMN-pal.Rh | 0.18 | 337 | VAN-pal.Lh | -0.09 |
| 330 | SMN-hip.Rh | -0.03 | 339 | VAN-amy.Lh | 0.27 |
| 331 | SMN-amy.Rh | 0.19 | 341 | VAN-vdc.Lh | -0.2 |
| 332 | SMN-acc.Rh | 0.1 | 342 | VAN-b.s. | -0.24 |
| 333 | SMN-vdc.Rh | 0.16 | 343 | VAN-tha.Rh | 0.19 |
| 334 | VAN-tha.Lh | -0.1 | 345 | VAN-put.Rh | -0.05 |
| 337 | VAN-pal.Lh | -0.08 | 347 | VAN-hip.Rh | -0.17 |
| 339 | VAN-amy.Lh | 0.24 | 348 | VAN-amy.Rh | 0.11 |
| 341 | VAN-vdc.Lh | -0.11 | 350 | VAN-vdc.Rh | -0.22 |
| 344 | VAN-cau.Rh | -0.31 | 351 | VIN-tha.Lh | -0.18 |
| 345 | VAN-put.Rh | -0.02 | 352 | VIN-cau.Lh | -0.1 |
| 346 | VAN-pal.Rh | 0.12 | 353 | VIN-put.Lh | -0.05 |
| 347 | VAN-hip.Rh | -0.16 | 354 | VIN-pal.Lh | -0.22 |
| 348 | VAN-amy.Rh | 0.11 | 356 | VIN-amy.Lh | -0.11 |
| 349 | VAN-acc.Rh | 0.11 | 357 | VIN-acc.Lh | 0.15 |
| 350 | VAN-vdc.Rh | -0.25 | 359 | VIN-b.s. | 0.08 |
| 352 | VIN-cau.Lh | -0.21 | 361 | VIN-cau.Rh | -0.15 |
| 353 | VIN-put.Lh | 0.03 | 362 | VIN-put.Rh | -0.17 |
| 354 | VIN-pal.Lh | -0.21 | 363 | VIN-pal.Rh | -0.34 |
| 356 | VIN-amy.Lh | -0.15 | 364 | VIN-hip.Rh | -0.34 |
| 358 | VIN-vdc.Lh | 0.14 | 365 | VIN-amy.Rh | 0.08 |
| 359 | VIN-b.s. | 0.07 | 366 | VIN-acc.Rh | -0.26 |
| 361 | VIN-cau.Rh | -0.05 | 367 | VIN-vdc.Rh | -0.13 |

Bolded loading values imply that their 95% and 99% confidence intervals did not cross zero which were considered significant. The above sort the loading values in descending order according to both loading values and significance. AUN, Auditory Network; CON, Cingulo_opercular Network; CPN, Cingulo_parietal Network; DMN, Default Network; DAN, Dorsal Attention Network; FPN, Fronto_parietal Network; RTN, Retrosplenial Temporal Network; SAN, Salience Network; SHN, Sensorimotor Hand Network; SMN, Sensorimotor Mouth Network; VAN, Ventral Attention Network; VIN, Visual Network. tha., thalamus, cau., caudate; put., putamen; pal., (globus) pallidus; b.s., brain stem; hip., hippocampus; amy., amygdala; acc., accumbens; vdc., ventral diencephalon; Lh, left hemisphere; Rh, right hemisphere.

**Table S4. Sleep differences across three adolescent biotypes**

|  | Main effect | | | biotype 2 - biotype 1 | | | biotype 3 - biotype 1 | | biotype 3 - biotype 2 | |
| --- | --- | --- | --- | --- | --- | --- | --- | --- | --- | --- |
| Sleep measures | F value | *P* value | t value | | *P* value | T value | | *P* value | t value | *P* value |
| sleep duration: total, min | 46.29 | <0.001 | 5.84 | | 5.60E-09 | 9.61 | | 1.36E-21 | 4.34 | 1.47E-05 |
| sleep duration: wakefulness, min | 21.71 | <0.001 | 3.29 | | 0.001 | 6.57 | | 6.02E-11 | 3.71 | 0.0002 |
| sleep duration: light sleep, min | 34.09 | <0.001 | 4.51 | | 6.69E-06 | 8.25 | | 2.21E-16 | 4.26 | 2.06E-05 |
| sleep duration: deep sleep, min | 7.21 | 0.0008 | 2.89 | | 0.004 | 3.67 | | 0.0002 | 0.95 | 0.34 |
| sleep duration: REM, min | 10.88 | 2.00E-05 | 3.29 | | 0.001 | 4.59 | | 4.53E-06 | 1.54 | 0.12 |
| number of awakenings | 26.79 | <0.001 | 5.13 | | 3.04E-07 | 7.22 | | 6.6E-13 | 2.47 | 0.01 |
| heart rate: wakefulness, bpm | 1.88 | 0.15 | -0.60 | | 0.55 | -1.85 | | 0.06 | -1.40 | 0.16 |
| heart rate: light sleep, bpm | 2.99 | 0.05 | -1.16 | | 0.25 | -2.43 | | 0.02 | -1.43 | 0.15 |
| heart rate: deep sleep, bpm | 3.43 | 0.03 | -0.77 | | 0.44 | -2.49 | | 0.01 | -1.92 | 0.05 |
| heart rate: REM, bpm | 1.96 | 0.14 | -1.09 | | 0.28 | -1.98 | | 0.05 | -1.02 | 0.31 |
| inbed time, hour | 23.52 | <0.001 | -2.77 | | 0.006 | -6.73 | | 1.99E-11 | -4.45 | 9.04E-06 |
| sleep onset time, hour | 24.39 | <0.001 | -2.87 | | 0.004 | -6.87 | | 7.98E-12 | -4.49 | 7.46E-06 |
| wakefulness time, hour | 1.74 | 0.18 | 1.59 | | 0.11 | 0.11 | | 0.91 | -1.57 | 0.12 |
| outbed time, hour | 1.35 | 0.26 | 1.38 | | 0.17 | 0.05 | | 0.96 | -1.41 | 0.16 |
| gap: sleep onset-inbed, min | 0.71 | 0.49 | -0.94 | | 0.35 | -1.14 | | 0.25 | -0.26 | 0.80 |
| gap: outbed-wakefulness, min | 0.46 | 0.6321 | -0.91 | | 0.36 | -0.28 | | 0.78 | 0.66 | 0.51 |
| deep sleep ratio, % | 2.07 | 0.13 | -0.52 | | 0.60 | -1.91 | | 0.06 | -1.54 | 0.12 |
| REM sleep ratio, % | 0.28 | 0.75 | 0.66 | | 0.51 | 0.09 | | 0.93 | -0.61 | 0.54 |

### **Table S5. Differences of sleep characteristics across three adolescent biotypes**

| Sleep characteristics | F value | P value | biotype 2 - biotype 1 | | biotype 3 - biotype 2 | | biotype 3 - biotype 1 | | biotype1 (N) | biotype2 (N) | biotype3 (N) | total number |
| --- | --- | --- | --- | --- | --- | --- | --- | --- | --- | --- | --- | --- |
|  |  |  | t value | p value | t value | p value | t value | p value |  |  |  |  |
| sleep duration: total | 46.29 | <0.001 | 5.84 | 5.61E-09 | 4.34 | 0.00001 | 9.61 | 1.36E-21 | 1058 | 1049 | 1193 | 3300 |
| sleep duration: wakefulness | 21.71 | <0.001 | 3.29 | 0.001 | 3.71 | 0.0002 | 6.57 | 6.02E-11 | 1058 | 1049 | 1193 | 3300 |
| sleep duration: light sleep | 34.09 | <0.001 | 4.51 | 0.000007 | 4.26 | 0.00002 | 8.25 | 2.21E-16 | 1058 | 1049 | 1193 | 3300 |
| sleep duration: deep sleep | 7.21 | 0.0008 | 2.89 | 0.004 | 0.95 | 0.34 | 3.67 | 0.0002 | 1058 | 1049 | 1193 | 3300 |
| sleep duration: REM | 10.88 | 2.00E-05 | 3.29 | 1.01E-03 | 1.54 | 0.12 | 4.59 | 4.53E-06 | 1058 | 1049 | 1193 | 3300 |
| number of awakenings | 26.79 | <0.001 | 5.13 | 0.0000003 | 2.47 | 0.01 | 7.22 | 6.6E-13 | 1058 | 1049 | 1193 | 3300 |
| heart rate: wakefulness | 1.88 | 0.15 | -0.59 | 0.55 | -1.40 | 0.16 | -1.85 | 0.06 | 1058 | 1049 | 1193 | 3300 |
| heart rate: light sleep | 2.99 | 0.05 | -1.16 | 0.25 | -1.43 | 0.15 | -2.43 | 0.02 | 1058 | 1049 | 1193 | 3300 |
| heart rate: deep sleep | 3.43 | 0.03 | -0.77 | 0.44 | -1.92 | 0.05 | -2.49 | 0.01 | 1058 | 1049 | 1193 | 3300 |
| heart rate: REM | 1.96 | 0.14 | -1.09 | 0.28 | -1.02 | 0.31 | -1.98 | 0.05 | 1058 | 1049 | 1193 | 3300 |
| inbed time | 23.52 | <0.001 | -0.94 | 0.35 | -0.26 | 0.80 | -1.14 | 0.25 | 1058 | 1049 | 1193 | 3300 |
| sleep onset time | 24.39 | <0.001 | -0.91 | 0.36 | 0.66 | 0.51 | -0.28 | 0.78 | 1058 | 1049 | 1193 | 3300 |
| wakefulness time | 1.74 | 0.18 | -2.77 | 0.006 | -4.45 | 0.000009 | -6.73 | 1.99E-11 | 1058 | 1049 | 1193 | 3300 |
| out bed time | 1.35 | 0.26 | -2.87 | 0.004 | -4.49 | 0.000007 | -6.86 | 7.98E-12 | 1058 | 1049 | 1193 | 3300 |
| gap: sleep onset-inbed | 0.71 | 0.49 | 1.38 | 0.17 | -1.41 | 0.16 | 0.05 | 0.96 | 1058 | 1049 | 1193 | 3300 |
| gap: outbed-wakefulness | 0.46 | 0.6321 | 1.59 | 0.11 | -1.57 | 0.12 | 0.11 | 0.91 | 1058 | 1049 | 1193 | 3300 |
| deep sleep ratio | 2.07 | 0.13 | -0.52 | 0.60 | -1.54 | 0.12 | -1.91 | 0.06 | 1058 | 1049 | 1193 | 3300 |
| REM sleep ratio | 0.28 | 0.75 | 0.66 | 0.51 | -0.61 | 0.54 | 0.09 | 0.93 | 1058 | 1049 | 1193 | 3300 |

### **Table S6. Cognitive differences across three adolescent biotypes**

| cognitive performance | F value | *P* vale | biotype 2 – biotype 1 | | biotype 3 – biotype 2 | | biotype 3 -biotype 1 | | biotype 1 (N) | biotype 2 (N) | biotype 3 (N) | total number |
| --- | --- | --- | --- | --- | --- | --- | --- | --- | --- | --- | --- | --- |
|  |  |  | t value | *P* value | t value | *P* value | t value | *P* value |  |  |  |  |
| picture vocabulary | 20.04 | <0.001 | 2.55 | 0.01 | 6.21 | 6.03E-10 | 4.11 | 4.05E-05 | 1024 | 1012 | 1159 | 3195 |
| flanker | 2.90 | 0.06 | 1.51 | 0.13 | 2.40 | 0.02 | 1.03 | 0.30 | 1024 | 1010 | 1146 | 3180 |
| pattern | 0.96 | 0.38 | 0.08 | 0.93 | -1.09 | 0.28 | -1.28 | 0.20 | 1017 | 1008 | 1139 | 3164 |
| picture | 1.14 | 0.32 | 0.84 | 0.40 | 1.51 | 0.13 | 0.76 | 0.45 | 1041 | 1028 | 1170 | 3239 |
| reading | 13.35 | <0.001 | 2.40 | 0.02 | 5.13 | 3.15E-07 | 3.08 | 0.002 | 1020 | 1010 | 1155 | 3185 |
| crystalized intelligence | 22.21 | <0.001 | 2.71 | 0.007 | 6.55 | 6.76E-11 | 4.30 | 1.72E-05 | 1011 | 999 | 1124 | 3134 |

### **Table S7. Brain differences across three adolescent biotypes**

| ID | brain measures | F value | *P* value | biotype 2 - biotype 1 | | biotype 3 - biotype 2 | | biotype 3 - biotype 1 | |
| --- | --- | --- | --- | --- | --- | --- | --- | --- | --- |
|  |  |  |  | t value | *P* value | t value | *P* value | t value | *P* value |
| 1 | Lh_BanksOfSuperiorTemporalSulcus | 45.58 | <0.001 | 5.28 | 1.37E-07 | 9.55 | 2.56E-21 | 4.86 | 1.20E-06 |
| 2 | Lh_caudalanteriorcingulate | 35.88 | <0.001 | 2.66 | 7.86E-03 | 8.12 | 6.54E-16 | 6.08 | 1.30E-09 |
| 3 | Lh_caudalmiddlefrontal | 87.39 | <0.001 | 6.88 | 7.34E-12 | 13.2 | 8.87E-39 | 7.18 | 8.88E-13 |
| 4 | Lh_cuneus | 90.97 | <0.001 | 5.57 | 2.82E-08 | 13.26 | 3.77E-39 | 8.64 | 8.52E-18 |
| 5 | Lh_entorhinal | 26.45 | <0.001 | 3.36 | 7.85E-04 | 7.21 | 6.75E-13 | 4.34 | 1.44E-05 |
| 6 | Lh_fusiform | 122.95 | <0.001 | 8.18 | 3.95E-16 | 15.65 | 2.52E-53 | 8.49 | 3.21E-17 |
| 7 | Lh_inferiorparietal | 61.66 | <0.001 | 5.44 | 5.75E-08 | 11.05 | 6.51E-28 | 6.35 | 2.43E-10 |
| 8 | Lh_inferiortemporal | 83.59 | <0.001 | 7.5 | 7.92E-14 | 12.93 | 2.46E-37 | 6.22 | 5.74E-10 |
| 9 | Lh_isthmuscingulate | 95.5 | <0.001 | 5.46 | 5.23E-08 | 13.54 | 1.13E-40 | 9.06 | 2.20E-19 |
| 10 | Lh_lateraloccipital | 118.8 | <0.001 | 6.83 | 1.01E-11 | 15.24 | 9.69E-51 | 9.47 | 5.13E-21 |
| 11 | Lh_lateralorbitofrontal | 171.04 | <0.001 | 7.58 | 4.35E-14 | 18.18 | 1.99E-70 | 11.89 | 6.00E-32 |
| 12 | Lh_lingual | 110.56 | <0.001 | 5.72 | 1.16E-08 | 14.53 | 2.02E-46 | 9.87 | 1.15E-22 |
| 13 | Lh_medialorbitofrontal | 110.42 | <0.001 | 5.11 | 3.47E-07 | 14.37 | 1.85E-45 | 10.34 | 1.07E-24 |
| 14 | Lh_middletemporal | 108.21 | <0.001 | 7.36 | 2.30E-13 | 14.66 | 3.54E-47 | 8.27 | 1.98E-16 |
| 15 | Lh_parahippocampal | 72.45 | <0.001 | 4.95 | 7.76E-07 | 11.83 | 1.15E-31 | 7.73 | 1.47E-14 |
| 16 | Lh_paracentral | 116.66 | <0.001 | 5.08 | 4.00E-07 | 14.72 | 1.45E-47 | 10.76 | 1.46E-26 |
| 17 | Lh_parsopercularis | 82.98 | <0.001 | 6.97 | 3.75E-12 | 12.87 | 4.87E-37 | 6.72 | 2.13E-11 |
| 18 | Lh_parsorbitalis | 88.02 | <0.001 | 6.47 | 1.11E-10 | 13.2 | 8.03E-39 | 7.61 | 3.50E-14 |
| 19 | Lh_parstriangularis | 58.27 | <0.001 | 4.71 | 2.55E-06 | 10.66 | 4.04E-26 | 6.69 | 2.54E-11 |
| 20 | Lh_pericalcarine | 84.23 | <0.001 | 5.28 | 1.35E-07 | 12.75 | 2.29E-36 | 8.38 | 8.06E-17 |
| 21 | Lh_postcentral | 116.84 | <0.001 | 6.11 | 1.14E-09 | 14.99 | 3.52E-49 | 9.96 | 4.79E-23 |
| 22 | Lh_posteriorcingulate | 87.2 | <0.001 | 4.49 | 7.44E-06 | 12.76 | 2.06E-36 | 9.23 | 4.69E-20 |
| 23 | Lh_precentral | 113.15 | <0.001 | 5.1 | 3.53E-07 | 14.53 | 2.12E-46 | 10.52 | 1.74E-25 |
| 24 | Lh_precuneus | 173.04 | <0.001 | 8.1 | 7.61E-16 | 18.37 | 7.77E-72 | 11.55 | 2.69E-30 |
| 25 | Lh_rostralanteriorcingulate | 110.56 | <0.001 | 6.21 | 5.87E-10 | 14.64 | 4.81E-47 | 9.46 | 5.61E-21 |
| 26 | Lh_rostralmiddlefrontal | 144.87 | <0.001 | 6.64 | 3.57E-11 | 16.66 | 7.22E-60 | 11.22 | 1.07E-28 |
| 27 | Lh_superiorfrontal | 180.12 | <0.001 | 8.16 | 4.69E-16 | 18.73 | 2.02E-74 | 11.88 | 6.83E-32 |
| 28 | Lh_superiorparietal | 83.51 | <0.001 | 5.52 | 3.66E-08 | 12.74 | 2.42E-36 | 8.12 | 6.52E-16 |
| 29 | Lh_superiortemporal | 139.28 | <0.001 | 7.54 | 6.26E-14 | 16.53 | 5.30E-59 | 10.13 | 9.00E-24 |
| 30 | Lh_supramarginal | 87.57 | <0.001 | 6.39 | 1.89E-10 | 13.16 | 1.35E-38 | 7.65 | 2.54E-14 |
| 31 | Lh_frontalpole | 57.88 | <0.001 | 4 | 6.43E-05 | 10.48 | 2.60E-25 | 7.25 | 5.11E-13 |
| 32 | Lh_temporalpole | 60.37 | <0.001 | 6.38 | 2.01E-10 | 10.99 | 1.31E-27 | 5.28 | 1.38E-07 |
| 33 | Lh_transversetemporal | 72.15 | <0.001 | 6.33 | 2.80E-10 | 12 | 1.78E-32 | 6.44 | 1.38E-10 |
| 34 | Lh_insula | 124.84 | <0.001 | 6.84 | 9.58E-12 | 15.6 | 5.79E-53 | 9.85 | 1.39E-22 |
| 35 | Rh_BanksOfSuperiorTemporalSulcus | 43.27 | <0.001 | 4.43 | 9.95E-06 | 9.24 | 4.15E-20 | 5.44 | 5.63E-08 |
| 36 | Rh_caudalanteriorcingulate | 55.91 | <0.001 | 4.49 | 7.34E-06 | 10.42 | 4.81E-25 | 6.67 | 3.07E-11 |
| 37 | Rh_caudalmiddlefrontal | 82.21 | <0.001 | 5.62 | 2.09E-08 | 12.67 | 6.00E-36 | 7.93 | 2.89E-15 |
| 38 | Rh_cuneus | 117.43 | <0.001 | 5.19 | 2.26E-07 | 14.8 | 5.13E-48 | 10.73 | 2.06E-26 |
| 39 | Rh_entorhinal | 32.69 | <0.001 | 2.9 | 3.70E-03 | 7.85 | 5.51E-15 | 5.53 | 3.44E-08 |
| 40 | Rh_fusiform | 149.32 | <0.001 | 7.44 | 1.23E-13 | 17.05 | 1.53E-62 | 10.8 | 9.34E-27 |
| 41 | Rh_inferiorparietal | 73.75 | <0.001 | 6.53 | 7.69E-11 | 12.13 | 3.51E-33 | 6.38 | 2.01E-10 |
| 42 | Rh_inferiortemporal | 124.58 | <0.001 | 8.9 | 8.76E-19 | 15.78 | 3.72E-54 | 7.86 | 5.09E-15 |
| 43 | Rh_isthmuscingulate | 96.44 | <0.001 | 5.83 | 6.14E-09 | 13.68 | 1.91E-41 | 8.81 | 1.91E-18 |
| 44 | Rh_lateraloccipital | 142.02 | <0.001 | 6.46 | 1.22E-10 | 16.46 | 1.37E-58 | 11.21 | 1.24E-28 |
| 45 | Rh_lateralorbitofrontal | 141.46 | <0.001 | 6.98 | 3.54E-12 | 16.55 | 3.90E-59 | 10.74 | 1.78E-26 |
| 46 | Rh_lingual | 92.93 | <0.001 | 5.53 | 3.51E-08 | 13.39 | 7.99E-40 | 8.82 | 1.88E-18 |
| 47 | Rh_medialorbitofrontal | 158.28 | <0.001 | 5.95 | 2.91E-09 | 17.16 | 2.78E-63 | 12.51 | 4.28E-35 |
| 48 | Rh_middletemporal | 133.93 | <0.001 | 8.58 | 1.41E-17 | 16.34 | 9.13E-58 | 8.81 | 1.94E-18 |
| 49 | Rh_parahippocampal | 75.77 | <0.001 | 5.79 | 7.92E-09 | 12.22 | 1.25E-33 | 7.27 | 4.59E-13 |
| 50 | Rh_paracentral | 83.32 | <0.001 | 5.55 | 3.14E-08 | 12.73 | 2.69E-36 | 8.08 | 8.88E-16 |
| 51 | Rh_parsopercularis | 73.43 | <0.001 | 6.5 | 9.15E-11 | 12.11 | 4.78E-33 | 6.38 | 2.02E-10 |
| 52 | Rh_parsorbitalis | 84.95 | <0.001 | 6.13 | 9.65E-10 | 12.94 | 2.07E-37 | 7.69 | 1.96E-14 |
| 53 | Rh_parstriangularis | 55.63 | <0.001 | 4.04 | 5.56E-05 | 10.3 | 1.59E-24 | 7.02 | 2.71E-12 |
| 54 | Rh_pericalcarine | 100.28 | <0.001 | 5.2 | 2.16E-07 | 13.78 | 4.86E-42 | 9.6 | 1.51E-21 |
| 55 | Rh_postcentral | 107.19 | <0.001 | 5.97 | 2.61E-09 | 14.38 | 1.56E-45 | 9.44 | 6.90E-21 |
| 56 | Rh_posteriorcingulate | 82.26 | <0.001 | 5.49 | 4.33E-08 | 12.65 | 7.58E-36 | 8.05 | 1.16E-15 |
| 57 | Rh_precentral | 106.46 | <0.001 | 6.08 | 1.37E-09 | 14.36 | 2.16E-45 | 9.3 | 2.48E-20 |
| 58 | Rh_precuneus | 183.11 | <0.001 | 8.59 | 1.35E-17 | 18.94 | 5.09E-76 | 11.66 | 8.02E-31 |
| 59 | Rh_rostralanteriorcingulate | 48.23 | <0.001 | 4.08 | 4.70E-05 | 9.66 | 8.55E-22 | 6.27 | 4.03E-10 |
| 60 | Rh_rostralmiddlefrontal | 124.78 | <0.001 | 6.51 | 8.53E-11 | 15.53 | 1.48E-52 | 10.13 | 9.45E-24 |
| 61 | Rh_superiorfrontal | 138.77 | <0.001 | 7.51 | 7.46E-14 | 16.5 | 8.60E-59 | 10.12 | 9.95E-24 |
| 62 | Rh_superiorparietal | 96.53 | <0.001 | 6.39 | 1.91E-10 | 13.78 | 5.09E-42 | 8.33 | 1.16E-16 |
| 63 | Rh_superiortemporal | 147.67 | <0.001 | 7.66 | 2.35E-14 | 17 | 3.35E-62 | 10.51 | 1.86E-25 |
| 64 | Rh_supramarginal | 92.05 | <0.001 | 6.29 | 3.55E-10 | 13.46 | 3.07E-40 | 8.09 | 8.59E-16 |
| 65 | Rh_frontalpole | 70.22 | <0.001 | 4.23 | 2.40E-05 | 11.5 | 4.83E-30 | 8.13 | 6.15E-16 |
| 66 | Rh_temporalpole | 74.26 | <0.001 | 6.04 | 1.69E-09 | 12.14 | 3.37E-33 | 6.9 | 6.19E-12 |
| 67 | Rh_transversetemporal | 60.85 | <0.001 | 5.84 | 5.65E-09 | 11.02 | 9.55E-28 | 5.88 | 4.42E-09 |
| 68 | Rh_insula | 120.38 | <0.001 | 6.59 | 5.16E-11 | 15.29 | 4.69E-51 | 9.78 | 2.69E-22 |
| 69 | Lh_thalamus | 167.59 | <0.001 | 7.43 | 1.33E-13 | 17.98 | 5.34E-69 | 11.83 | 1.20E-31 |
| 70 | Lh_caudate | 77.66 | <0.001 | 5.75 | 9.67E-09 | 12.36 | 2.48E-34 | 7.45 | 1.16E-13 |
| 71 | Lh_putamen | 61.42 | <0.001 | 5.63 | 1.90E-08 | 11.05 | 6.62E-28 | 6.14 | 9.14E-10 |
| 72 | Lh_pallidum | 92.11 | <0.001 | 5.13 | 3.09E-07 | 13.24 | 4.88E-39 | 9.08 | 1.77E-19 |
| 73 | brain stem | 108.03 | <0.001 | 5.03 | 5.18E-07 | 14.21 | 1.66E-44 | 10.25 | 2.80E-24 |
| 74 | Lh_hippocampus | 175.72 | <0.001 | 8.24 | 2.55E-16 | 18.53 | 5.86E-73 | 11.58 | 2.01E-30 |
| 75 | Lh_amygdala | 108.02 | <0.001 | 6.01 | 2.08E-09 | 14.44 | 7.03E-46 | 9.46 | 5.47E-21 |
| 76 | Lh_accumbens | 92.94 | <0.001 | 5.24 | 1.74E-07 | 13.32 | 1.79E-39 | 9.06 | 2.26E-19 |
| 77 | Lh_ventraldc | 141.4 | <0.001 | 7.15 | 1.06E-12 | 16.58 | 2.49E-59 | 10.59 | 8.26E-26 |
| 78 | Rh_thalamus | 156.29 | <0.001 | 7.02 | 2.72E-12 | 17.33 | 1.99E-64 | 11.56 | 2.59E-30 |
| 79 | Rh_caudate | 88.31 | <0.001 | 6.37 | 2.09E-10 | 13.21 | 7.20E-39 | 7.73 | 1.47E-14 |
| 80 | Rh_putamen | 82.37 | <0.001 | 5.52 | 3.71E-08 | 12.66 | 6.46E-36 | 8.03 | 1.30E-15 |
| 81 | Rh_pallidum | 77.91 | <0.001 | 7.11 | 1.44E-12 | 12.48 | 5.69E-35 | 6.15 | 8.87E-10 |
| 82 | Rh_hippocampus | 166.89 | <0.001 | 8.98 | 4.60E-19 | 18.19 | 1.67E-70 | 10.42 | 4.83E-25 |
| 83 | Rh_amygdala | 157.94 | <0.001 | 7.78 | 9.53E-15 | 17.56 | 4.79E-66 | 11 | 1.13E-27 |
| 84 | Rh_accumbens | 95.65 | <0.001 | 6.29 | 3.58E-10 | 13.7 | 1.33E-41 | 8.35 | 9.61E-17 |
| 85 | Rh_ventraldc | 149.99 | <0.001 | 7.33 | 2.88E-13 | 17.07 | 1.23E-62 | 10.94 | 2.20E-27 |
| 86 | AUN-AUN | 16.62 | <0.001 | -4.18 | 2.98E-05 | 0.87 | 3.85E-01 | 5.39 | 7.36E-08 |
| 87 | AUN-CON | 145.84 | <0.001 | 7.18 | 8.82E-13 | 16.82 | 5.89E-61 | 10.83 | 7.24E-27 |
| 88 | AUN-CPN | 37.85 | <0.001 | 6.87 | 7.46E-12 | 8.31 | 1.44E-16 | 1.81 | 7.06E-02 |
| 89 | AUN-DMN | 103.58 | <0.001 | 7.94 | 2.72E-15 | 14.39 | 1.43E-45 | 7.35 | 2.51E-13 |
| 90 | AUN-DAN | 7.31 | 0.0007 | -0.72 | 0.47 | -3.49 | 0.0005 | -3.07 | 0.002 |
| 91 | AUN-FPN | 238.51 | <0.001 | 13.91 | 8.68E-43 | 21.77 | 3.03E-98 | 9.1 | 1.52E-19 |
| 92 | AUN-RTN | 37.4 | <0.001 | -4.96 | 7.33E-07 | -8.65 | 8.04E-18 | -4.22 | 2.55E-05 |
| 93 | AUN-SAN | 457.4 | <0.001 | 15.54 | 1.35E-52 | 30.17 | 9.23E-177 | 16.59 | 1.90E-59 |
| 94 | AUN-SHN | 239.59 | <0.001 | -16.98 | 4.88E-62 | -21.04 | 2.59E-92 | -5.05 | 4.76E-07 |
| 95 | AUN-SMN | 61.96 | <0.001 | -9.13 | 1.20E-19 | -10.45 | 3.78E-25 | -1.76 | 7.81E-02 |
| 96 | AUN-VAN | 114.87 | <0.001 | 2.51 | 1.22E-02 | 13.7 | 1.39E-41 | 12.37 | 2.31E-34 |
| 97 | AUN-VIN | 275.16 | <0.001 | -12.96 | 1.65E-37 | -23.45 | 1.36E-112 | -11.96 | 2.72E-32 |
| 98 | CON-CON | 60.25 | <0.001 | 3.87 | 1.12E-04 | 10.64 | 5.07E-26 | 7.56 | 5.05E-14 |
| 99 | CON-CPN | 2.16 | 0.12 | -0.13 | 0.89 | -1.78 | 0.08 | -1.81 | 0.07 |
| 100 | CON-DMN | 13.61 | <0.001 | 2.08 | 3.80E-02 | 5.11 | 3.33E-07 | 3.41 | 6.67E-04 |
| 101 | CON-DAN | 2.42 | 0.09 | -1.25 | 0.21 | -2.2 | 0.028 | -1.08 | 0.28 |
| 102 | CON-FPN | 26.56 | <0.001 | 3.4 | 6.72E-04 | 7.23 | 5.82E-13 | 4.32 | 1.61E-05 |
| 103 | CON-RTN | 52.22 | <0.001 | -4.14 | 3.49E-05 | -10.03 | 2.33E-23 | -6.6 | 4.63E-11 |
| 104 | CON-SAN | 74.03 | <0.001 | 4.32 | 1.61E-05 | 11.8 | 1.61E-31 | 8.36 | 9.14E-17 |
| 105 | CON-SHN | 46.09 | <0.001 | 2.61 | 9.02E-03 | 9.07 | 1.92E-19 | 7.18 | 8.60E-13 |
| 106 | CON-SMN | 30.82 | <0.001 | 2.41 | 1.59E-02 | 7.51 | 7.58E-14 | 5.68 | 1.50E-08 |
| 107 | CON-VAN | 146 | <0.001 | 6.01 | 2.00E-09 | 16.56 | 3.08E-59 | 11.78 | 2.10E-31 |
| 108 | CON-VIN | 111.17 | <0.001 | -5.49 | 4.23E-08 | -14.51 | 2.57E-46 | -10.09 | 1.39E-23 |
| 109 | CPN-CPN | 14.39 | <0.001 | -1.84 | 6.55E-02 | -5.19 | 2.25E-07 | -3.73 | 1.91E-04 |
| 110 | CPN-DMN | 0.33 | 0.72 | -0.8 | 0.42 | -0.59 | 0.56 | 0.21 | 0.83 |
| 111 | CPN-DAN | 1.77 | 0.17 | 0.74 | 0.46 | 1.84 | 0.07 | 1.24 | 0.22 |
| 112 | CPN-FPN | 21.45 | <0.001 | -3.87 | 1.13E-04 | -6.55 | 6.77E-11 | -3.08 | 2.12E-03 |
| 113 | CPN-RTN | 6.89 | <0.001 | 0.5 | 0.62 | 3.3 | 0.001 | 3.09 | 0.002 |
| 114 | CPN-SAN | 42.97 | <0.001 | -5.33 | 1.05E-07 | -9.27 | 3.27E-20 | -4.51 | 6.78E-06 |
| 115 | CPN-SHN | 58.66 | <0.001 | 6.38 | 2.01E-10 | 10.83 | 7.08E-27 | 5.1 | 3.57E-07 |
| 116 | CPN-SMN | 46.35 | <0.001 | 5.91 | 3.71E-09 | 9.61 | 1.33E-21 | 4.27 | 2.05E-05 |
| 117 | CPN-VAN | 38.53 | <0.001 | 5.35 | 9.59E-08 | 8.77 | 2.84E-18 | 3.94 | 8.31E-05 |
| 118 | CPN-VIN | 7.56 | <0.001 | 2.91 | 0.004 | 3.78 | 0.0002 | 1.05 | 0.29 |
| 119 | DMN-DMN | 10.27 | <0.001 | -0.26 | 7.95E-01 | 3.56 | 3.70E-04 | 4.19 | 2.90E-05 |
| 120 | DMN-DAN | 5.28 | 0.01 | -1.91 | 0.06 | -3.25 | 0.001 | -1.54 | 0.12 |
| 121 | DMN-FPN | 9.67 | <0.001 | -4.33 | 1.50E-05 | -3.09 | 2.03E-03 | 1.22 | 2.24E-01 |
| 122 | DMN-RTN | 5.12 | 0.01 | 1.3 | 0.19 | 3.14 | 0.002 | 2.07 | 0.04 |
| 123 | DMN-SAN | 0.77 | 0.46 | -1.21 | 0.23 | -0.44 | 0.66 | 0.8 | 0.42 |
| 124 | DMN-SHN | 78.26 | <0.001 | 9.14 | 1.11E-19 | 12.24 | 1.08E-33 | 3.72 | 2.05E-04 |
| 125 | DMN-SMN | 21.53 | <0.001 | 5.42 | 6.57E-08 | 6.14 | 9.49E-10 | 0.98 | 3.28E-01 |
| 126 | DMN-VAN | 10.99 | <0.001 | 1.88 | 6.01E-02 | 4.6 | 4.40E-06 | 3.05 | 2.33E-03 |
| 127 | DMN-VIN | 3.95 | 0.02 | 1.25 | 0.21 | -1.35 | 0.18 | -2.81 | 0.005 |
| 128 | DAN-DAN | 8.03 | <0.001 | 1.54 | 1.25E-01 | 3.92 | 9.19E-05 | 2.66 | 7.77E-03 |
| 129 | DAN-FPN | 1.9 | 0.15 | 1.24 | 0.21 | 1.94 | 0.05 | 0.81 | 0.42 |
| 130 | DAN-RTN | 2.43 | 0.09 | 1.66 | 0.1 | 2.14 | 0.03 | 0.58 | 0.56 |
| 131 | DAN-SAN | 1.89 | 0.15 | -1.62 | 0.11 | -1.81 | 0.07 | -0.27 | 0.78 |
| 132 | DAN-SHN | 8.8 | <0.001 | 2.45 | 1.45E-02 | 4.2 | 2.79E-05 | 2 | 4.52E-02 |
| 133 | DAN-SMN | 0.81 | 0.45 | -0.39 | 0.7 | -1.21 | 0.22 | -0.92 | 0.36 |
| 134 | DAN-VAN | 4.71 | 0.01 | -2.57 | 0.01 | -2.85 | 0.004 | -0.39 | 0.7 |
| 135 | DAN-VIN | 12.24 | <0.001 | 2.57 | 1.01E-02 | 4.94 | 8.27E-07 | 2.68 | 7.29E-03 |
| 136 | FPN-FPN | 14.69 | <0.001 | -5.08 | 3.97E-07 | -4.45 | 8.78E-06 | 0.51 | 6.08E-01 |
| 137 | FPN-RTN | 1.3 | 0.27 | 0.55 | 0.58 | -0.93 | 0.35 | -1.6 | 0.11 |
| 138 | FPN-SAN | 22.02 | <0.001 | -6.1 | 1.21E-09 | -5.64 | 1.87E-08 | 0.29 | 7.70E-01 |
| 139 | FPN-SHN | 225.29 | <0.001 | 14.21 | 1.70E-44 | 21.06 | 1.78E-92 | 8.01 | 1.54E-15 |
| 140 | FPN-SMN | 129.08 | <0.001 | 11.37 | 2.12E-29 | 15.81 | 2.38E-54 | 5.27 | 1.44E-07 |
| 141 | FPN-VAN | 36.65 | <0.001 | 3.57 | 3.64E-04 | 8.43 | 5.30E-17 | 5.45 | 5.35E-08 |
| 142 | FPN-VIN | 16.14 | <0.001 | 5.6 | 2.34E-08 | 3.99 | 6.63E-05 | -1.57 | 1.17E-01 |
| 143 | RTN-RTN | 37.22 | <0.001 | 3.66 | 2.52E-04 | 8.5 | 2.74E-17 | 5.44 | 5.84E-08 |
| 144 | RTN-SAN | 6.1 | <0.001 | -0.94 | 0.35 | -3.3 | 0.001 | -2.62 | 0.009 |
| 145 | RTN-SHN | 33.85 | <0.001 | -3.94 | 8.22E-05 | -8.18 | 4.01E-16 | -4.79 | 1.78E-06 |
| 146 | RTN-SMN | 28.01 | <0.001 | -3.11 | 1.91E-03 | -7.36 | 2.27E-13 | -4.78 | 1.86E-06 |
| 147 | RTN-VAN | 9.38 | <0.001 | -2.23 | 2.61E-02 | -4.32 | 1.59E-05 | -2.38 | 1.75E-02 |
| 148 | RTN-VIN | 28.07 | <0.001 | 2.91 | 3.68E-03 | 7.33 | 2.92E-13 | 4.95 | 7.71E-07 |
| 149 | SAN-SAN | 3.07 | 0.05 | 0.36 | 0.72 | 2.22 | 0.03 | 2.05 | 0.04 |
| 150 | SAN-SHN | 238.9 | <0.001 | 12.52 | 3.81E-35 | 21.86 | 5.29E-99 | 10.68 | 3.28E-26 |
| 151 | SAN-SMN | 215.91 | <0.001 | 11.59 | 1.86E-30 | 20.78 | 3.44E-90 | 10.48 | 2.56E-25 |
| 152 | SAN-VAN | 213.43 | <0.001 | 9.33 | 1.92E-20 | 20.46 | 1.18E-87 | 12.53 | 3.09E-35 |
| 153 | SAN-VIN | 8.82 | <0.001 | 3.76 | 0.0002 | 0.6 | 0.55 | -3.33 | 0.0009 |
| 154 | SHN-SHN | 268.58 | <0.001 | -16.92 | 1.30E-61 | -22.67 | 8.03E-106 | -6.9 | 6.34E-12 |
| 155 | SHN-SMN | 351.39 | <0.001 | -18.12 | 5.14E-70 | -26.24 | 6.76E-138 | -9.53 | 2.93E-21 |
| 156 | SHN-VAN | 2.33 | 0.1 | -1.97 | 0.05 | -1.86 | 0.06 | 0.05 | 0.95 |
| 157 | SHN-VIN | 218.68 | <0.001 | -13.39 | 7.19E-40 | -20.83 | 1.20E-90 | -8.63 | 9.64E-18 |
| 158 | SMN-SMN | 9.35 | <0.001 | -4.28 | 0.00002 | -2.96 | 0.003 | 1.3 | 0.19 |
| 159 | SMN-VAN | 11.96 | <0.001 | 0.71 | 4.79E-01 | 4.38 | 1.25E-05 | 4.05 | 5.28E-05 |
| 160 | SMN-VIN | 49.31 | <0.001 | -6.95 | 4.44E-12 | -9.79 | 2.42E-22 | -3.36 | 7.83E-04 |
| 161 | VAN-VAN | 24.58 | <0.001 | 0.96 | 3.37E-01 | 6.25 | 4.69E-10 | 5.83 | 5.91E-09 |
| 162 | VAN-VIN | 87.36 | <0.001 | -6.23 | 5.15E-10 | -13.13 | 2.10E-38 | -7.78 | 9.72E-15 |
| 163 | VIN-VIN | 48.86 | <0.001 | 2.95 | 3.25E-03 | 9.43 | 7.73E-21 | 7.21 | 6.78E-13 |
| 164 | AUN-tha.Lh | 0.39 | 0.67 | -0.85 | 0.39 | -0.29 | 0.77 | 0.59 | 0.56 |
| 165 | AUN-cau.Lh | 49.93 | <0.001 | -7.29 | 3.95E-13 | -9.78 | 2.87E-22 | -2.98 | 2.88E-03 |
| 166 | AUN-put.Lh | 176.15 | <0.001 | -10.07 | 1.60E-23 | -18.75 | 1.26E-74 | -9.87 | 1.13E-22 |
| 167 | AUN-pal.Lh | 2.49 | 0.08 | 1.05 | 0.29 | 2.22 | 0.03 | 1.31 | 0.19 |
| 168 | AUN-hip.Lh | 40.59 | <0.001 | -3.36 | 7.91E-04 | -8.78 | 2.58E-18 | -6.06 | 1.48E-09 |
| 169 | AUN-amy.Lh | 4.21 | 0.01 | -0.97 | 0.33 | -2.8 | 0.005 | -2.04 | 0.04 |
| 170 | AUN-acc.Lh | 243.68 | <0.001 | 15.1 | 7.78E-50 | 21.85 | 6.76E-99 | 7.93 | 3.03E-15 |
| 171 | AUN-vdc.Lh | 147.41 | <0.001 | 11.57 | 2.37E-30 | 17.02 | 2.38E-62 | 6.39 | 1.90E-10 |
| 172 | AUN-b.s. | 146.21 | <0.001 | -11.35 | 2.71E-29 | -16.98 | 4.54E-62 | -6.58 | 5.55E-11 |
| 173 | AUN-tha.Rh | 11.12 | <0.001 | -0.45 | 6.54E-01 | -4.11 | 4.08E-05 | -4.03 | 5.66E-05 |
| 174 | AUN-cau.Rh | 106.88 | <0.001 | 10.28 | 2.01E-24 | 14.41 | 1.13E-45 | 4.88 | 1.10E-06 |
| 175 | AUN-put.Rh | 516.1 | <0.001 | 17.28 | 4.05E-64 | 32.1 | 6.66E-197 | 16.86 | 3.26E-61 |
| 176 | AUN-pal.Rh | 109.09 | <0.001 | 7.2 | 7.69E-13 | 14.7 | 2.06E-47 | 8.48 | 3.35E-17 |
| 177 | AUN-hip.Rh | 84.02 | <0.001 | -8.9 | 8.92E-19 | -12.82 | 9.41E-37 | -4.61 | 4.22E-06 |
| 178 | AUN-amy.Rh | 15.36 | <0.001 | -5.33 | 1.04E-07 | -4.29 | 1.84E-05 | 0.96 | 3.38E-01 |
| 179 | AUN-acc.Rh | 1.28 | 0.28 | 1.47 | 0.14 | 0.33 | 0.74 | -1.2 | 0.23 |
| 180 | AUN-vdc.Rh | 67.82 | <0.001 | -8.76 | 2.99E-18 | -11.3 | 4.33E-29 | -3.09 | 2.02E-03 |
| 181 | CON-tha.Lh | 9.57 | <0.001 | 3.58 | 3.50E-04 | 4.11 | 4.05E-05 | 0.71 | 4.80E-01 |
| 182 | CON-cau.Lh | 315.78 | <0.001 | 14.4 | 1.17E-45 | 25.13 | 1.26E-127 | 12.27 | 7.41E-34 |
| 183 | CON-put.Lh | 275.02 | <0.001 | 13.4 | 6.87E-40 | 23.45 | 1.35E-112 | 11.5 | 5.15E-30 |
| 184 | CON-pal.Lh | 8.84 | <0.001 | -4.06 | 5.08E-05 | -3.23 | 1.26E-03 | 0.77 | 4.41E-01 |
| 185 | CON-hip.Lh | 20.21 | <0.001 | 6.36 | 2.34E-10 | 3.56 | 3.73E-04 | -2.85 | 4.46E-03 |
| 186 | CON-amy.Lh | 676.54 | <0.001 | 20.15 | 3.38E-85 | 36.77 | 3.40E-248 | 18.94 | 5.65E-76 |
| 187 | CON-acc.Lh | 149.97 | <0.001 | 8.15 | 4.92E-16 | 17.2 | 1.54E-63 | 10.2 | 4.36E-24 |
| 188 | CON-vdc.Lh | 82.97 | <0.001 | -8.45 | 4.29E-17 | -12.81 | 1.10E-36 | -5.07 | 4.13E-07 |
| 189 | CON-b.s. | 242.84 | <0.001 | 11.55 | 2.69E-30 | 22 | 3.32E-100 | 11.86 | 8.11E-32 |
| 190 | CON-tha.Rh | 0.72 | 0.49 | 0.72 | 0.47 | -0.38 | 0.71 | -1.18 | 0.24 |
| 191 | CON-cau.Rh | 24.86 | <0.001 | -6.16 | 8.19E-10 | -6.34 | 2.64E-10 | -0.41 | 6.82E-01 |
| 192 | CON-put.Rh | 51.01 | <0.001 | 3.68 | 2.33E-04 | 9.82 | 1.82E-22 | 6.86 | 8.07E-12 |
| 193 | CON-pal.Rh | 0.34 | 0.71 | -0.34 | 0.73 | -0.81 | 0.42 | -0.53 | 0.6 |
| 194 | CON-hip.Rh | 471.08 | <0.001 | 19.47 | 5.85E-80 | 30.6 | 3.75E-181 | 12.89 | 4.02E-37 |
| 195 | CON-amy.Rh | 324.47 | <0.001 | 15 | 3.07E-49 | 25.47 | 9.79E-131 | 12.01 | 1.58E-32 |
| 196 | CON-acc.Rh | 5.59 | <0.001 | -1.4 | 0.16 | 1.69 | 0.09 | 3.34 | 0.0008 |
| 197 | CON-vdc.Rh | 242.99 | <0.001 | 10.52 | 1.83E-25 | 21.91 | 2.01E-99 | 12.86 | 5.59E-37 |
| 198 | CPN-tha.Lh | 466.6 | <0.001 | 17.12 | 5.52E-63 | 30.55 | 1.37E-180 | 15.32 | 2.97E-51 |
| 199 | CPN-cau.Lh | 87.72 | <0.001 | 6.07 | 1.46E-09 | 13.13 | 2.04E-38 | 7.96 | 2.38E-15 |
| 200 | CPN-put.Lh | 30.05 | <0.001 | -4.68 | 2.92E-06 | -7.75 | 1.26E-14 | -3.52 | 4.37E-04 |
| 201 | CPN-pal.Lh | 41.7 | <0.001 | 3.63 | 2.83E-04 | 8.95 | 5.72E-19 | 5.96 | 2.79E-09 |
| 202 | CPN-hip.Lh | 7.68 | <0.001 | 0.63 | 0.53 | 3.53 | 0.0004 | 3.21 | 0.001 |
| 203 | CPN-amy.Lh | 52.7 | <0.001 | 5.35 | 9.45E-08 | 10.25 | 2.80E-24 | 5.56 | 2.92E-08 |
| 204 | CPN-acc.Lh | 2.54 | 0.08 | -0.06 | 0.95 | -1.88 | 0.06 | -2 | 0.05 |
| 205 | CPN-vdc.Lh | 186.44 | <0.001 | 11.86 | 8.20E-32 | 19.28 | 1.41E-78 | 8.55 | 1.83E-17 |
| 206 | CPN-b.s. | 4.76 | 0.01 | -0.91 | 0.36 | -2.94 | 0.003 | -2.25 | 0.02 |
| 207 | CPN-tha.Rh | 50.3 | <0.001 | 3.33 | 8.86E-04 | 9.67 | 8.18E-22 | 7.07 | 1.90E-12 |
| 208 | CPN-cau.Rh | 361.76 | <0.001 | 12.98 | 1.38E-37 | 26.75 | 8.68E-143 | 15.56 | 9.79E-53 |
| 209 | CPN-put.Rh | 39.62 | <0.001 | -2.62 | 8.82E-03 | -8.48 | 3.38E-17 | -6.52 | 8.23E-11 |
| 210 | CPN-pal.Rh | 345.61 | <0.001 | 14.26 | 8.37E-45 | 26.27 | 2.85E-138 | 13.68 | 1.87E-41 |
| 211 | CPN-hip.Rh | 72.54 | <0.001 | 4.53 | 5.97E-06 | 11.75 | 3.00E-31 | 8.07 | 9.66E-16 |
| 212 | CPN-amy.Rh | 27.47 | <0.001 | -5.32 | 1.09E-07 | -7.27 | 4.33E-13 | -2.33 | 2.01E-02 |
| 213 | CPN-acc.Rh | 0.89 | 4.10E-01 | -0.73 | 0.46 | -1.33 | 0.18 | -0.68 | 0.49 |
| 214 | CPN-vdc.Rh | 0.07 | 0.93 | -0.32 | 0.75 | -0.33 | 0.74 | -0.03 | 0.98 |
| 215 | DMN-tha.Lh | 2.91 | 0.05 | 1.27 | 0.2 | 2.41 | 0.02 | 1.29 | 0.2 |
| 216 | DMN-cau.Lh | 5.36 | <0.001 | -1.54 | 0.12 | -3.25 | 0.001 | -1.93 | 0.05 |
| 217 | DMN-put.Lh | 212.71 | <0.001 | 11.36 | 2.20E-29 | 20.62 | 6.27E-89 | 10.55 | 1.35E-25 |
| 218 | DMN-pal.Lh | 199.77 | <0.001 | 10.69 | 3.20E-26 | 19.97 | 8.02E-84 | 10.55 | 1.24E-25 |
| 219 | DMN-hip.Lh | 137.05 | <0.001 | 8.72 | 4.25E-18 | 16.53 | 4.90E-59 | 8.87 | 1.18E-18 |
| 220 | DMN-amy.Lh | 3.78 | 0.02 | 0.33 | 0.74 | -2.05 | 0.04 | -2.6 | 0.009 |
| 221 | DMN-acc.Lh | 134.71 | <0.001 | 9.65 | 9.54E-22 | 16.41 | 3.13E-58 | 7.75 | 1.20E-14 |
| 222 | DMN-vdc.Lh | 12.96 | <0.001 | 2.27 | 2.34E-02 | 5.04 | 4.98E-07 | 3.12 | 1.85E-03 |
| 223 | DMN-b.s. | 3.18 | 0.04 | -0.96 | 0.34 | 1.37 | 0.17 | 2.52 | 0.01 |
| 224 | DMN-tha.Rh | 19.51 | <0.001 | -3.38 | 7.26E-04 | -6.24 | 4.84E-10 | -3.26 | 1.14E-03 |
| 225 | DMN-cau.Rh | 1.66 | 0.19 | 1.13 | 0.26 | 1.82 | 0.07 | 0.8 | 0.42 |
| 226 | DMN-put.Rh | 33.68 | <0.001 | -5.27 | 1.49E-07 | -8.18 | 4.15E-16 | -3.37 | 7.47E-04 |
| 227 | DMN-pal.Rh | 0.57 | 5.60E-01 | 0.09 | 0.93 | -0.83 | 0.41 | -1 | 0.32 |
| 228 | DMN-hip.Rh | 0.12 | 0.89 | 0.48 | 0.63 | 0.33 | 0.74 | -0.15 | 0.88 |
| 229 | DMN-amy.Rh | 133.49 | <0.001 | 9.81 | 2.06E-22 | 16.33 | 1.08E-57 | 7.49 | 8.68E-14 |
| 230 | DMN-acc.Rh | 89.77 | <0.001 | 7.11 | 1.38E-12 | 13.38 | 8.20E-40 | 7.12 | 1.28E-12 |
| 231 | DMN-vdc.Rh | 18.27 | <0.001 | -3.52 | 4.42E-04 | -6.04 | 1.67E-09 | -2.89 | 3.83E-03 |
| 232 | DAN-tha.Lh | 12.35 | <0.001 | 4.34 | 1.44E-05 | 4.46 | 8.31E-06 | 0.28 | 7.78E-01 |
| 233 | DAN-cau.Lh | 12.81 | <0.001 | -4.91 | 9.48E-07 | -3.81 | 1.39E-04 | 1.03 | 3.01E-01 |
| 234 | DAN-put.Lh | 18.17 | <0.001 | -3.64 | 2.80E-04 | -6.02 | 1.90E-09 | -2.74 | 6.11E-03 |
| 235 | DAN-pal.Lh | 4.93 | 0.01 | 1.6 | 0.11 | 3.13 | 0.002 | 1.74 | 0.08 |
| 236 | DAN-hip.Lh | 45.03 | <0.001 | -3.99 | 6.80E-05 | -9.35 | 1.64E-20 | -6.02 | 1.99E-09 |
| 237 | DAN-amy.Lh | 5.36 | <0.001 | 1.21 | 0.23 | 3.19 | 0.001 | 2.21 | 0.03 |
| 238 | DAN-acc.Lh | 63 | <0.001 | 3.87 | 1.12E-04 | 10.86 | 5.26E-27 | 7.8 | 8.15E-15 |
| 239 | DAN-vdc.Lh | 15.03 | <0.001 | 3.05 | 2.29E-03 | 5.48 | 4.51E-08 | 2.77 | 5.61E-03 |
| 240 | DAN-b.s. | 113.88 | <0.001 | 6.48 | 1.03E-10 | 14.89 | 1.46E-48 | 9.45 | 6.49E-21 |
| 241 | DAN-tha.Rh | 58.06 | <0.001 | -6.65 | 3.35E-11 | -10.76 | 1.49E-26 | -4.73 | 2.29E-06 |
| 242 | DAN-cau.Rh | 17.3 | <0.001 | 3.08 | 2.11E-03 | 5.87 | 4.74E-09 | 3.17 | 1.52E-03 |
| 243 | DAN-put.Rh | 13.07 | <0.001 | 0.38 | 7.02E-01 | 4.4 | 1.11E-05 | 4.42 | 1.00E-05 |
| 244 | DAN-pal.Rh | 40.99 | <0.001 | -4.15 | 3.44E-05 | -8.98 | 4.65E-19 | -5.44 | 5.74E-08 |
| 245 | DAN-hip.Rh | 15.93 | <0.001 | -5.13 | 3.03E-07 | -4.86 | 1.22E-06 | 0.12 | 9.04E-01 |
| 246 | DAN-amy.Rh | 16.07 | <0.001 | -3.69 | 2.32E-04 | -5.64 | 1.83E-08 | -2.27 | 2.31E-02 |
| 247 | DAN-acc.Rh | 25.09 | <0.001 | 4.53 | 6.03E-06 | 7.06 | 2.06E-12 | 2.93 | 3.45E-03 |
| 248 | DAN-vdc.Rh | 114.31 | <0.001 | 7.36 | 2.29E-13 | 15.04 | 1.61E-49 | 8.68 | 5.92E-18 |
| 249 | FPN-tha.Lh | 9.44 | <0.001 | 2.63 | 8.55E-03 | 4.34 | 1.46E-05 | 1.97 | 4.93E-02 |
| 250 | FPN-cau.Lh | 85.06 | <0.001 | 6.51 | 8.39E-11 | 13 | 1.09E-37 | 7.34 | 2.77E-13 |
| 251 | FPN-put.Lh | 31.25 | <0.001 | 4.78 | 1.83E-06 | 7.9 | 3.80E-15 | 3.59 | 3.38E-04 |
| 252 | FPN-pal.Lh | 67.48 | <0.001 | -8.71 | 4.86E-18 | -11.29 | 5.15E-29 | -3.13 | 1.75E-03 |
| 253 | FPN-hip.Lh | 32.41 | <0.001 | 4.66 | 3.30E-06 | 8.05 | 1.13E-15 | 3.88 | 1.05E-04 |
| 254 | FPN-amy.Lh | 8.13 | <0.001 | -0.13 | 0.9 | 3.23 | 0.001 | 3.68 | 0.0002 |
| 255 | FPN-acc.Lh | 57.15 | <0.001 | -4.22 | 2.50E-05 | -10.47 | 2.87E-25 | -7 | 2.99E-12 |
| 256 | FPN-vdc.Lh | 72.79 | <0.001 | 7.14 | 1.15E-12 | 12.06 | 8.23E-33 | 5.65 | 1.76E-08 |
| 257 | FPN-b.s. | 61.76 | <0.001 | -6.31 | 3.21E-10 | -11.11 | 3.40E-28 | -5.49 | 4.30E-08 |
| 258 | FPN-tha.Rh | 22.21 | <0.001 | -4.01 | 6.19E-05 | -6.66 | 3.21E-11 | -3.04 | 2.35E-03 |
| 259 | FPN-cau.Rh | 23.85 | <0.001 | 2.07 | 3.89E-02 | 6.59 | 5.16E-11 | 5.03 | 5.09E-07 |
| 260 | FPN-put.Rh | 17.75 | <0.001 | -1.39 | 1.65E-01 | -5.55 | 3.12E-08 | -4.61 | 4.15E-06 |
| 261 | FPN-pal.Rh | 17.95 | <0.001 | -4.47 | 8.06E-06 | -5.83 | 6.16E-09 | -1.64 | 1.00E-01 |
| 262 | FPN-hip.Rh | 57.55 | <0.001 | 4.49 | 7.50E-06 | 10.56 | 1.16E-25 | 6.82 | 1.08E-11 |
| 263 | FPN-amy.Rh | 6.57 | <0.001 | 1.54 | 0.12 | 3.57 | 0.0004 | 2.28 | 0.02 |
| 264 | FPN-acc.Rh | 70.25 | <0.001 | 7.84 | 6.08E-15 | 11.78 | 2.19E-31 | 4.59 | 4.58E-06 |
| 265 | FPN-vdc.Rh | 20.55 | <0.001 | 4.46 | 8.31E-06 | 6.33 | 2.83E-10 | 2.2 | 2.80E-02 |
| 266 | RTN-tha.Lh | 0.22 | 0.8 | 0.46 | 0.65 | 0.65 | 0.51 | 0.23 | 0.82 |
| 267 | RTN-cau.Lh | 130.3 | <0.001 | -10.45 | 3.80E-25 | -16.07 | 5.36E-56 | -6.53 | 7.47E-11 |
| 268 | RTN-put.Lh | 27.43 | <0.001 | -3.34 | 8.58E-04 | -7.33 | 2.81E-13 | -4.5 | 7.02E-06 |
| 269 | RTN-pal.Lh | 1.74 | 0.18 | -0.61 | 0.54 | -1.79 | 0.07 | -1.32 | 0.19 |
| 270 | RTN-hip.Lh | 118.28 | <0.001 | 10.53 | 1.54E-25 | 15.22 | 1.40E-50 | 5.5 | 4.02E-08 |
| 271 | RTN-amy.Lh | 180.52 | <0.001 | -11.18 | 1.75E-28 | -19 | 1.95E-76 | -8.97 | 4.94E-19 |
| 272 | RTN-acc.Lh | 29.14 | <0.001 | -1.94 | 5.20E-02 | -7.17 | 9.31E-13 | -5.8 | 7.27E-09 |
| 273 | RTN-vdc.Lh | 80.87 | <0.001 | 8.76 | 3.06E-18 | 12.57 | 1.95E-35 | 4.48 | 7.55E-06 |
| 274 | RTN-b.s. | 217.19 | <0.001 | 15.01 | 2.74E-49 | 20.44 | 1.51E-87 | 6.49 | 1.02E-10 |
| 275 | RTN-tha.Rh | 99.36 | <0.001 | 6.49 | 1.02E-10 | 13.98 | 3.61E-43 | 8.44 | 4.55E-17 |
| 276 | RTN-cau.Rh | 92.36 | <0.001 | -9.13 | 1.18E-19 | -13.48 | 2.38E-40 | -5.09 | 3.79E-07 |
| 277 | RTN-put.Rh | 18.09 | <0.001 | -4.89 | 1.08E-06 | -5.67 | 1.54E-08 | -1.03 | 3.03E-01 |
| 278 | RTN-pal.Rh | 4.76 | 0.01 | 1.87 | 0.06 | 3.08 | 0.002 | 1.4 | 0.16 |
| 279 | RTN-hip.Rh | 70.76 | <0.001 | -9.08 | 1.88E-19 | -11.5 | 5.06E-30 | -2.97 | 3.00E-03 |
| 280 | RTN-amy.Rh | 4.38 | 0.01 | 0.73 | 0.47 | 2.77 | 0.006 | 2.27 | 0.02 |
| 281 | RTN-acc.Rh | 11.15 | <0.001 | 2.95 | 3.18E-03 | 4.71 | 2.56E-06 | 2.03 | 4.22E-02 |
| 282 | RTN-vdc.Rh | 291.6 | <0.001 | 14.36 | 2.08E-45 | 24.14 | 1.17E-118 | 11.22 | 1.02E-28 |
| 283 | SAN-tha.Lh | 175.06 | <0.001 | 10.11 | 1.13E-23 | 18.7 | 3.21E-74 | 9.77 | 2.90E-22 |
| 284 | SAN-cau.Lh | 0.09 | 0.91 | 0.09 | 0.93 | 0.4 | 0.69 | 0.34 | 0.73 |
| 285 | SAN-put.Lh | 153.26 | <0.001 | 9.56 | 2.23E-21 | 17.5 | 1.25E-65 | 9.04 | 2.58E-19 |
| 286 | SAN-pal.Lh | 1.07 | 0.34 | 0.52 | 0.6 | -0.82 | 0.41 | -1.45 | 0.15 |
| 287 | SAN-hip.Lh | 26.19 | <0.001 | -3.94 | 8.18E-05 | -7.23 | 5.84E-13 | -3.75 | 1.83E-04 |
| 288 | SAN-amy.Lh | 7.89 | <0.001 | 2.17 | 3.02E-02 | 3.97 | 7.34E-05 | 2.05 | 4.02E-02 |
| 289 | SAN-acc.Lh | 144.37 | <0.001 | 8.09 | 8.41E-16 | 16.89 | 2.08E-61 | 9.93 | 6.39E-23 |
| 290 | SAN-vdc.Lh | 47.55 | <0.001 | -3.8 | 1.46E-04 | -9.54 | 2.63E-21 | -6.43 | 1.47E-10 |
| 291 | SAN-b.s. | 22.51 | <0.001 | -5.89 | 4.23E-09 | -6 | 2.15E-09 | -0.33 | 7.44E-01 |
| 292 | SAN-tha.Rh | 64.24 | <0.001 | 4.82 | 1.49E-06 | 11.17 | 1.79E-28 | 7.14 | 1.18E-12 |
| 293 | SAN-cau.Rh | 18.98 | <0.001 | 2.97 | 3.02E-03 | 6.13 | 1.00E-09 | 3.57 | 3.64E-04 |
| 294 | SAN-put.Rh | 44.53 | <0.001 | -5.48 | 4.55E-08 | -9.44 | 7.04E-21 | -4.53 | 6.12E-06 |
| 295 | SAN-pal.Rh | 74.72 | <0.001 | -7.67 | 2.32E-14 | -12.2 | 1.73E-33 | -5.23 | 1.76E-07 |
| 296 | SAN-hip.Rh | 27.87 | <0.001 | 4.44 | 9.26E-06 | 7.46 | 1.08E-13 | 3.47 | 5.28E-04 |
| 297 | SAN-amy.Rh | 17.96 | <0.001 | 1.53 | 1.26E-01 | 5.63 | 1.96E-08 | 4.55 | 5.55E-06 |
| 298 | SAN-acc.Rh | 45.27 | <0.001 | -5.07 | 4.15E-07 | -9.51 | 3.72E-21 | -5.04 | 4.93E-07 |
| 299 | SAN-vdc.Rh | 45.87 | <0.001 | -8.13 | 5.84E-16 | -8.8 | 2.17E-18 | -1.01 | 3.11E-01 |
| 300 | SHN-tha.Lh | 18.33 | <0.001 | -5.36 | 8.91E-08 | -5.37 | 8.23E-08 | -0.2 | 8.40E-01 |
| 301 | SHN-cau.Lh | 237.16 | <0.001 | 11.71 | 4.42E-31 | 21.76 | 3.40E-98 | 11.43 | 1.10E-29 |
| 302 | SHN-put.Lh | 23.87 | <0.001 | 6.49 | 9.90E-11 | 5.66 | 1.68E-08 | -0.69 | 4.90E-01 |
| 303 | SHN-pal.Lh | 661.78 | <0.001 | 20.4 | 3.61E-87 | 36.38 | 8.95E-244 | 18.24 | 7.13E-71 |
| 304 | SHN-hip.Lh | 97.66 | <0.001 | -8.78 | 2.60E-18 | -13.94 | 5.85E-43 | -5.97 | 2.68E-09 |
| 305 | SHN-amy.Lh | 3.81 | 2.00E-02 | -2.21 | 0.03 | 0.09 | 0.93 | 2.45 | 0.01 |
| 306 | SHN-acc.Lh | 0.06 | 0.94 | -0.31 | 0.75 | -0.06 | 0.95 | 0.26 | 0.79 |
| 307 | SHN-vdc.Lh | 28.52 | <0.001 | -6.96 | 3.99E-12 | -6.38 | 2.02E-10 | 0.4 | 6.90E-01 |
| 308 | SHN-b.s. | 140.79 | <0.001 | 7.74 | 1.34E-14 | 16.64 | 9.25E-60 | 10.04 | 2.31E-23 |
| 309 | SHN-tha.Rh | 1.26 | 0.28 | -0.59 | 0.55 | -1.54 | 0.12 | -1.07 | 0.29 |
| 310 | SHN-cau.Rh | 478.58 | <0.001 | 18.94 | 4.83E-76 | 30.9 | 3.01E-184 | 13.77 | 5.51E-42 |
| 311 | SHN-put.Rh | 329.9 | <0.001 | 15.39 | 1.15E-51 | 25.67 | 1.28E-132 | 11.81 | 1.46E-31 |
| 312 | SHN-pal.Rh | 3.94 | 0.02 | -1.84 | 0.07 | 0.69 | 0.49 | 2.71 | 0.007 |
| 313 | SHN-hip.Rh | 294.11 | <0.001 | 12.19 | 1.84E-33 | 24.17 | 5.97E-119 | 13.57 | 7.65E-41 |
| 314 | SHN-amy.Rh | 7.48 | <0.001 | 3.16 | 0.002 | -0.03 | 0.98 | -3.39 | 0.0007 |
| 315 | SHN-acc.Rh | 436.98 | <0.001 | 17.28 | 3.87E-64 | 29.56 | 1.76E-170 | 14.06 | 1.14E-43 |
| 316 | SHN-vdc.Rh | 88.85 | <0.001 | 7.38 | 2.05E-13 | 13.33 | 1.68E-39 | 6.78 | 1.38E-11 |
| 317 | SMN-tha.Lh | 35.25 | <0.001 | 2.83 | 4.70E-03 | 8.1 | 7.42E-16 | 5.89 | 4.37E-09 |
| 318 | SMN-cau.Lh | 4.84 | 0.01 | -0.3 | 0.76 | -2.71 | 0.007 | -2.66 | 0.008 |
| 319 | SMN-put.Lh | 4.93 | 0.01 | -0.77 | 0.44 | 2.05 | 0.04 | 3.08 | 0.002 |
| 320 | SMN-pal.Lh | 44.23 | <0.001 | 4.38 | 1.23E-05 | 9.33 | 1.84E-20 | 5.59 | 2.52E-08 |
| 321 | SMN-hip.Lh | 168.41 | <0.001 | 11.89 | 6.25E-32 | 18.27 | 4.41E-71 | 7.41 | 1.57E-13 |
| 322 | SMN-amy.Lh | 130.51 | <0.001 | 10.73 | 2.06E-26 | 16.04 | 7.89E-56 | 6.2 | 6.17E-10 |
| 323 | SMN-acc.Lh | 41.89 | <0.001 | 3.54 | 4.13E-04 | 8.95 | 5.89E-19 | 6.06 | 1.50E-09 |
| 324 | SMN-vdc.Lh | 380.24 | <0.001 | 13.13 | 2.12E-38 | 27.4 | 4.50E-149 | 16.12 | 2.63E-56 |
| 325 | SMN-b.s. | 0.84 | 0.43 | 0.02 | 0.98 | -1.05 | 0.29 | -1.17 | 0.24 |
| 326 | SMN-tha.Rh | 300.29 | <0.001 | 13.53 | 1.30E-40 | 24.5 | 6.92E-122 | 12.51 | 4.27E-35 |
| 327 | SMN-cau.Rh | 66.23 | <0.001 | 4.97 | 6.90E-07 | 11.36 | 2.34E-29 | 7.18 | 8.75E-13 |
| 328 | SMN-put.Rh | 36.24 | <0.001 | -5.35 | 9.62E-08 | -8.49 | 3.04E-17 | -3.64 | 2.80E-04 |
| 329 | SMN-pal.Rh | 1.65 | 1.90E-01 | 0 | 0.99 | -1.49 | 0.14 | -1.63 | 0.1 |
| 330 | SMN-hip.Rh | 0.17 | 0.85 | -0.09 | 0.92 | 0.41 | 0.68 | 0.55 | 0.58 |
| 331 | SMN-amy.Rh | 8.75 | <0.001 | -2.48 | 1.31E-02 | -4.18 | 2.98E-05 | -1.95 | 5.13E-02 |
| 332 | SMN-acc.Rh | 0.5 | 0.6 | 0.88 | 0.38 | 0.1 | 0.92 | -0.82 | 0.41 |
| 333 | SMN-vdc.Rh | 5.65 | <0.001 | -0.84 | 0.4 | -3.15 | 0.002 | -2.57 | 0.01 |
| 334 | VAN-tha.Lh | 29.08 | <0.001 | 4.57 | 5.06E-06 | 7.62 | 3.26E-14 | 3.51 | 4.61E-04 |
| 335 | VAN-cau.Lh | 136.99 | <0.001 | 8.14 | 5.64E-16 | 16.48 | 1.11E-58 | 9.43 | 7.34E-21 |
| 336 | VAN-put.Lh | 45.73 | <0.001 | -4.3 | 1.73E-05 | -9.47 | 5.27E-21 | -5.81 | 6.67E-09 |
| 337 | VAN-pal.Lh | 15.11 | <0.001 | 3.38 | 7.46E-04 | 5.49 | 4.33E-08 | 2.44 | 1.49E-02 |
| 338 | VAN-hip.Lh | 35.36 | <0.001 | 3.54 | 4.00E-04 | 8.28 | 1.72E-16 | 5.32 | 1.09E-07 |
| 339 | VAN-amy.Lh | 99.26 | <0.001 | -10.21 | 3.95E-24 | -13.8 | 3.65E-42 | -4.29 | 1.83E-05 |
| 340 | VAN-acc.Lh | 102.63 | <0.001 | -9.21 | 5.81E-20 | -14.27 | 7.23E-45 | -5.87 | 4.71E-09 |
| 341 | VAN-vdc.Lh | 41.32 | <0.001 | 4.92 | 9.15E-07 | 9.08 | 1.75E-19 | 4.74 | 2.21E-06 |
| 342 | VAN-b.s. | 79.23 | <0.001 | 6.47 | 1.11E-10 | 12.56 | 2.29E-35 | 6.9 | 6.16E-12 |
| 343 | VAN-tha.Rh | 69.59 | <0.001 | -5.86 | 5.03E-09 | -11.75 | 2.91E-31 | -6.66 | 3.10E-11 |
| 344 | VAN-cau.Rh | 58.03 | <0.001 | 5.67 | 1.58E-08 | 10.76 | 1.51E-26 | 5.78 | 8.13E-09 |
| 345 | VAN-put.Rh | 1.5 | 0.22 | 1.29 | 0.2 | 1.69 | 0.09 | 0.48 | 0.63 |
| 346 | VAN-pal.Rh | 16.32 | <0.001 | -4.43 | 9.71E-06 | -5.49 | 4.27E-08 | -1.32 | 1.88E-01 |
| 347 | VAN-hip.Rh | 25.9 | <0.001 | 2.38 | 1.74E-02 | 6.93 | 4.89E-12 | 5.08 | 4.01E-07 |
| 348 | VAN-amy.Rh | 13.78 | <0.001 | -1.44 | 1.49E-01 | -4.97 | 7.14E-07 | -3.92 | 9.17E-05 |
| 349 | VAN-acc.Rh | 17.34 | <0.001 | -4.65 | 3.43E-06 | -5.62 | 2.05E-08 | -1.23 | 2.20E-01 |
| 350 | VAN-vdc.Rh | 64.24 | <0.001 | 4.82 | 1.49E-06 | 11.17 | 1.79E-28 | 7.14 | 1.18E-12 |
| 351 | VIN-tha.Lh | 64.66 | <0.001 | 7.38 | 1.93E-13 | 11.32 | 3.71E-29 | 4.57 | 5.06E-06 |
| 352 | VIN-cau.Lh | 14.72 | <0.001 | 3.85 | 1.21E-04 | 5.34 | 1.00E-07 | 1.77 | 7.73E-02 |
| 353 | VIN-put.Lh | 0.66 | 0.52 | 0.28 | 0.78 | 1.08 | 0.28 | 0.88 | 0.38 |
| 354 | VIN-pal.Lh | 50.94 | <0.001 | 6.31 | 3.20E-10 | 10.07 | 1.62E-23 | 4.35 | 1.43E-05 |
| 355 | VIN-hip.Lh | 239.78 | <0.001 | 12.79 | 1.41E-36 | 21.9 | 2.58E-99 | 10.44 | 4.19E-25 |
| 356 | VIN-amy.Lh | 11.37 | <0.001 | 1.42 | 1.57E-01 | 4.55 | 5.64E-06 | 3.48 | 5.03E-04 |
| 357 | VIN-acc.Lh | 47.93 | <0.001 | -7.8 | 8.51E-15 | -9.32 | 2.14E-20 | -1.94 | 5.27E-02 |
| 358 | VIN-vdc.Lh | 8.15 | <0.001 | -3.76 | 0.0002 | -3.36 | 0.0008 | 0.31 | 0.76 |
| 359 | VIN-b.s. | 6.49 | <0.001 | 0.59 | 0.55 | -2.58 | 0.01 | -3.46 | 0.0006 |
| 360 | VIN-tha.Rh | 61.7 | <0.001 | -8 | 1.69E-15 | -10.9 | 3.49E-27 | -3.45 | 5.63E-04 |
| 361 | VIN-cau.Rh | 17.01 | <0.001 | 2.91 | 3.63E-03 | 5.81 | 6.80E-09 | 3.28 | 1.04E-03 |
| 362 | VIN-put.Rh | 15.38 | <0.001 | 2.24 | 2.50E-02 | 5.44 | 5.58E-08 | 3.59 | 3.34E-04 |
| 363 | VIN-pal.Rh | 206.11 | <0.001 | 12.76 | 2.03E-36 | 20.25 | 4.95E-86 | 8.66 | 7.07E-18 |
| 364 | VIN-hip.Rh | 183.97 | <0.001 | 11.98 | 2.24E-32 | 19.14 | 1.67E-77 | 8.28 | 1.84E-16 |
| 365 | VIN-amy.Rh | 25.29 | <0.001 | -5.55 | 3.11E-08 | -6.82 | 1.06E-11 | -1.59 | 1.12E-01 |
| 366 | VIN-acc.Rh | 78.24 | <0.001 | 7.31 | 3.32E-13 | 12.51 | 4.22E-35 | 5.95 | 2.88E-09 |
| 367 | VIN-vdc.Rh | 28.57 | <0.001 | 6.56 | 6.17E-11 | 6.83 | 1.01E-11 | 0.52 | 6.02E-01 |

### **Table S8. Longitudinal changes of cognitive performances across three adolescent biotypes**

| Cognitive performance | F value | *P* value | biotype1  (N) | biotype2  (N) | biotype3  (N) | Total number |
| --- | --- | --- | --- | --- | --- | --- |
| picture vocabulary | 1.74 | 0.14 | 564 | 558 | 620 | 1742 |
| flanker | 0.33 | 0.86 | 472 | 465 | 510 | 1448 |
| pattern | 1.25 | 0.29 | 468 | 465 | 506 | 1439 |
| picture | 2.03 | 0.09 | 570 | 571 | 624 | 1765 |
| reading | 0.94 | 0.44 | 555 | 554 | 616 | 1725 |
| crystalized intelligence | 1.43 | 0.22 | 98 | 103 | 104 | 305 |

### **Table S9. Longitudinal development of brain volumes and network functional connectivitites across three adolescent biotypes**

| ID | brain measures | F value | *P* value | biotype 1  (N) | biotype 2  (N) | biotype 3  (N) | total number |
| --- | --- | --- | --- | --- | --- | --- | --- |
| 1 | Lh_BanksOfSuperiorTemporalSulcus | 0.59 | 0.67 | 391 | 395 | 429 | 1215 |
| 2 | Lh_caudalanteriorcingulate | 0.4 | 0.81 | 391 | 395 | 429 | 1215 |
| 3 | Lh_caudalmiddlefrontal | 2.11 | 0.08 | 391 | 395 | 429 | 1215 |
| 4 | Lh_cuneus | 1.02 | 0.4 | 391 | 395 | 429 | 1215 |
| 5 | Lh_entorhinal | 1.15 | 0.33 | 391 | 395 | 429 | 1215 |
| 6 | Lh_fusiform | 0.25 | 0.91 | 391 | 395 | 429 | 1215 |
| 7 | Lh_inferiorparietal | 3.19 | 0.01 | 391 | 395 | 429 | 1215 |
| 8 | Lh_inferiortemporal | 1.64 | 0.16 | 391 | 395 | 429 | 1215 |
| 9 | Lh_isthmuscingulate | 2.12 | 0.08 | 391 | 395 | 429 | 1215 |
| 10 | Lh_lateraloccipital | 1.07 | 0.37 | 391 | 395 | 429 | 1215 |
| 11 | Lh_lateralorbitofrontal | 0.98 | 0.42 | 391 | 395 | 429 | 1215 |
| 12 | Lh_lingual | 1.08 | 0.36 | 391 | 395 | 429 | 1215 |
| 13 | Lh_medialorbitofrontal | 0.36 | 0.84 | 391 | 395 | 429 | 1215 |
| 14 | Lh_middletemporal | 3.59 | 0.01 | 391 | 395 | 429 | 1215 |
| 15 | Lh_parahippocampal | 1.01 | 0.4 | 391 | 395 | 429 | 1215 |
| 16 | Lh_paracentral | 2.27 | 0.06 | 391 | 395 | 429 | 1215 |
| 17 | Lh_parsopercularis | 0.94 | 0.44 | 391 | 395 | 429 | 1215 |
| 18 | Lh_parsorbitalis | 1.7 | 0.15 | 391 | 395 | 429 | 1215 |
| 19 | Lh_parstriangularis | 1.79 | 0.13 | 391 | 395 | 429 | 1215 |
| 20 | Lh_pericalcarine | 2.2 | 0.07 | 391 | 395 | 429 | 1215 |
| 21 | Lh_postcentral | 1.07 | 0.37 | 391 | 395 | 429 | 1215 |
| 22 | Lh_posteriorcingulate | 0.77 | 0.54 | 391 | 395 | 429 | 1215 |
| 23 | Lh_precentral | 4.43 | <0.001 | 391 | 395 | 429 | 1215 |
| 24 | Lh_precuneus | 2.15 | 0.07 | 391 | 395 | 429 | 1215 |
| 25 | Lh_rostralanteriorcingulate | 0.16 | 0.96 | 391 | 395 | 429 | 1215 |
| 26 | Lh_rostralmiddlefrontal | 0.36 | 0.83 | 391 | 395 | 429 | 1215 |
| 27 | Lh_superiorfrontal | 2.19 | 0.07 | 391 | 395 | 429 | 1215 |
| 28 | Lh_superiorparietal | 1.11 | 0.35 | 391 | 395 | 429 | 1215 |
| 29 | Lh_superiortemporal | 4.37 | <0.001 | 391 | 395 | 429 | 1215 |
| 30 | Lh_supramarginal | 0.68 | 0.61 | 391 | 395 | 429 | 1215 |
| 31 | Lh_frontalpole | 1.03 | 0.39 | 391 | 395 | 429 | 1215 |
| 32 | Lh_temporalpole | 2 | 0.09 | 391 | 395 | 429 | 1215 |
| 33 | Lh_transversetemporal | 2.54 | 0.04 | 391 | 395 | 429 | 1215 |
| 34 | Lh_insula | 1.05 | 0.38 | 391 | 395 | 429 | 1215 |
| 35 | Rh_BanksOfSuperiorTemporalSulcus | 1.4 | 0.23 | 391 | 395 | 429 | 1215 |
| 36 | Rh_caudalanteriorcingulate | 1.1 | 0.36 | 391 | 395 | 429 | 1215 |
| 37 | Rh_caudalmiddlefrontal | 1.23 | 0.3 | 391 | 395 | 429 | 1215 |
| 38 | Rh_cuneus | 0.94 | 0.44 | 391 | 395 | 429 | 1215 |
| 39 | Rh_entorhinal | 0.84 | 0.5 | 391 | 395 | 429 | 1215 |
| 40 | Rh_fusiform | 0.57 | 0.69 | 391 | 395 | 429 | 1215 |
| 41 | Rh_inferiorparietal | 1.43 | 0.22 | 391 | 395 | 429 | 1215 |
| 42 | Rh_inferiortemporal | 0.68 | 0.6 | 391 | 395 | 429 | 1215 |
| 43 | Rh_isthmuscingulate | 1.75 | 0.14 | 391 | 395 | 429 | 1215 |
| 44 | Rh_lateraloccipital | 0.39 | 0.82 | 391 | 395 | 429 | 1215 |
| 45 | Rh_lateralorbitofrontal | 0.62 | 0.65 | 391 | 395 | 429 | 1215 |
| 46 | Rh_lingual | 1.03 | 0.39 | 391 | 395 | 429 | 1215 |
| 47 | Rh_medialorbitofrontal | 0.36 | 0.84 | 391 | 395 | 429 | 1215 |
| 48 | Rh_middletemporal | 1.8 | 0.13 | 391 | 395 | 429 | 1215 |
| 49 | Rh_parahippocampal | 1.45 | 0.22 | 391 | 395 | 429 | 1215 |
| 50 | Rh_paracentral | 1.57 | 0.18 | 391 | 395 | 429 | 1215 |
| 51 | Rh_parsopercularis | 0.22 | 0.93 | 391 | 395 | 429 | 1215 |
| 52 | Rh_parsorbitalis | 1.46 | 0.21 | 391 | 395 | 429 | 1215 |
| 53 | Rh_parstriangularis | 1.91 | 0.11 | 391 | 395 | 429 | 1215 |
| 54 | Rh_pericalcarine | 1.21 | 0.3 | 391 | 395 | 429 | 1215 |
| 55 | Rh_postcentral | 1.14 | 0.34 | 391 | 395 | 429 | 1215 |
| 56 | Rh_posteriorcingulate | 0.35 | 0.85 | 391 | 395 | 429 | 1215 |
| 57 | Rh_precentral | 2.27 | 0.06 | 391 | 395 | 429 | 1215 |
| 58 | Rh_precuneus | 1.09 | 0.36 | 391 | 395 | 429 | 1215 |
| 59 | Rh_rostralanteriorcingulate | 2.15 | 0.07 | 391 | 395 | 429 | 1215 |
| 60 | Rh_rostralmiddlefrontal | 0.39 | 0.82 | 391 | 395 | 429 | 1215 |
| 61 | Rh_superiorfrontal | 1.14 | 0.34 | 391 | 395 | 429 | 1215 |
| 62 | Rh_superiorparietal | 1.2 | 0.31 | 391 | 395 | 429 | 1215 |
| 63 | Rh_superiortemporal | 3.58 | 0.01 | 391 | 395 | 429 | 1215 |
| 64 | Rh_supramarginal | 0.15 | 0.96 | 391 | 395 | 429 | 1215 |
| 65 | Rh_frontalpole | 0.76 | 0.55 | 391 | 395 | 429 | 1215 |
| 66 | Rh_temporalpole | 0.39 | 0.82 | 391 | 395 | 429 | 1215 |
| 67 | Rh_transversetemporal | 1.75 | 0.14 | 391 | 395 | 429 | 1215 |
| 68 | Rh_insula | 1.38 | 0.24 | 391 | 395 | 429 | 1215 |
| 69 | Lh_thalamus | 3.68 | 0.01 | 391 | 395 | 429 | 1215 |
| 70 | Lh_caudate | 0.53 | 0.71 | 391 | 395 | 429 | 1215 |
| 71 | Lh_putamen | 3.11 | 0.01 | 391 | 395 | 429 | 1215 |
| 72 | Lh_pallidum | 0.67 | 0.61 | 391 | 395 | 429 | 1215 |
| 73 | brain stem | 6.34 | <0.001 | 391 | 395 | 429 | 1215 |
| 74 | Lh_hippocampus | 0.41 | 0.8 | 391 | 395 | 429 | 1215 |
| 75 | Lh_amygdala | 3.2 | 0.01 | 391 | 395 | 429 | 1215 |
| 76 | Lh_accumbens | 0.92 | 0.45 | 391 | 395 | 429 | 1215 |
| 77 | Lh_ventraldc | 1.08 | 0.37 | 391 | 395 | 429 | 1215 |
| 78 | Rh_thalamus | 5.27 | <0.001 | 391 | 395 | 429 | 1215 |
| 79 | Rh_caudate | 0.27 | 0.9 | 391 | 395 | 429 | 1215 |
| 80 | Rh_putamen | 2.44 | 0.04 | 391 | 395 | 429 | 1215 |
| 81 | Rh_pallidum | 1.19 | 0.32 | 391 | 395 | 429 | 1215 |
| 82 | Rh_hippocampus | 1.94 | 0.1 | 391 | 395 | 429 | 1215 |
| 83 | Rh_amygdala | 0.94 | 0.44 | 391 | 395 | 429 | 1215 |
| 84 | Rh_accumbens | 1.82 | 0.12 | 391 | 395 | 429 | 1215 |
| 85 | Rh_ventraldc | 0.97 | 0.42 | 391 | 395 | 429 | 1215 |
| 86 | AUN-AUN | 6.89 | <0.001 | 391 | 395 | 428 | 1214 |
| 87 | AUN-CON | 12.13 | <0.001 | 391 | 394 | 427 | 1212 |
| 88 | AUN-CPN | 2.46 | 0.04 | 391 | 395 | 428 | 1214 |
| 89 | AUN-DMN | 3.45 | 0.01 | 391 | 394 | 427 | 1212 |
| 90 | AUN-DAN | 2.78 | 0.03 | 391 | 395 | 427 | 1213 |
| 91 | AUN-FPN | 10.16 | <0.001 | 391 | 394 | 428 | 1213 |
| 92 | AUN-RTN | 6.64 | <0.001 | 391 | 394 | 428 | 1213 |
| 93 | AUN-SAN | 26.17 | <0.001 | 391 | 395 | 428 | 1214 |
| 94 | AUN-SHN | 10.95 | <0.001 | 389 | 394 | 426 | 1209 |
| 95 | AUN-SMN | 3.18 | 0.01 | 391 | 395 | 428 | 1214 |
| 96 | AUN-VAN | 14.01 | <0.001 | 391 | 395 | 427 | 1213 |
| 97 | AUN-VIN | 14.63 | <0.001 | 391 | 394 | 428 | 1213 |
| 98 | CON-CON | 5.03 | <0.001 | 391 | 394 | 427 | 1212 |
| 99 | CON-CPN | 0.94 | 0.44 | 391 | 394 | 427 | 1212 |
| 100 | CON-DMN | 3 | 0.02 | 391 | 393 | 427 | 1211 |
| 101 | CON-DAN | 1.94 | 0.1 | 391 | 394 | 427 | 1212 |
| 102 | CON-FPN | 2.89 | 0.02 | 391 | 393 | 427 | 1211 |
| 103 | CON-RTN | 7.24 | <0.001 | 391 | 393 | 427 | 1211 |
| 104 | CON-SAN | 9.36 | <0.001 | 391 | 394 | 427 | 1212 |
| 105 | CON-SHN | 1.35 | 0.25 | 389 | 394 | 426 | 1209 |
| 106 | CON-SMN | 3.39 | 0.01 | 391 | 394 | 427 | 1212 |
| 107 | CON-VAN | 6.61 | <0.001 | 391 | 394 | 427 | 1212 |
| 108 | CON-VIN | 4.79 | <0.001 | 391 | 393 | 427 | 1211 |
| 109 | CPN-CPN | 1.83 | 0.12 | 391 | 395 | 429 | 1215 |
| 110 | CPN-DMN | 0.95 | 0.44 | 391 | 394 | 427 | 1212 |
| 111 | CPN-DAN | 0.99 | 0.41 | 391 | 395 | 427 | 1213 |
| 112 | CPN-FPN | 4.5 | <0.001 | 391 | 394 | 428 | 1213 |
| 113 | CPN-RTN | 2.87 | 0.02 | 391 | 394 | 428 | 1213 |
| 114 | CPN-SAN | 4.46 | <0.001 | 391 | 395 | 429 | 1215 |
| 115 | CPN-SHN | 6.29 | <0.001 | 389 | 394 | 427 | 1210 |
| 116 | CPN-SMN | 2.74 | 0.03 | 391 | 395 | 429 | 1215 |
| 117 | CPN-VAN | 0.86 | 0.49 | 391 | 395 | 427 | 1213 |
| 118 | CPN-VIN | 1.3 | 0.27 | 391 | 394 | 428 | 1213 |
| 119 | DMN-DMN | 4.17 | <0.001 | 391 | 394 | 427 | 1212 |
| 120 | DMN-DAN | 3.05 | 0.02 | 391 | 394 | 427 | 1212 |
| 121 | DMN-FPN | 3.39 | 0.01 | 391 | 394 | 427 | 1212 |
| 122 | DMN-RTN | 5.86 | <0.001 | 391 | 394 | 427 | 1212 |
| 123 | DMN-SAN | 3.05 | 0.02 | 391 | 394 | 427 | 1212 |
| 124 | DMN-SHN | 4.87 | <0.001 | 389 | 393 | 426 | 1208 |
| 125 | DMN-SMN | 0.31 | 0.87 | 391 | 394 | 427 | 1212 |
| 126 | DMN-VAN | 1.26 | 0.28 | 391 | 394 | 427 | 1212 |
| 127 | DMN-VIN | 3.6 | 0.01 | 391 | 394 | 427 | 1212 |
| 128 | DAN-DAN | 2.87 | 0.02 | 391 | 395 | 427 | 1213 |
| 129 | DAN-FPN | 1.42 | 0.22 | 391 | 394 | 427 | 1212 |
| 130 | DAN-RTN | 1.44 | 0.22 | 391 | 394 | 427 | 1212 |
| 131 | DAN-SAN | 0.26 | 0.91 | 391 | 395 | 427 | 1213 |
| 132 | DAN-SHN | 0.46 | 0.76 | 389 | 394 | 426 | 1209 |
| 133 | DAN-SMN | 1.15 | 0.33 | 391 | 395 | 427 | 1213 |
| 134 | DAN-VAN | 0.71 | 0.59 | 391 | 395 | 427 | 1213 |
| 135 | DAN-VIN | 1.19 | 0.31 | 391 | 394 | 427 | 1212 |
| 136 | FPN-FPN | 3.09 | 0.02 | 391 | 394 | 428 | 1213 |
| 137 | FPN-RTN | 1.18 | 0.32 | 391 | 394 | 428 | 1213 |
| 138 | FPN-SAN | 3.03 | 0.02 | 391 | 394 | 428 | 1213 |
| 139 | FPN-SHN | 13.82 | <0.001 | 389 | 393 | 426 | 1208 |
| 140 | FPN-SMN | 6.23 | <0.001 | 391 | 394 | 428 | 1213 |
| 141 | FPN-VAN | 2.18 | 0.07 | 391 | 394 | 427 | 1212 |
| 142 | FPN-VIN | 3.7 | 0.01 | 391 | 394 | 428 | 1213 |
| 143 | RTN-RTN | 2.26 | 0.06 | 391 | 394 | 428 | 1213 |
| 144 | RTN-SAN | 5.73 | <0.001 | 391 | 394 | 428 | 1213 |
| 145 | RTN-SHN | 3.25 | 0.01 | 389 | 393 | 426 | 1208 |
| 146 | RTN-SMN | 6.17 | <0.001 | 391 | 394 | 428 | 1213 |
| 147 | RTN-VAN | 2.66 | 0.03 | 391 | 394 | 427 | 1212 |
| 148 | RTN-VIN | 1.63 | 0.16 | 391 | 394 | 428 | 1213 |
| 149 | SAN-SAN | 1.33 | 0.26 | 391 | 395 | 429 | 1215 |
| 150 | SAN-SHN | 14.12 | <0.001 | 389 | 394 | 427 | 1210 |
| 151 | SAN-SMN | 12.25 | <0.001 | 391 | 395 | 429 | 1215 |
| 152 | SAN-VAN | 4.91 | <0.001 | 391 | 395 | 427 | 1213 |
| 153 | SAN-VIN | 1.41 | 0.23 | 391 | 394 | 428 | 1213 |
| 154 | SHN-SHN | 17 | <0.001 | 389 | 394 | 427 | 1210 |
| 155 | SHN-SMN | 14.21 | <0.001 | 389 | 394 | 427 | 1210 |
| 156 | SHN-VAN | 0.75 | 0.56 | 389 | 394 | 426 | 1209 |
| 157 | SHN-VIN | 13.48 | <0.001 | 389 | 393 | 426 | 1208 |
| 158 | SMN-SMN | 2.25 | 0.06 | 391 | 395 | 429 | 1215 |
| 159 | SMN-VAN | 1.03 | 0.39 | 391 | 395 | 427 | 1213 |
| 160 | SMN-VIN | 4.28 | <0.001 | 391 | 394 | 428 | 1213 |
| 161 | VAN-VAN | 2.89 | 0.02 | 391 | 395 | 427 | 1213 |
| 162 | VAN-VIN | 5.01 | <0.001 | 391 | 394 | 427 | 1212 |
| 163 | VIN-VIN | 3.74 | <0.001 | 391 | 394 | 428 | 1213 |
| 164 | AUN-tha.Lh | 1.25 | 0.29 | 391 | 394 | 427 | 1212 |
| 165 | AUN-cau.Lh | 2.16 | 0.07 | 391 | 395 | 428 | 1214 |
| 166 | AUN-put.Lh | 5.58 | <0.001 | 391 | 394 | 427 | 1212 |
| 167 | AUN-pal.Lh | 1.98 | 0.09 | 391 | 395 | 427 | 1213 |
| 168 | AUN-hip.Lh | 4.3 | <0.001 | 390 | 393 | 426 | 1209 |
| 169 | AUN-amy.Lh | 1.43 | 0.22 | 391 | 394 | 428 | 1213 |
| 170 | AUN-acc.Lh | 8.3 | <0.001 | 389 | 394 | 426 | 1209 |
| 171 | AUN-vdc.Lh | 10.12 | <0.001 | 391 | 395 | 428 | 1214 |
| 172 | AUN-b.s. | 6.61 | <0.001 | 391 | 394 | 428 | 1213 |
| 173 | AUN-tha.Rh | 1.05 | 0.38 | 391 | 395 | 427 | 1213 |
| 174 | AUN-cau.Rh | 5.9 | <0.001 | 391 | 394 | 428 | 1213 |
| 175 | AUN-put.Rh | 22.99 | <0.001 | 391 | 395 | 428 | 1214 |
| 176 | AUN-pal.Rh | 6.21 | <0.001 | 391 | 394 | 427 | 1212 |
| 177 | AUN-hip.Rh | 6.05 | <0.001 | 391 | 395 | 429 | 1215 |
| 178 | AUN-amy.Rh | 2.88 | 0.02 | 391 | 394 | 427 | 1212 |
| 179 | AUN-acc.Rh | 0.71 | 0.59 | 391 | 395 | 427 | 1213 |
| 180 | AUN-vdc.Rh | 8.43 | <0.001 | 391 | 394 | 428 | 1213 |
| 181 | CON-tha.Lh | 0.95 | 0.43 | 391 | 394 | 428 | 1213 |
| 182 | CON-cau.Lh | 12.79 | <0.001 | 389 | 394 | 427 | 1210 |
| 183 | CON-put.Lh | 18.05 | <0.001 | 391 | 395 | 429 | 1215 |
| 184 | CON-pal.Lh | 5.08 | <0.001 | 391 | 395 | 429 | 1215 |
| 185 | CON-hip.Lh | 1.34 | 0.25 | 391 | 394 | 428 | 1213 |
| 186 | CON-amy.Lh | 33.26 | <0.001 | 391 | 395 | 428 | 1214 |
| 187 | CON-acc.Lh | 10.12 | <0.001 | 391 | 394 | 427 | 1212 |
| 188 | CON-vdc.Lh | 11.11 | <0.001 | 391 | 395 | 429 | 1215 |
| 189 | CON-b.s. | 7.94 | <0.001 | 391 | 395 | 427 | 1213 |
| 190 | CON-tha.Rh | 1.3 | 0.27 | 391 | 395 | 427 | 1213 |
| 191 | CON-cau.Rh | 3.57 | 0.01 | 391 | 394 | 428 | 1213 |
| 192 | CON-put.Rh | 2.87 | 0.02 | 390 | 393 | 426 | 1209 |
| 193 | CON-pal.Rh | 2.13 | 0.07 | 391 | 394 | 428 | 1213 |
| 194 | CON-hip.Rh | 22.71 | <0.001 | 389 | 394 | 427 | 1210 |
| 195 | CON-amy.Rh | 18.98 | <0.001 | 391 | 395 | 429 | 1215 |
| 196 | CON-acc.Rh | 1.15 | 0.33 | 391 | 395 | 429 | 1215 |
| 197 | CON-vdc.Rh | 4.15 | <0.001 | 391 | 395 | 427 | 1213 |
| 198 | CPN-tha.Lh | 19.86 | <0.001 | 391 | 395 | 428 | 1214 |
| 199 | CPN-cau.Lh | 4.34 | <0.001 | 391 | 394 | 427 | 1212 |
| 200 | CPN-put.Lh | 5.83 | <0.001 | 391 | 395 | 429 | 1215 |
| 201 | CPN-pal.Lh | 1.69 | 0.15 | 391 | 394 | 427 | 1212 |
| 202 | CPN-hip.Lh | 0.76 | 0.55 | 391 | 394 | 428 | 1213 |
| 203 | CPN-amy.Lh | 3.03 | 0.02 | 390 | 393 | 426 | 1209 |
| 204 | CPN-acc.Lh | 3.64 | 0.01 | 391 | 394 | 428 | 1213 |
| 205 | CPN-vdc.Lh | 8.85 | <0.001 | 389 | 394 | 427 | 1210 |
| 206 | CPN-b.s. | 1.4 | 0.23 | 391 | 395 | 427 | 1213 |
| 207 | CPN-tha.Rh | 5.43 | <0.001 | 391 | 395 | 429 | 1215 |
| 208 | CPN-cau.Rh | 9.52 | <0.001 | 391 | 395 | 427 | 1213 |
| 209 | CPN-put.Rh | 2.62 | 0.03 | 391 | 394 | 428 | 1213 |
| 210 | CPN-pal.Rh | 17.35 | <0.001 | 391 | 395 | 428 | 1214 |
| 211 | CPN-hip.Rh | 7.4 | <0.001 | 391 | 394 | 427 | 1212 |
| 212 | CPN-amy.Rh | 2.91 | 0.02 | 391 | 395 | 429 | 1215 |
| 213 | CPN-acc.Rh | 3.18 | 0.01 | 391 | 394 | 427 | 1212 |
| 214 | CPN-vdc.Rh | 0.33 | 0.86 | 391 | 395 | 427 | 1213 |
| 215 | DMN-tha.Lh | 2.06 | 0.08 | 390 | 393 | 426 | 1209 |
| 216 | DMN-cau.Lh | 2.85 | 0.02 | 391 | 394 | 428 | 1213 |
| 217 | DMN-put.Lh | 11.35 | <0.001 | 389 | 394 | 427 | 1210 |
| 218 | DMN-pal.Lh | 12.32 | <0.001 | 391 | 395 | 429 | 1215 |
| 219 | DMN-hip.Lh | 2.88 | 0.02 | 391 | 395 | 427 | 1213 |
| 220 | DMN-amy.Lh | 1.55 | 0.19 | 391 | 394 | 428 | 1213 |
| 221 | DMN-acc.Lh | 6.58 | <0.001 | 391 | 395 | 428 | 1214 |
| 222 | DMN-vdc.Lh | 1.46 | 0.21 | 391 | 394 | 427 | 1212 |
| 223 | DMN-b.s. | 1.25 | 0.29 | 391 | 395 | 429 | 1215 |
| 224 | DMN-tha.Rh | 1.16 | 0.33 | 391 | 394 | 427 | 1212 |
| 225 | DMN-cau.Rh | 0.6 | 0.66 | 391 | 395 | 427 | 1213 |
| 226 | DMN-put.Rh | 4.08 | <0.001 | 391 | 394 | 428 | 1213 |
| 227 | DMN-pal.Rh | 2.77 | 0.03 | 390 | 393 | 426 | 1209 |
| 228 | DMN-hip.Rh | 2.46 | 0.04 | 391 | 394 | 428 | 1213 |
| 229 | DMN-amy.Rh | 4.83 | <0.001 | 389 | 394 | 426 | 1209 |
| 230 | DMN-acc.Rh | 6.42 | <0.001 | 391 | 395 | 428 | 1214 |
| 231 | DMN-vdc.Rh | 3.13 | 0.01 | 391 | 395 | 428 | 1214 |
| 232 | DAN-tha.Lh | 0.32 | 0.87 | 391 | 394 | 428 | 1213 |
| 233 | DAN-cau.Lh | 1.38 | 0.24 | 391 | 395 | 428 | 1214 |
| 234 | DAN-put.Lh | 4.16 | <0.001 | 391 | 394 | 427 | 1212 |
| 235 | DAN-pal.Lh | 1.17 | 0.32 | 391 | 395 | 428 | 1214 |
| 236 | DAN-hip.Lh | 3.37 | 0.01 | 391 | 395 | 427 | 1213 |
| 237 | DAN-amy.Lh | 0.39 | 0.82 | 391 | 394 | 428 | 1213 |
| 238 | DAN-acc.Lh | 4.55 | <0.001 | 390 | 393 | 426 | 1209 |
| 239 | DAN-vdc.Lh | 0.68 | 0.6 | 391 | 394 | 428 | 1213 |
| 240 | DAN-b.s. | 7.12 | <0.001 | 391 | 394 | 427 | 1212 |
| 241 | DAN-tha.Rh | 3.11 | 0.01 | 391 | 395 | 428 | 1214 |
| 242 | DAN-cau.Rh | 4.21 | <0.001 | 391 | 395 | 428 | 1214 |
| 243 | DAN-put.Rh | 3.85 | <0.001 | 391 | 395 | 427 | 1213 |
| 244 | DAN-pal.Rh | 3.94 | <0.001 | 391 | 394 | 428 | 1213 |
| 245 | DAN-hip.Rh | 0.73 | 0.57 | 391 | 394 | 428 | 1213 |
| 246 | DAN-amy.Rh | 1 | 0.41 | 391 | 393 | 427 | 1211 |
| 247 | DAN-acc.Rh | 1.61 | 0.17 | 391 | 394 | 428 | 1213 |
| 248 | DAN-vdc.Rh | 4.33 | <0.001 | 391 | 394 | 427 | 1212 |
| 249 | FPN-tha.Lh | 0.78 | 0.54 | 391 | 394 | 428 | 1213 |
| 250 | FPN-cau.Lh | 4.96 | <0.001 | 390 | 393 | 426 | 1209 |
| 251 | FPN-put.Lh | 1.52 | 0.19 | 391 | 394 | 428 | 1213 |
| 252 | FPN-pal.Lh | 6.29 | <0.001 | 389 | 393 | 426 | 1208 |
| 253 | FPN-hip.Lh | 4.43 | <0.001 | 391 | 394 | 428 | 1213 |
| 254 | FPN-amy.Lh | 4.11 | <0.001 | 391 | 394 | 427 | 1212 |
| 255 | FPN-acc.Lh | 1.93 | 0.1 | 391 | 394 | 428 | 1213 |
| 256 | FPN-vdc.Lh | 3.79 | <0.001 | 391 | 395 | 428 | 1214 |
| 257 | FPN-b.s. | 1.96 | 0.1 | 391 | 394 | 428 | 1213 |
| 258 | FPN-tha.Rh | 0.44 | 0.78 | 391 | 395 | 429 | 1215 |
| 259 | FPN-cau.Rh | 0.72 | 0.57 | 391 | 394 | 427 | 1212 |
| 260 | FPN-put.Rh | 0.85 | 0.49 | 391 | 395 | 427 | 1213 |
| 261 | FPN-pal.Rh | 1.13 | 0.34 | 391 | 394 | 428 | 1213 |
| 262 | FPN-hip.Rh | 2.4 | 0.05 | 390 | 393 | 426 | 1209 |
| 263 | FPN-amy.Rh | 0.21 | 0.93 | 391 | 394 | 428 | 1213 |
| 264 | FPN-acc.Rh | 3.41 | 0.01 | 389 | 394 | 427 | 1210 |
| 265 | FPN-vdc.Rh | 1.71 | 0.14 | 391 | 395 | 429 | 1215 |
| 266 | RTN-tha.Lh | 0.15 | 0.96 | 391 | 395 | 427 | 1213 |
| 267 | RTN-cau.Lh | 3.15 | 0.01 | 391 | 394 | 428 | 1213 |
| 268 | RTN-put.Lh | 1.54 | 0.19 | 390 | 393 | 426 | 1209 |
| 269 | RTN-pal.Lh | 0.91 | 0.46 | 391 | 394 | 428 | 1213 |
| 270 | RTN-hip.Lh | 8.49 | <0.001 | 391 | 395 | 428 | 1214 |
| 271 | RTN-amy.Lh | 5.54 | <0.001 | 391 | 395 | 428 | 1214 |
| 272 | RTN-acc.Lh | 3.92 | <0.001 | 391 | 395 | 427 | 1213 |
| 273 | RTN-vdc.Lh | 2.58 | 0.04 | 391 | 394 | 428 | 1213 |
| 274 | RTN-b.s. | 6.86 | <0.001 | 389 | 394 | 426 | 1209 |
| 275 | RTN-tha.Rh | 5.04 | <0.001 | 391 | 394 | 427 | 1212 |
| 276 | RTN-cau.Rh | 6.07 | <0.001 | 391 | 395 | 429 | 1215 |
| 277 | RTN-put.Rh | 0.67 | 0.61 | 391 | 394 | 427 | 1212 |
| 278 | RTN-pal.Rh | 1.72 | 0.14 | 391 | 395 | 427 | 1213 |
| 279 | RTN-hip.Rh | 8.41 | <0.001 | 391 | 394 | 428 | 1213 |
| 280 | RTN-amy.Rh | 0.97 | 0.42 | 390 | 393 | 426 | 1209 |
| 281 | RTN-acc.Rh | 0.99 | 0.41 | 391 | 394 | 428 | 1213 |
| 282 | RTN-vdc.Rh | 9.39 | <0.001 | 389 | 394 | 427 | 1210 |
| 283 | SAN-tha.Lh | 6.94 | <0.001 | 391 | 395 | 429 | 1215 |
| 284 | SAN-cau.Lh | 4.13 | <0.001 | 391 | 395 | 429 | 1215 |
| 285 | SAN-put.Lh | 2.47 | 0.04 | 391 | 395 | 427 | 1213 |
| 286 | SAN-pal.Lh | 3.67 | 0.01 | 391 | 394 | 428 | 1213 |
| 287 | SAN-hip.Lh | 4.6 | <0.001 | 391 | 394 | 427 | 1212 |
| 288 | SAN-amy.Lh | 2.56 | 0.04 | 391 | 395 | 428 | 1214 |
| 289 | SAN-acc.Lh | 9.76 | <0.001 | 391 | 394 | 427 | 1212 |
| 290 | SAN-vdc.Lh | 3.18 | 0.01 | 391 | 395 | 427 | 1213 |
| 291 | SAN-b.s. | 4.46 | <0.001 | 391 | 395 | 428 | 1214 |
| 292 | SAN-tha.Rh | 3.12 | 0.01 | 390 | 393 | 426 | 1209 |
| 293 | SAN-cau.Rh | 0.99 | 0.41 | 391 | 394 | 428 | 1213 |
| 294 | SAN-put.Rh | 1.69 | 0.15 | 389 | 394 | 426 | 1209 |
| 295 | SAN-pal.Rh | 5.43 | <0.001 | 391 | 395 | 428 | 1214 |
| 296 | SAN-hip.Rh | 3.43 | 0.01 | 391 | 395 | 428 | 1214 |
| 297 | SAN-amy.Rh | 2.66 | 0.03 | 391 | 395 | 427 | 1213 |
| 298 | SAN-acc.Rh | 2.78 | 0.03 | 391 | 394 | 428 | 1213 |
| 299 | SAN-vdc.Rh | 2.13 | 0.07 | 391 | 394 | 428 | 1213 |
| 300 | SHN-tha.Lh | 3.2 | 0.01 | 391 | 395 | 429 | 1215 |
| 301 | SHN-cau.Lh | 7.2 | <0.001 | 391 | 395 | 427 | 1213 |
| 302 | SHN-put.Lh | 2.11 | 0.08 | 391 | 394 | 428 | 1213 |
| 303 | SHN-pal.Lh | 32.04 | <0.001 | 391 | 395 | 428 | 1214 |
| 304 | SHN-hip.Lh | 9.66 | <0.001 | 391 | 395 | 429 | 1215 |
| 305 | SHN-amy.Lh | 1.53 | 0.19 | 391 | 394 | 427 | 1212 |
| 306 | SHN-acc.Lh | 0.7 | 0.59 | 391 | 395 | 427 | 1213 |
| 307 | SHN-vdc.Lh | 4.16 | <0.001 | 391 | 394 | 428 | 1213 |
| 308 | SHN-b.s. | 12.31 | <0.001 | 391 | 394 | 427 | 1212 |
| 309 | SHN-tha.Rh | 1 | 0.41 | 391 | 394 | 428 | 1213 |
| 310 | SHN-cau.Rh | 24.76 | <0.001 | 389 | 394 | 427 | 1210 |
| 311 | SHN-put.Rh | 20.08 | <0.001 | 391 | 395 | 429 | 1215 |
| 312 | SHN-pal.Rh | 1.05 | 0.38 | 391 | 395 | 429 | 1215 |
| 313 | SHN-hip.Rh | 6.78 | <0.001 | 391 | 395 | 427 | 1213 |
| 314 | SHN-amy.Rh | 0.44 | 0.78 | 391 | 394 | 428 | 1213 |
| 315 | SHN-acc.Rh | 18.76 | <0.001 | 391 | 395 | 428 | 1214 |
| 316 | SHN-vdc.Rh | 4.74 | <0.001 | 391 | 394 | 427 | 1212 |
| 317 | SMN-tha.Lh | 2.55 | 0.04 | 391 | 394 | 427 | 1212 |
| 318 | SMN-cau.Lh | 2.52 | 0.04 | 391 | 395 | 427 | 1213 |
| 319 | SMN-put.Lh | 0.66 | 0.62 | 391 | 394 | 428 | 1213 |
| 320 | SMN-pal.Lh | 2.73 | 0.03 | 390 | 393 | 426 | 1209 |
| 321 | SMN-hip.Lh | 9.43 | <0.001 | 389 | 394 | 427 | 1210 |
| 322 | SMN-amy.Lh | 10.28 | <0.001 | 391 | 395 | 429 | 1215 |
| 323 | SMN-acc.Lh | 8.22 | <0.001 | 391 | 395 | 429 | 1215 |
| 324 | SMN-vdc.Lh | 11.38 | <0.001 | 391 | 395 | 427 | 1213 |
| 325 | SMN-b.s. | 3.55 | 0.01 | 391 | 394 | 428 | 1213 |
| 326 | SMN-tha.Rh | 10.82 | <0.001 | 391 | 395 | 428 | 1214 |
| 327 | SMN-cau.Rh | 4.55 | <0.001 | 391 | 394 | 427 | 1212 |
| 328 | SMN-put.Rh | 1.84 | 0.12 | 391 | 395 | 429 | 1215 |
| 329 | SMN-pal.Rh | 4.71 | <0.001 | 391 | 394 | 427 | 1212 |
| 330 | SMN-hip.Rh | 2.05 | 0.08 | 391 | 395 | 427 | 1213 |
| 331 | SMN-amy.Rh | 0.71 | 0.58 | 391 | 394 | 428 | 1213 |
| 332 | SMN-acc.Rh | 0.79 | 0.53 | 390 | 393 | 426 | 1209 |
| 333 | SMN-vdc.Rh | 1.06 | 0.38 | 391 | 394 | 428 | 1213 |
| 334 | VAN-tha.Lh | 1.09 | 0.36 | 391 | 394 | 428 | 1213 |
| 335 | VAN-cau.Lh | 3.78 | <0.001 | 391 | 394 | 427 | 1212 |
| 336 | VAN-put.Lh | 2.12 | 0.08 | 391 | 394 | 427 | 1212 |
| 337 | VAN-pal.Lh | 0.27 | 0.9 | 391 | 394 | 428 | 1213 |
| 338 | VAN-hip.Lh | 2.68 | 0.03 | 391 | 394 | 428 | 1213 |
| 339 | VAN-amy.Lh | 3.41 | 0.01 | 389 | 393 | 426 | 1208 |
| 340 | VAN-acc.Lh | 5.13 | <0.001 | 391 | 394 | 428 | 1213 |
| 341 | VAN-vdc.Lh | 2.57 | 0.04 | 391 | 394 | 428 | 1213 |
| 342 | VAN-b.s. | 4.59 | <0.001 | 390 | 393 | 426 | 1209 |
| 343 | VAN-tha.Rh | 0.48 | 0.75 | 391 | 394 | 428 | 1213 |
| 344 | VAN-cau.Rh | 4.17 | <0.001 | 391 | 395 | 428 | 1214 |
| 345 | VAN-put.Rh | 0.93 | 0.45 | 391 | 394 | 427 | 1212 |
| 346 | VAN-pal.Rh | 0.46 | 0.77 | 391 | 395 | 429 | 1215 |
| 347 | VAN-hip.Rh | 1.63 | 0.17 | 391 | 394 | 427 | 1212 |
| 348 | VAN-amy.Rh | 1.63 | 0.16 | 391 | 395 | 427 | 1213 |
| 349 | VAN-acc.Rh | 0.64 | 0.64 | 391 | 394 | 428 | 1213 |
| 350 | VAN-vdc.Rh | 3.12 | 0.01 | 390 | 393 | 426 | 1209 |
| 351 | VIN-tha.Lh | 4.92 | <0.001 | 389 | 394 | 427 | 1210 |
| 352 | VIN-cau.Lh | 2.49 | 0.04 | 391 | 395 | 429 | 1215 |
| 353 | VIN-put.Lh | 0.26 | 0.91 | 391 | 395 | 429 | 1215 |
| 354 | VIN-pal.Lh | 1.06 | 0.38 | 391 | 395 | 427 | 1213 |
| 355 | VIN-hip.Lh | 5.69 | <0.001 | 391 | 395 | 428 | 1214 |
| 356 | VIN-amy.Lh | 2.64 | 0.03 | 391 | 394 | 427 | 1212 |
| 357 | VIN-acc.Lh | 3.96 | <0.001 | 391 | 395 | 428 | 1214 |
| 358 | VIN-vdc.Lh | 0.78 | 0.54 | 391 | 394 | 427 | 1212 |
| 359 | VIN-b.s. | 1.35 | 0.25 | 391 | 394 | 428 | 1213 |
| 360 | VIN-tha.Rh | 4.05 | <0.001 | 391 | 394 | 428 | 1213 |
| 361 | VIN-cau.Rh | 0.94 | 0.44 | 390 | 393 | 426 | 1209 |
| 362 | VIN-put.Rh | 1.27 | 0.28 | 391 | 394 | 428 | 1213 |
| 363 | VIN-pal.Rh | 5.99 | <0.001 | 389 | 394 | 426 | 1209 |
| 364 | VIN-hip.Rh | 6.65 | <0.001 | 391 | 395 | 428 | 1214 |
| 365 | VIN-amy.Rh | 4.01 | <0.001 | 391 | 395 | 428 | 1214 |
| 366 | VIN-acc.Rh | 1.09 | 0.36 | 391 | 395 | 427 | 1213 |
| 367 | VIN-vdc.Rh | 1.85 | 0.12 | 391 | 394 | 428 | 1213 |
